# Mendelian randomisation for mediation analysis: current methods and challenges for implementation

**DOI:** 10.1101/835819

**Authors:** Alice R Carter, Eleanor Sanderson, Gemma Hammerton, Rebecca C Richmond, George Davey Smith, Jon Heron, Amy E Taylor, Neil M Davies, Laura D Howe

## Abstract

Mediation analysis seeks to explain the pathway(s) through which an exposure affects an outcome. Mediation analysis experiences a number of methodological difficulties, including bias due to confounding and measurement error. Mendelian randomisation (MR) can be used to improve causal inference for mediation analysis. We describe two approaches that can be used for estimating mediation analysis with MR: multivariable Mendelian randomisation (MVMR) and two-step Mendelian randomisation. We outline the approaches and provide code to demonstrate how they can be used in mediation analysis. We review issues that can affect analyses, including confounding, measurement error, weak instrument bias, and analysis of multiple mediators. Description of the methods is supplemented by simulated and real data examples. Although Mendelian randomisation relies on large sample sizes and strong assumptions, such as having strong instruments and no horizontally pleiotropic pathways, our examples demonstrate that it is unlikely to be affected by confounders of the exposure or mediator and the outcome, reverse causality and non-differential measurement error of the exposure or mediator. Both MVMR and two-step MR can be implemented in both individual-level MR and summary data MR, and can improve causal inference in mediation analysis.

## Introduction

Mediation analysis can improve aetiological understanding, and identify intermediate variables as potential intervention targets when intervening on an exposure is not feasible. However, in order to make causal inferences, phenotypic mediation analysis requires strong assumptions. Mendelian randomisation (MR) is an alternative causal inference approach using genetic variants as instrumental variables for a phenotype [1]. In this paper we compare phenotypic regression-based methods for mediation analysis with MR methods for mediation analysis, and describe the assumptions required for MR mediation methods to make valid causal inferences.

### Mediation Analysis

Methods for mediation analysis emerged in the early twentieth-century, although often not described as such at the time, with formal methods developed by Baron and Kenny in the 1980s [2, 3]. More recently, a large amount of research has built on and improved mediation methods for better causal inference [4].

Three parameters are typically estimated in traditional mediation analysis i) the total effect (the effect of the exposure on the outcome through all potential pathways) ii) the direct effect (the remaining effect of the exposure on the outcome that acts through pathways other than the specified mediator or set of mediators) and iii) the indirect effect (the path from exposure to outcome that acts through the mediator(s)). In situations where the total effect, direct effect and indirect effect all act in the same direction, an estimate of the “proportion mediated” (i.e. proportion of the total effect explained by the mediator) can be calculated. Two common approaches to estimate the indirect effect are; the product of coefficients method and the difference in coefficients method [5] (see Figure 1A).

**Figure 1:**
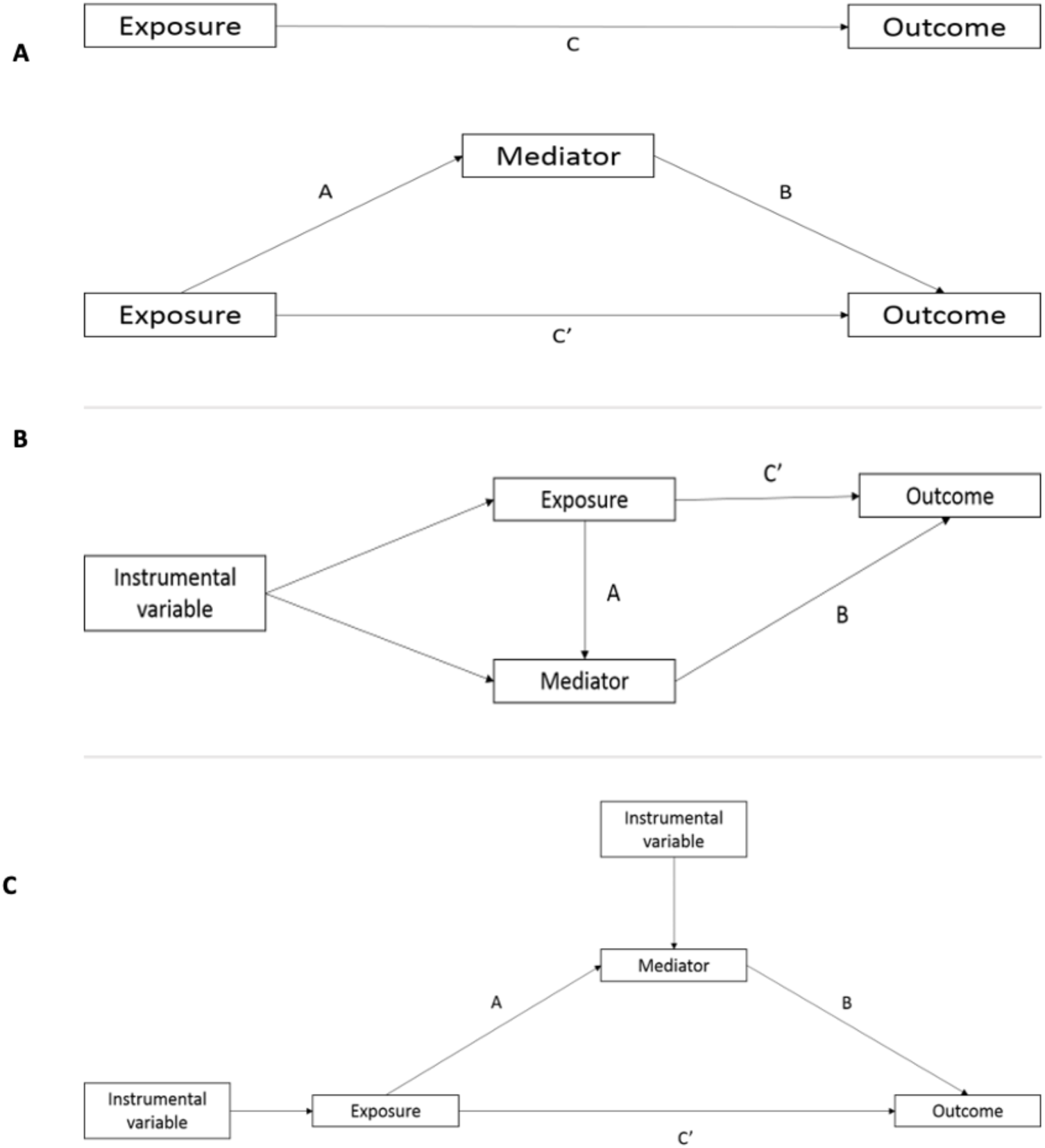
The decomposed effects in A) phenotypic regression-based mediation analysis where C represents the total effect, C’ represe nts the direct effect and the indirect effect can be calculated by subtracting C’ from C (difference method) or multiplying A times B (product of coefficients method) B) multivariable MR, using a combined genetic instrument for both the exposure and mediator of interest, to estimate the direct effect (C’) of the exposure and C) two-step Mendelian randomisation, where the effect of the exposure on the mediator (A) and mediator on the outcome (B) are estimated separately, using separate genetic instrumental variables for both the exposure and mediator. These estimates are then multiplied together to estimate the indirect effect of the mediator (A*B)

Traditional mediation methods, such as Baron and Kenny methods, rely on several strong, untestable assumptions including, among others i) a causal effect of the exposure on the outcome, exposure on the mediator and mediator on the outcome ii) no unmeasured confounding between the exposure, mediator and outcome iii) no exposure-caused confounders of the mediator and outcome (intermediate confounders, see Figure 2A) and iv) no exposure-mediator interaction [4, 6, 7]. Furthermore, measurement error in either the exposure or mediator can introduce bias [8].

**Figure 2:**
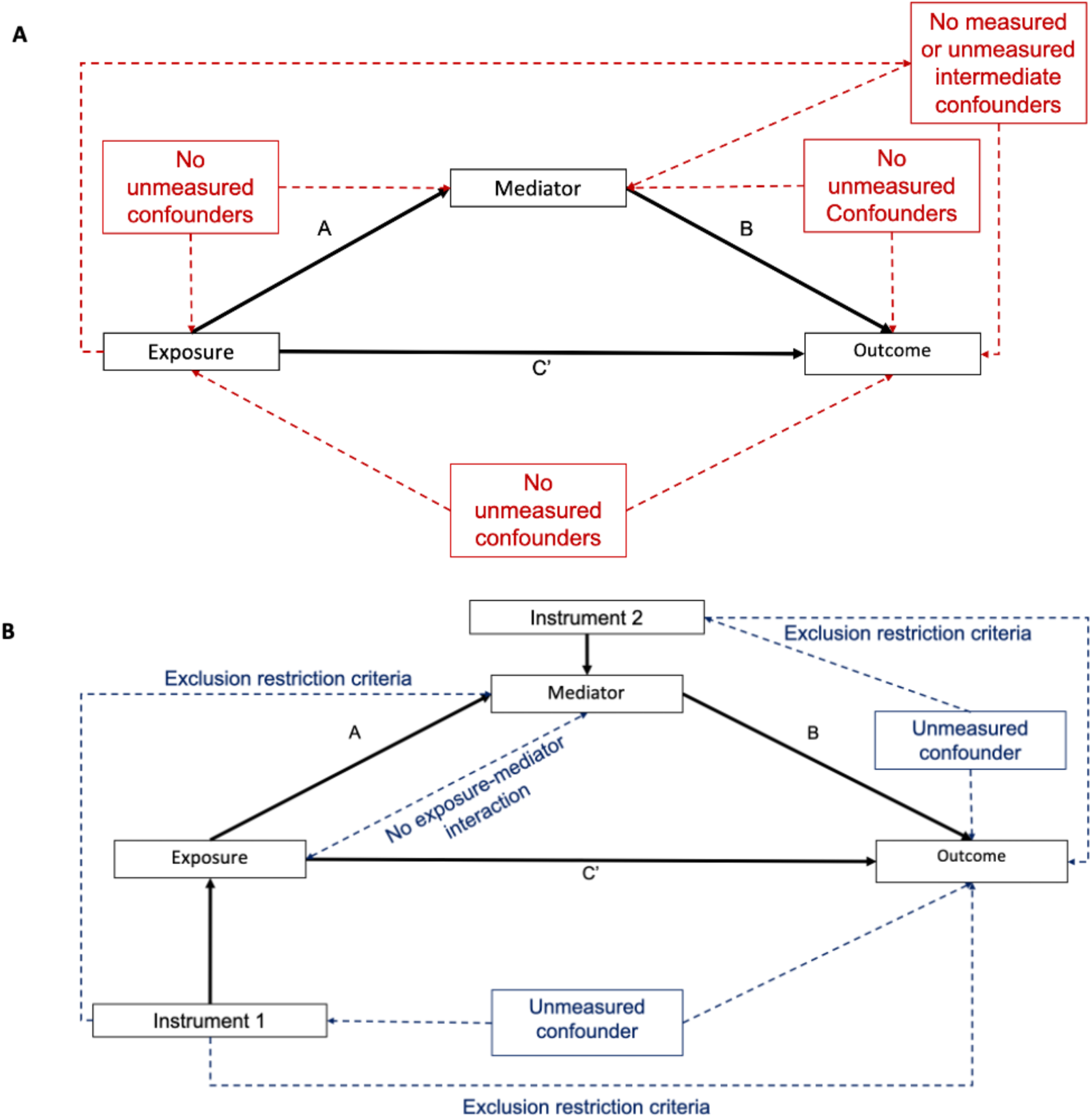
Schematic diagram illustrating the causal assumptions (dashed lines) in A) phenotypic regression-based mediation methods and B) Mendelian randomisation mediation analysis with the measured associations in solid black lines. Additional assumption for phenotypic mediation is that of no measurement error in the exposure or mediator In Mendelian randomisation, the exclusion restriction criteria means there are no alternative pathways from the instrument to the outcome other than via the exposure (or mediator) of interest.

Baron and Kenny methods were introduced to estimate mediation with a continuous exposure, outcome and mediator, although they are also now often applied to binary variables. In the presence of a continuous or rare binary outcome the estimates from the difference in coefficients and the product of coefficients method should coincide [4, 9].

Counterfactual reasoning has been used to develop methods that can address some of the previously described strong assumptions [10-14]. These methods can estimate mediation in the presence of exposure-mediator interactions and account for measured intermediate confounders. Additionally, these more flexible counterfactual methods can allow for binary mediators and outcomes. However, these methods remain biased in the presence of unmeasured confounding, measurement error in the exposure or mediator, or in a mis-specified model with reverse causality [4, 15]. In counterfactual methods, the estimated direct effect is described as being a “controlled direct effect” if the value of the mediator is controlled at a certain value for all individuals in the population, or a “natural direct effect”, when the value of the mediator is allowed to take the value for each person that it would have taken naturally had they been unexposed, in a counterfactual scenario. The “natural indirect effect” represents the average change in an outcome if the value of the exposure was fixed, but the value of the mediator changes from its natural value when exposed to its natural value when unexposed. If there is no interaction between the exposure and mediator, the estimate of the natural direct effect is equivalent to the controlled direct effect, and indeed would align with estimates from Baron and Kenny approaches to mediation [4, 9, 16].

### Mendelian randomisation

In Mendelian randomisation (MR) randomly allocated genetic variants are used as instrumental variables for a phenotype [1, 17, 18]. Given the random allocation of genetic variants at conception, MR estimates are robust to bias from confounding, reverse causation and measurement error [17]. Three core assumptions are required for a genetic variant to be a valid instrumental variable, these are i) the genetic variants are robustly associated with the exposure (the relevance assumption) ii) the genetic instruments are exchangeable with the outcome (the independence assumption) and iii) the genetic variants do not affect the outcome via any variable other than the exposure (the exclusion restriction criteria) (Online Resource 1: sFigure 1) [1].

### Rationale for using Mendelian randomisation in mediation analysis

Mendelian randomisation can be used to overcome some of the previously described strong assumptions required for causal inference in mediation analysis. For example, estimates are robust to bias from unmeasured confounding, including that of intermediate confounding, and estimates cannot be biased by reverse causality.

In mediation terms, a univariable MR estimates the total effect of the exposure on the outcome. Two differing MR approaches can then be used which broadly mirror traditional phenotypic regression-based approaches to mediation to decompose the direct and indirect effects: multivariable MR (MVMR) [19, 20] and two-step MR [21-23].

In MVMR the controlled direct effect of the exposure on the outcome, controlling for the mediator, is estimated [23, 19]. The genetic instrument for both the primary exposure and the second exposure (mediator) are included as instruments in the analysis (Figure 1B) [24, 25]. The indirect effect can then be estimated by subtracting the direct effect from the total effect (akin to the difference in coefficients method). MVMR assumes no interaction between the exposure and the mediator; therefore, the controlled direct effect estimated is equivalent to the natural direct effect where this assumption holds true. As such, we refer to this as the direct effect, without further distinction, throughout this manuscript.

Two-step Mendelian randomisation (also known as network MR) is akin to the product of coefficient methods. Two MR estimates are calculated i) the causal effect of the exposure on the mediator and ii) the causal effect of the mediator on the outcome (Figure 1C) [21, 26, 23]. These two estimates can then be multiplied together to estimate the indirect effect. Two-step MR also assumes no interaction between the exposure and the mediator.

These MR methods are increasingly being used in mediation analysis [19, 27-30]. In this paper, we demonstrate how MVMR and two-step MR can be used to estimate the direct effect, indirect effect and the proportion mediated, and which assumptions are required for the resulting estimates to be unbiased [25, 24, 23]. We provide guidance about how to carry out each method, with code provided, and illustrate each method using both simulated and real data (see Online Resource 2), applied to an individual level MR analysis.

## Methods

### Simulation study

We simulated data under the model illustrated in Figure 1 with continuous, rare binary (5% prevalence) and common binary (25% prevalence) outcomes. We varied the total effect of our exposure and proportion mediated and obtained results using phenotypic methods using both the difference and product of coefficients approaches, and MR methods using both MVMR and two-step MR. Additionally, we simulated results where the total effect of the exposure on the outcome is small, and where each of the exposure and mediator were subject to non-differential measurement error. Finally, we simulated how MR methods can estimate mediation in the presence of multiple mediators, these simulations are illustrated in Online Resource 1: sFigure2. The full range of scenarios simulated are presented in sTable 1. Simulation analyses were carried out using R version 3.5.1 and the corresponding code for the simulation studies can be found at https://github.com/eleanorsanderson/MediationMR.

### Applied example

Using data from UK Biobank (N = 195 218), we investigate the role of body mass index (BMI) and low-density lipoprotein cholesterol (LDL-C) in mediating the associations of education with systolic blood pressure, cardiovascular disease (CVD) and hypertension (continuous, rare binary and common binary outcomes, respectively). The effects on binary outcomes (hypertension and incident CVD) were estimated on risk difference, log odds ratio, and odds ratio scales. Applied analyses were performed using Stata version 15 (StataCorp LP, Texas) and corresponding code is available at https://github.com/alicerosecarter/MediationMR. The full worked through example is available in the Online Resource 2.

### Statistical analysis

The following approaches were applied to both applied analyses and simulated data. Equations describing each of these analyses are given in Online Resource 1.

#### Difference in coefficients method

Each outcome was regressed on the exposure adjusting for the mediator to estimate the direct effect of the exposure. The direct effect was subtracted from the total effect, estimated using multivariable regression adjusting for potential confounders, to estimate the indirect effect. In all simulation scenarios the standard deviation of the regression coefficients was calculated across repeats to evaluate precision.

#### Product of coefficients method

Two regression models were estimated. Firstly, the mediator was regressed on the exposure. Secondly, the outcome was regressed on the mediator, adjusting for the exposure. These two estimates were multiplied together to estimate the indirect effect.

#### Multivariable Mendelian randomisation

Using MVMR to estimate the direct effect, in the first stage regression, the effect of the instrument for the exposure and the weighted allele score for the mediator are used to predict each exposure respectively. In the second stage regression, the outcome was regressed on the predicted values of each exposure. The direct effect was then subtracted from the total effect, estimated using two-stage least squares regression, to estimate the indirect effect.

#### Two-step Mendelian randomisation

A univariable MR model was carried out to estimate the effect of the exposure on the mediator. A second model estimating the effect of the mediator on each outcome was carried out using MVMR. Both the genetic variants for the mediator and the exposure were included in the first and second stage regressions in MVMR. Previous approaches in the literature have not used MVMR for this second step [21, 23] and propose carrying out a univariable MR of the effect of the mediator on the outcome. However, using MVMR ensures any effect of the mediator on the outcome is independent of the exposure. Additionally, this method provides an estimate of the direct effect of the exposure on the outcome. The two regression estimates from the second stage regression are multiplied together to estimate the indirect effect.

#### Multiple Mediators

In phenotypic analyses, to estimate the direct effect attributable to multiple mediators, the outcome was regressed on the exposure, controlling for all mediators, using multivariable regression. Here, the coefficient for the exposure reflects the direct effect [31]. This direct effect was then subtracted from the total effect to estimate the indirect effect. Secondly, the product of coefficients method was used to estimate the indirect effect of each mediator individually. The combined effect of all three mediators was then estimated by summing together each individual effect.

In Mendelian randomisation analyses, the direct effect attributable to multiple mediators was assessed using MVMR, controlling for all mediators. This direct effect was then subtracted from the total effect to estimate the combined indirect effect. Secondly two-step MR was used, as previously described, considering each mediator individually and summing the effects together to obtain the indirect effect of all mediators combined.

#### Proportion mediated

The proportion mediated is calculated by dividing the indirect effect by the total effect. In individual-level MR, the confidence intervals can be estimated via bootstrapping.

## Testing the assumptions of mediation analysis

In this analysis, we have simulated a number of scenarios where phenotypic or MR methods for mediation analysis may provide biased answers. In this section we outline these results and any implications for analyses.

### Unmeasured confounding

Many of the key causal assumptions in phenotypic mediation analysis relate to assumptions of no unmeasured confounding between all of the exposure, mediator and outcome, including where confounders of the mediator and outcome are descendants of the exposure (intermediate confounding). Controlling for confounders in multivariable regression analyses often leads to residual confounding because it is generally impossible to measure all confounders, and frequently those that are measured are measured with error.

Indeed, in our simulations where residual covariance was simulated to reflect confounding, both the phenotypic difference method and phenotypic product of coefficients method were equally biased (Figure 3 and Online Resource 1: sTables 2). Where no confounding was simulated in the case of no true total effect, estimates from phenotypic approaches were free from bias (Online Resource 1: sTable 3). In simulations both with and without residual covariance to reflect confounding, MVMR and two-step MR estimated the direct effect, indirect effect and proportion mediated with no bias (Figure 2 and Online Resource 1: sTables 4 and 5).

**Figure 3:**
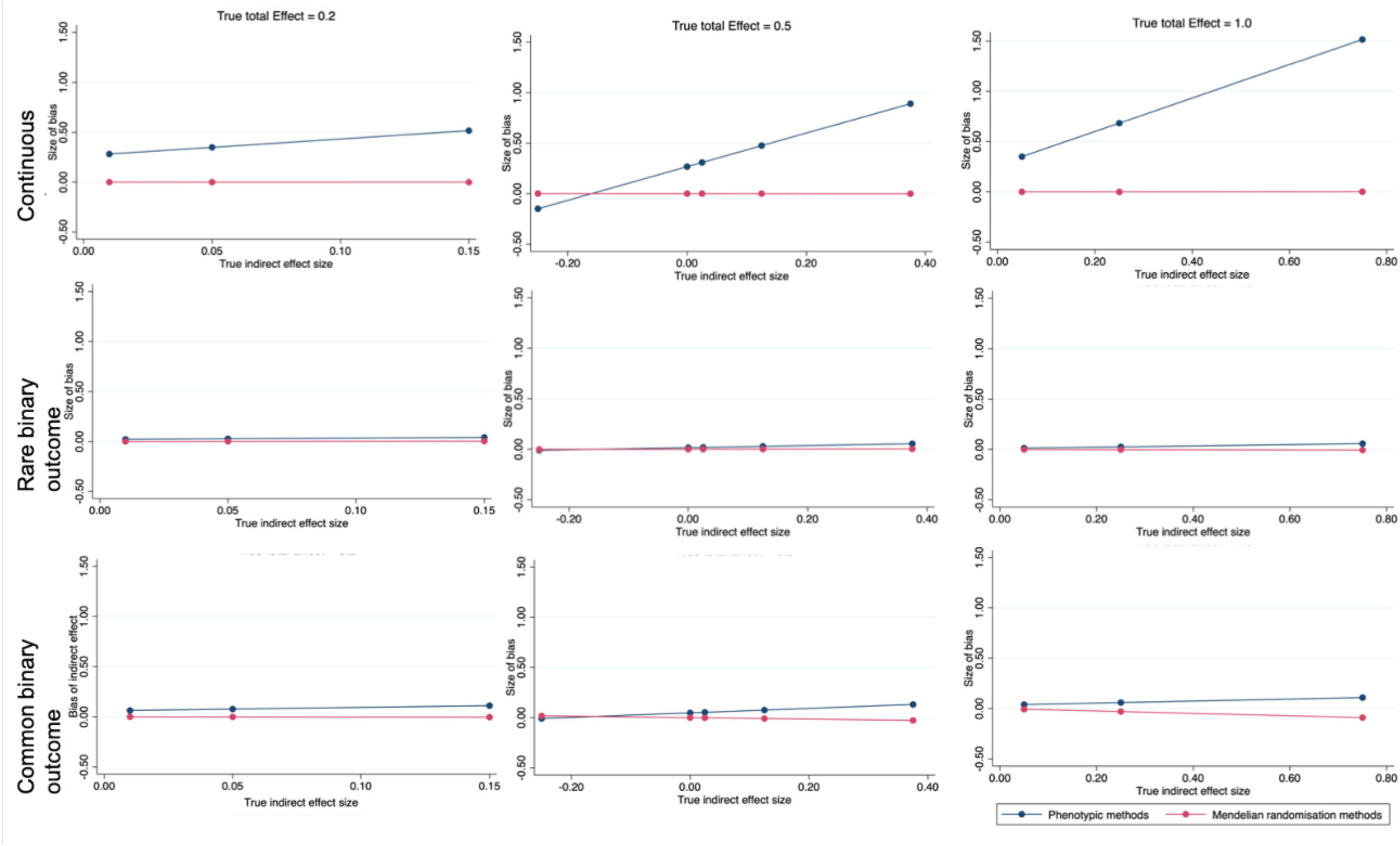
Size of absolute bias for the indirect effect of an exposure on range of outcomes through a continuous mediator, for a range of fixed true total effect sizes (0.2, 0.5 and 1.0) and range of true indirect effect sizes using phenotypic mediation methods or Mendelian randomisation, on the relative scale (simulated N = 5000)

Collider bias can be introduced by adjusting for the mediator in the presence of un-or mis-measured mediator-outcome confounders, where a backdoor path opens up between the exposure and the confounder (Online Resource 1: sFigure 3) [6, 32, 33]. Given that MR estimates are unbiased by unmeasured confounding of the exposure-outcome and mediator-outcome relationships [1, 17], this means that within MR analyses, adjusting for the mediator does not result in collider bias.

### Analysis of binary outcomes

Mediation analysis of binary outcome is challenging because of the non-collapsibility of odds ratio. This means the association between an exposure and outcome would not be constant on the odds-ratio scale by strata of categorical covariate [34, 35]. In mediation analysis, including the mediator in the model estimating the direct effect, means the model is no longer comparable with that for the total effect.

The mediation literature indicates that to estimate the direct and indirect effects of a binary outcome, the outcome must be rare (less than 10% prevalence), so the odds ratio approximates the risk ratio, and the product of coefficients method should be used for phenotypic data [9]. In the presence of a common binary outcome, estimates from the product of coefficients method and difference method are unlikely to align (and indeed the literature suggests both are likely biased) [4].

In our simulations, both the difference in coefficients and the product of coefficients phenotypic methods, with common and rare binary outcomes on a linear relative scale were biased as expected (Figure 3 and Online Resource 1: sTables 6-9). In simulated MR scenarios with common and rare binary outcomes on a linear relative scale, estimated effects were concordant between MVMR and two-step MR, with little to no bias (Figure 3 and Online Resource 1: sTables 10-13).

In the scenarios simulated, there was some bias when analysing binary outcomes on the log odds ratio scale using both MVMR and two-step MR, for both common and rare binary outcomes (Online Resource 1: sTables 14 and 15). This bias was small and typically would not alter conclusions made, although typically the size of absolute bias increased as the size of the true proportion mediator increased. However, the exact bias from non-collapsibility will be unique to each scenario, including depending on the strength of the mediators. Analyses in individual level MR can be conducted on the risk difference scale, which reduced bias due to non-collapsibility. In simulation scenarios explored, neither MVMR nor two-step MR were able to estimate the mediated effects without bias when using the odds ratio scale (Online Resource 1: sTables 16 and 17).

### Measurement error in the exposure or mediator

Our results show that in phenotypic approaches, with a continuous exposure and mediator, non-differential measurement error in the mediator leads to an underestimate of the mediated effect. This is consistent with previous methodological and applied work [8]. Where non-differential measurement error was simulated in the exposure, the mediated effect was over estimated (Online Resource 1: sTable 18).

In Mendelian randomisation simulations, both MVMR and two-step MR estimated the mediated effects with little bias when non-differential measurement error was simulated either in the exposure or the mediator (Online Resource 1: sTable 19). This is consistent with the previous literature demonstrating that MR estimates are less prone to bias by measurement error than conventional phenotypic analyses [1, 17].

### Weak instrument bias

In order to obtain valid causal inference for mediation, all standard MR assumptions must be met. This includes having strong instruments, typically determined through an F-statistic or conditional F-statistic of greater than 10. When the instruments in the simulation were weakly associated with the exposure both MVMR and two-step MR estimates of the indirect effect and proportion mediated were biased. The size of bias was greatest for a common binary outcome. When weak instruments were simulated for the mediator, estimates of the indirect effect and proportion mediated from both MVMR and two-step MR were biased (Online Resource 1: sFigure 4). Bias due to weak instruments have been discussed extensively in the literature [36-38], and some methods are now available for testing for weak instrument bias in MVMR [39].

**Figure 4:**
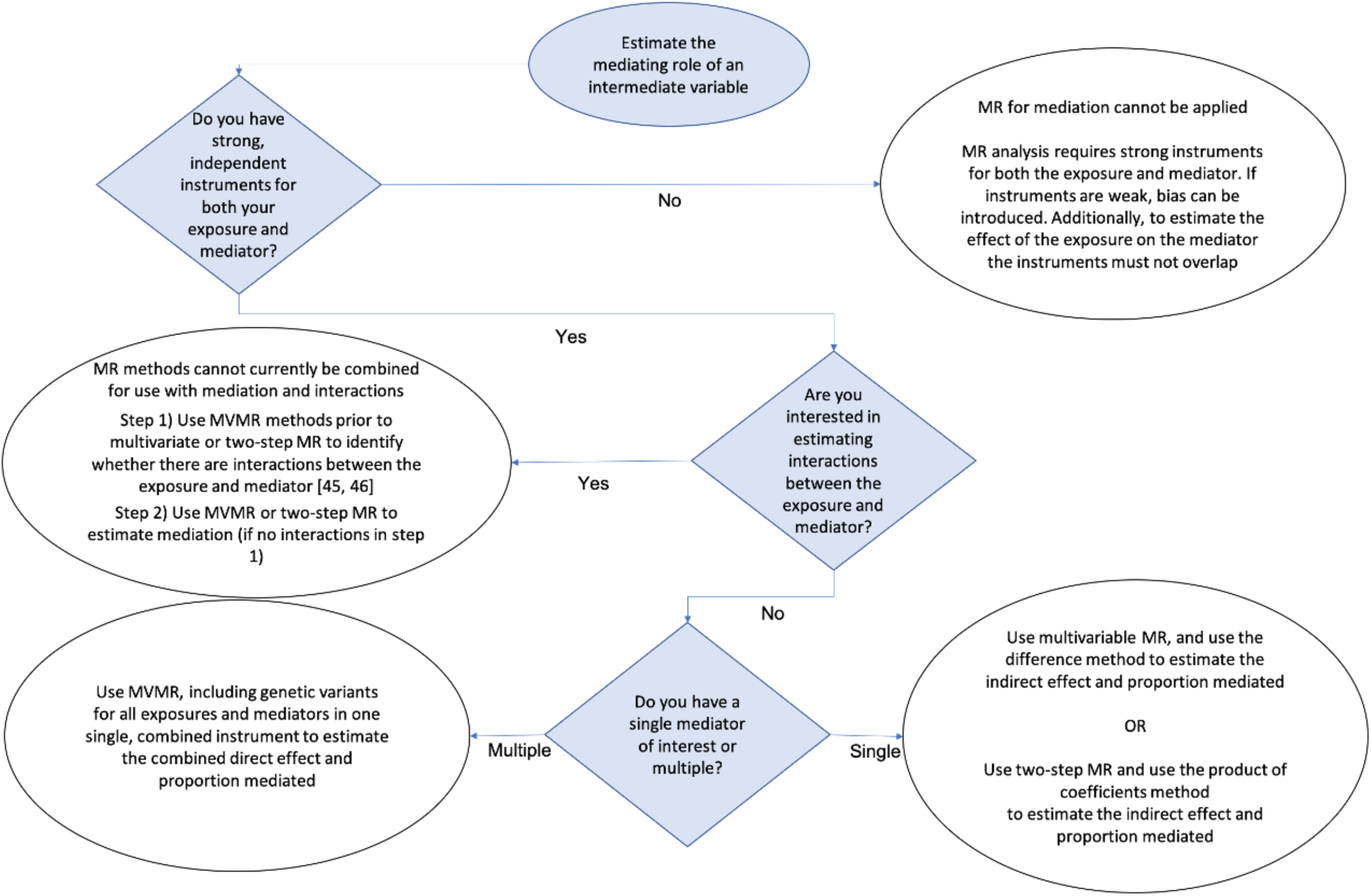
Decision flow chart to determine most appropriate mediation method

### Small total effects

In simulation studies with no true total effect the MR estimate of the proportion mediated is implausible (Online Resource 1: sTable 4). Where there is no evidence of a total effect, consideration should be given as to whether it is appropriate to continue with mediation analyses. Although an indirect effect can be estimated in the absence of a significant total effect, or absence of total effect when the indirect effect and direct effect act in opposing directions and cancel each other out, these estimates are prone to inflated type 1 errors (i.e. false positive results) [40].

Where the total effect is weak or estimated imprecisely (with confidence intervals crossing the null) simulations show the indirect effect and the proportion mediated using MR can be estimated but have large standard deviations (Online Resource 1: sTable 20-21). In this case, results should be interpreted with caution, especially considering the bounds of error.

## Analysis of multiple mediators

The direct effect of an exposure controlling for multiple mediators in a single model can be assessed using MVMR, with no evidence of bias (Online Resource 1: sTable 24). Here, non-overlapping SNPs for all exposures and mediators are included in one set of instruments. The estimated direct effect attributable to multiple mediators is unbiased, even in the presence of mediator-mediator relationships, which in our simulations was demonstrated by M2 causing M3 (Online Resource 1: sFigure 2).

Where there are no mediator-mediator relationships, estimates of the indirect effects and proportion mediated from both MVMR (mutually adjusting for all mediators) and two-step MR (considering each mediator individually and summing together) will coincide (Online Resource 1: sTable 24). In our simulations, both MR methods estimated the indirect effect of each mediator, and the three mediators jointly, with no bias (Online Resource 1: sTable 24). This is consistent with the existing literature on phenotypic multiple mediators [31].

Where mediator-mediator relationships are present, the indirect effect estimated via two-step MR captures both the amount of the association explained by the mediator of interest, and the amount of the mediator-outcome association captured by related mediators. In our example, this means that the effect of M3 is estimated twice, once directly and once via M2. As such, the estimate for the proportion mediated summing all three mediators together will likely be an overestimate of the combined proportion mediated, but the estimated direct effecteffects remain unbiased. In our simulations, the combined proportion mediated was over-estimated by 6% (Online Resource 1: sTable 24), which is equivalent to the proportion explained by M3 via M2. The indirect effect of M2 estimated using two-step MR is however unbiased and reflects both the direct effect of M2 on the outcome and the indirect effect via M3 (Online Resource 1: sFigure 2).

## Limitations of Mendelian randomisation applied to mediation analysis

### Instrument selection

Instruments associated with multiple exposures can be included in a MVMR analysis when MVMR is being used to test for potential pleiotropic pathways [24, 41, 42]. However, when MVMR is used to test for mediation, these overlapping instruments should not be included. If overlapping instruments were included and an attenuation of the direct effect compared with the total effect was observed, it would not be possible to distinguish whether this were attributable to mediation or pleiotropy (i.e. an effect of the SNP on the outcome via the mediator that is not due to the exposure). In a two-step MR mediation analysis, the mediator is considered as both an exposure (of the outcome) and as an outcome (of the exposure) and therefore any instruments for the exposure that are also instruments for the mediator are pleiotropic in the estimation of the effects of the exposure on the mediator and should be excluded. Where there are no independent SNPs, or the SNPs had a perfectly proportional effect on both the exposure and the mediator, then it would not be possible to use MR methods to estimate mediation.

The exclusion restriction criteria assuming no pleiotropic pathway is an important assumption of standard univariable MR, which applies equally when MR is used for mediation analysis. Some methods are available to assess pleiotropy including for the use of MVMR [43-45].

### Binary exposures and/or mediators

Very few binary exposures will be truly binary and are likely a dichotomization of an underlying liability, changing the interpretation of an MR analysis [46]. For example, smoking is often defined as ever versus never smokers, when the underlying exposure is a latent continuous variable reflecting smoking heaviness and duration. As a result, the exclusion restriction criteria are violated, where the genetic variant can influence the outcome via the latent continuous exposure, even if the binary exposure does not change [46]. In a mediation setting, the same would apply to a binary mediator. In these scenarios, two-step MR could be used to test whether there is evidence of a causal pathway between the binary exposure and/or mediator. However, the estimates of mediation would likely be biased.

### Interactions between the exposure and mediators

Within phenotypic analysis, exposure-mediator interactions can be accommodated when estimating mediation parameters. This is not possible in either MVMR or two-step MR Methods are available for estimating interactions in an MR framework with individual level data, but these do not currently extend to estimating mediation in the presence of exposure-mediator interactions [13, 47, 48]. Estimates of mediation from MR mediation methods will be require assuming effect homogeneity of both the exposure on the mediator and outcome, and mediator on the outcome. This means that the effects of the exposure and the mediator are the same for all individuals. Where interactions between the exposure and mediator are hypothesised this assumption may not hold true. Developing Mendelian randomisation methods which can account for these interactions will be important areas of future research.

### Power

Mendelian randomisation studies require very large sample sizes to achieve adequate statistical power. Conditional F-statistics in MVMR are typically weaker than standard F-statistics, and indeed are likely to become weaker with each additional mediator included, further decreasing the power of complex analyses. Therefore, to achieve adequate statistical power, or precision, sample sizes for mediation analysis likely need to be even larger than those needed in a univariable MR analyses.

In the absence of formal power calculators for complex MR scenarios, the power of these analyses can be considered by evaluating the precision of the confidence intervals for all of the total, direct and indirect effects, as well as assessing the conditional instrument strength.

### Confounding

Although assumptions about unmeasured confounding in MR can be relaxed compared with traditional phenotypic analyses, confounding can be introduced through population stratification, assortative mating, and dynastic effects [49]. Adjusting for genetic principal components and other explanatory variables that capture population structure or within family analyses can minimise bias.

## Mediation analysis with summary sample Mendelian randomisation

Methods applied in this paper can be used with summary data MR (see Box 1). Similar considerations will apply for both individual level MR, as presented here, and summary data MR. Importantly, all sources of summary statistics for the exposure, mediator and outcome should be non-overlapping [50]. As the mediator is considered an outcome in the exposure-mediator model, sample overlap can introduce bias [50]. As individual level data is not available in summary data MR, bootstrapping cannot be used to estimate the confidence intervals for the indirect effect or proportion mediated, but the delta method can be used to approximate these confidence intervals if samples are independent [30]. Analyses will also be restricted to the scale reported by the GWAS used, so consideration will need to be given for binary outcomes where sensitivity analyses to test potential non-collapsibility are limited.

## Which method and when

Although MR is robust to many of the untestable causal assumptions in phenotypic mediation analysis, these are replaced with a set of MR specific causal assumptions (Figure 2), and careful consideration should be given to which assumptions are most plausible. Additionally, the data available, or research question of interest may not be suitable to test in an MR framework. For example, where the research question is primarily interested in time varying exposures or mediators, MR becomes increasingly complex [51]. Mediation estimates from MR assume a time-fixed effect of the exposure and mediator, representing long-term relationships between the exposure and mediator [23]. In some unique cases instruments may be available for an exposure at different time points (e.g. childhood and adulthood BMI), but using these instruments come with additional methodological challenges [52].

Mendelian Randomisation has specific advantages compared with phenotypic methods where causal assumptions are required. The causal effect of the exposure on the outcome, the exposure on the mediator and the mediator on the outcome can all be tested. Additionally, bi-directional MR could be used to determine which of two variables is the causal exposure and causal mediator, where this is not known.

Our results demonstrate that both MVMR (akin to the difference in coefficients method) and two-step MR (akin to the product of coefficients method) can estimate the mediating effects for both continuous and binary outcomes, with little evidence of bias. However, caution is required in some instances, for example where total effects are weak. Where all exposures, mediators and outcomes are continuous, MVMR may confer an advantage of power, where the standard deviations for the simulated effects estimated in MVMR were smaller compared with the same effects estimated using two-step MR.

If an analysis is interested in estimating the effects of multiple mediators, consideration should be given to the causal question of interest when deciding which method to use to analyse multiple mediators. Where the causal question specifically relates to identifying the combined effects of multiple mediators, MVMR is likely to be the most appropriate method. Where the causal question aims to estimate the effect of multiple mediators individually, and potentially any impact of intervening on a mediator, two-step MR is likely to be most appropriate. However, it is important to note, that as the number of mediators included in an MVMR model increases, the power of the analysis would likely decrease. Additionally, future research should be carried out to determine if including increasing numbers of exposures in an MVMR model further violates any of the MR assumptions.

Although we have included a range of simulation scenarios, including both continuous and binary outcomes, this is not an exhaustive range of scenarios and there may be further scenarios where MR methods are biased.

The flow chart in Figure 4 aims to help with the decision-making process, based on practical limitations of MR. However, best practice would always be to triangulate across phenotypic and genetic approaches, and across multiple data sources wherever possible [53].

## Conclusions

Mendelian randomisation can be extended to estimate direct effects, indirect effects and proportions mediated. MR estimates are robust to violations of the often-untestable assumptions of phenotypic mediation analysis, including unmeasured confounding, reverse causality and measurement error. MR analysis makes its own strong, but distinct assumptions, especially relating to instrument validity. To estimate mediation using MR, we require large sample sizes and strong instruments for both the exposure and mediator.

#### Box 1: Summary of Mendelian randomisation

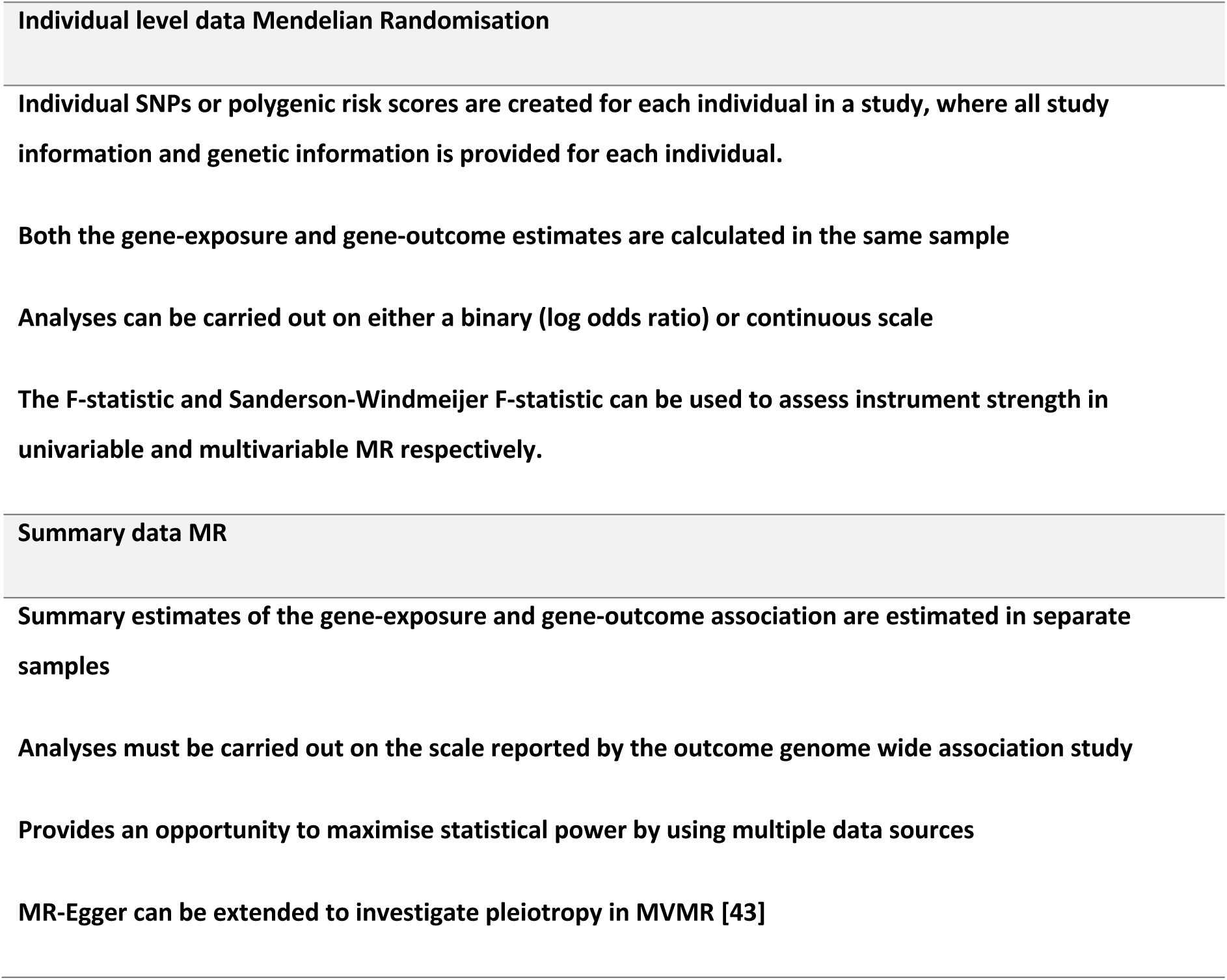

#### Box 2

**Key recommendations when using Mendelian randomisation for mediation analysis**

- Ensure strong instruments are available for exposures and mediators and test instrument strength using the F-statistic. Test the conditional instrument strength for multivariable MR using the Sanderson-Windmeijer F-statistic [54]
- Instruments for the exposure and mediator must be independent for both multivariable MR and two-step MR methods
- The instruments must not have a pleiotropic effect on the mediator or outcome
- Current MR methods are optimised for use with continuous exposures and mediators, where binary exposures or mediators which are a reflection of a true underlying continuous measure can lead to violation of the exclusion restriction criteria
- Use univariable MR to test for evidence of causal association in each step of the mediation path, from the exposure to the outcome, exposure to the mediator and mediator to the outcome
- Where individual-level data are being used and outcomes are binary estimate effects on a linear scale to alleviate potential bias from non-collapsibility of odds ratios
- If using summary level data with a binary outcome, estimate effects on the log odds ratio scale and transform after analysis if odds ratios are required

## Supporting information

Online Resource 1

Online Resource 2

## Declarations

### Conflicts of interest

There are no conflicts of interest

### Funding statement

No funding body has influenced data collection, analysis or its interpretations. This research was conducted using the UK Biobank Resource using application 10953. ARC is funded by the UK Medical Research Council Integrative Epidemiology Unit, University of Bristol (MC_UU_00011/1). All authors work in a unit that receives core funding from the UK Medical Research Council and University of Bristol (MC_UU_00011/1, MC_UU_00011/2, MC_UU_00011/3, MC_UU_00011/7). The Economics and Social Research Council support NMD via a Future Research Leaders grant (ES/N000757/1) and a Norwegian Research Council Grant number 295989. AET and GDS are supported by the National Institute for Health Research (NIHR) Biomedical Research Centre based at University Hospitals Bristol NHS Foundation and the University of Bristol. The views expressed are those of the authors and not necessarily those of the NHS, the NIHR, or the Department of Health. RCR is a de Pass Vice Chancellor’s Research Fellow at the University of Bristol. GH is supported by a Sir Henry Wellcome Postdoctoral Fellowship (209138/Z/17/Z). LDH is funded by a Career Development Award from the UK Medical Research Council (MR/M020894/1). This research has been conducted using the UK Biobank Resource under Application Number 10953.

### Data and code availability

All code for simulation analyses, applied analyses and example code is available on Github (simulations: https://github.com/eleanorsanderson/MediationMR, applied analyses and example code: https://github.com/alicerosecarter/MediationMR). The cleaned dataset for UK Biobank analyses will be archived with UK Biobank. Please contact for further information

### Ethical Approval

This project was approved by UK Biobank under the study application 10953. No specific ethical approval or patient involvement was sought for this project.

## Acknowledgments

We thank Kate Tilling for reading and commenting on an earlier draft of this manuscript

## Notes

### Competing Interest Statement

The authors have declared no competing interest.

### Summary of Updates

This manuscript has been revised to include simulations and an applied example of analysis the effect of multiple mediators using Mendelian randomisation mediation methods. Additionally, the organisation and layout of the manuscript has been amended to combine the results and discussion sections and provide two separate supplementary resources.

