## Supplementary material for "Mendelian randomisation for mediation analysis: current methods and challenges for implementation": Online Resource 1

#### Electronic Supplementary Material 1: Supplementary results and figures from simulation analyses

Alice R Carter [1,2]*, Eleanor Sanderson [1,2], Gemma Hammerton [1,2,3], Rebecca C Richmond [1,2], George Davey Smith [1,2,4], Jon Heron [1,2,3], Amy E Taylor [1,2,4], Neil M Davies [1,2,5], Laura D Howe [1,2]

1. MRC Integrative Epidemiology Unit, University of Bristol, Bristol, UK
2. Population Health Sciences, Bristol Medical School, University of Bristol, Bristol, UK
3. Centre for Academic Mental Health, University of Bristol, Bristol, UK
4. National Institute for Health Research Biomedical Research Centre at the University Hospitals Bristol NHS Foundation Trust and the University of Bristol, Bristol, UK
5. K.G. Jebsen Center for Genetic Epidemiology, Department of Public Health and Nursing, NTNU, Norwegian University of Science and Technology, Norway.

Corresponding Author:
Alice Carter

Oakfield House,
Oakfield Grove,
Bristol,
BS8 2BN
0117 3310098
ORCiD ID: 0000-0003-2817-4195

#### Supplementary digital content

#### Supplementary methods

Using the notation X = exposure, M = mediator, M1 = mediator 1, M2 = mediator 2, M3 – mediator 3, Y = outcome, G = genetic instruments, C = measured confounders, V = uncorrelated error term, μ = uncorrelated error term, four methods are compared (figure 1). Here we give the regression equations to estimate each of the methods described in the main paper. Notation and equations for the difference method and product of coefficients method are adapted from Vanderweele, 2015, where full details of the equations and notations are available [1]. Variables and parameters given in bold indicate the main coefficient(s) of interest in each case.

##### Single mediators

i. The difference method to estimate the direct effect (and infer the indirect effect) using phenotypic observed data

**Total:**

$$Y = \theta^{+}_0 + \boldsymbol{\theta^{+}_1 X} + \theta^{+}_3 C$$

**Direct:**

$$Y = \theta_0 + \boldsymbol{\theta_1 X} + \theta_2 M + \theta_4 C$$

**Indirect:**

$$\boldsymbol{\theta^{+}_1 - \theta_1}$$

ii. The product of coefficients method to estimate the indirect effect using phenotypic observed data

**Exposure-Mediator:**

$$M = \beta_0 + \boldsymbol{\beta_1 X} + \beta_3 C$$

**Direct:**

$$Y = \theta_0 + \boldsymbol{\theta_1 X} + \theta_2 M + \theta_4 C$$

**Indirect:**

$$\boldsymbol{\beta_1 \theta_2}$$

iii. Multivariable MR to estimate the direct effect and indirect effect using a single genetic instrumental variable for each of the exposure and mediator, using two-stage least squares regression

**Total:**

$$X = \pi_0 + \pi_1 G_x + v_1$$

$$Y = \beta_0 + \boldsymbol{\beta_{XT} X} + \mu_1$$

**Direct:**

$$X = \pi_0 + \pi_{1x} G_x + \pi_{2x} G_M + v_1$$

$$M = \pi_0 + \pi_{1z}G_x + \pi_{2z}G_M + v_2$$

$$Y = \beta_0 + \boldsymbol{\beta_X X} + \beta_M M + \mu_2$$

**Indirect:**

$$\boldsymbol{\beta_{XT} - \beta_X}$$

iv. Two-step MR to estimate the indirect effect using genetic instrumental variables for both the exposure and mediator, using two-stage least squares regression

**Exposure-Mediator:**

$$X = \pi_0 + \pi_1 G_x + v_1$$

$$M = \beta_0 + \boldsymbol{\beta_{XM} X} + \mu_1$$

**Direct**:

$$X = \pi_0 + \pi_{1x}G_x + \pi_{2x}G_M + v_1$$

$$M = \pi_0 + \pi_{1z}G_x + \pi_{2z}G_M + v_2$$

$$Y = \beta_0 + \boldsymbol{\beta_X X} + \beta_M M + \mu_2$$

**Indirect**:

$$\boldsymbol{\beta_{XM}\ \beta_M}$$

#### Multiple mediators

v. The difference method to estimate the direct effect and indirect effect using phenotypic observed data mutually adjusting for all mediators

**Total:**

$$Y = \theta^+_0 + \boldsymbol{\theta^+_1 X} + \theta^+_5 C$$

**Direct:**

$$Y = \theta_0 + \boldsymbol{\theta_1 X} + \theta_2 M1 + \theta_3 M2 + \theta_4 M3 + \theta_5 C$$

**Indirect:**

$$\boldsymbol{\theta^+_1 X - \theta_1 X}$$

vi. The product of coefficients method to estimate the indirect effect using phenotypic observed data, considering each mediator individually

**Mediator 1:**

**Exposure-Mediator:**

$$M1 = \beta_0 + \beta_1 X + \beta_4 C$$

**Direct:**

$$Y = \theta_0 + \boldsymbol{\theta_1 X} + \theta_{2M1}M1 + \theta_4 C$$

**Indirect:**

$$\boldsymbol{\beta_1\theta_{2M1}}$$

**Mediator 2:**

**Exposure-Mediator:**

$$M2 = \beta_0 + \boldsymbol{\beta_2 X} + \beta_4 C$$

**Direct:**

$$Y = \theta_0 + \boldsymbol{\theta_1 X} + \theta_{2M2}M2 + \theta_4 C$$

**Indirect:**

$$\boldsymbol{\beta_2\theta_{2M2}}$$

**Mediator 3:**

**Exposure-Mediator:**

$$M3 = \beta_0 + \beta_3 X + \beta_4 C$$

**Direct:**

$$Y = \theta_0 + \boldsymbol{\theta_1 X} + \theta_{2M3}M3 + \theta_4 C$$

**Indirect:**

$$\boldsymbol{\beta_3\theta_{2M3}}$$

**Combined indirect:**

$$\boldsymbol{\beta_1\theta_{2M1} + \beta_2\theta_{2M2} + \beta_3\theta_{2M3}}$$

vii. Multivariable MR to estimate the direct effect and indirect effect using a single genetic instrumental variable for each of the exposure and mediator, using two-stage least squares regression

**Total:**

$$X = \pi_0 + \pi_1 G_x + v_1$$

$$Y = \beta_0 + \boldsymbol{\beta_{XT} X} + \mu_1$$

**Direct:**

$$X = \pi_0 + \pi_{1x}G_x + \pi_{2x}G_{M1} + \pi_{3x}G_{M2} + \pi_{4x}G_{M3} + v_1$$

$$M1 = \pi_1 + \pi_{1z}G_x + \pi_{2z}G_{M1} + \pi_{3z}G_{M2} + \pi_{4z}G_{M3} + v_2$$

$$M2 = \pi_2 + \pi_{1\Omega}G_x + \pi_{2\Omega}G_{M1} + \pi_{3\Omega}G_{M2} + \pi_{4\Omega}G_{M3} + v_3$$

$$M3 = \pi_3 + \pi_{3\alpha}G_x + \pi_{2\alpha}G_{M1} + \pi_{2\alpha}G_{M2} + \pi_{4\alpha}G_{M3} + v_4$$

$$Y = \beta_0 + \boldsymbol{\beta_X X} + \beta_{M1}M1 + \beta_{M2}M2 + \beta_{M3}M3 + \mu_2$$

**Indirect:**

$$\boldsymbol{\beta_{XT} - \beta_X}$$

viii. Two-step MR to estimate the indirect effect using genetic instrumental variables for both the exposure and mediator, using two-stage least squares regression

**Mediator 1:**

**Exposure-Mediator:**

$$X = \pi_0 + \pi_1 G_x + v_X$$

$$M1 = \beta_0 + \boldsymbol{\beta_{XM1} X} + \mu_1$$

**Direct:**

$$X = \pi_0 + \pi_{1x}G_x + \pi_{2x}G_{M1} + v_{X1}$$

$$M1 = \pi_{01} + \pi_{11}G_x + \pi_{21}G_{M1} + v_{M1}$$

$$Y = \beta_0 + \boldsymbol{\beta_{X1} X} + \beta_{M1}M1 + \mu_2$$

**Indirect:**

$$\boldsymbol{\beta_{XM1}\beta_{M1}}$$

**Mediator 2:**

**Exposure-Mediator:**

$$X = \pi_0 + \pi_1 G_x + v_X$$

$$M2 = \beta_0 + \boldsymbol{\beta_{xM2} X} + \mu_3$$

**Direct:**

$$X = \pi_{02} + \pi_{12}G_x + \pi_{22}G_{M2} + v_{X2}$$

$$M2 = \pi_{0M2} + \pi_{1M2}G_x + \pi_{2M2}G_{M2} + v_{M2}$$

$$Y = \beta_1 + \boldsymbol{\beta_{X2} X} + \beta_{M2}M2 + \mu_4$$

**Indirect:**

$$\boldsymbol{\beta_{XM2}\beta_{M2}}$$

**Mediator 3:**

**Exposure-Mediator:**

$$X = \pi_0 + \pi_1 G_x + v_X$$

$$M3 = \beta_0 + \boldsymbol{\beta_{XM3}X} + \mu_3$$

**Direct:**

$$X = \pi_{03} + \pi_{13}\mathbf{G}_x + \pi_{23}G_{M3} + v_{X3}$$

$$M3 = \pi_{0M3} + \pi_{1M3}G_x + \pi_{2M3}\mathbf{G_{M2}} + v_{M3}$$

$$Y = \beta_2 + \boldsymbol{\beta_{X3}X} + \beta_{M3}M3 + \mu_6$$

**Indirect:**

$$\boldsymbol{\beta_{XM3}\beta_{M3}}$$

**Combined indirect:**

$$\boldsymbol{\beta_{XM1}\beta_{M1} + \beta_{XM2}\beta_{M2} + \beta_{XM3}\beta_{M3}}$$

### Supplementary Tables

#### sTable 1: Simulation scenarios

| | **Total effect** | **Proportion mediated** | | | | | **Sample Size** | **Measurement error** | **Weak instrument** |
|---|---|---|---|---|---|---|---|---|---|
| No Mediation | 0.5 | 0 | | | | | 5000 | | |
| Inconsistent mediation | 0.5 | -0.5 | | | | | 5000 | | |
| Varying total effect | 0 | 0.05 | 0.25 | 0.75 | | | 5000 | | |
| | 0.2 | 0.05 | 0.25 | 0.75 | | | 5000 | | |
| | 0.5 | 0.05 | 0.25 | 0.75 | | | 5000 | | |
| | 1 | 0.05 | 0.25 | 0.75 | | | 5000 | | |
| Small total effect | 0.01 | 0.05 | 0.25 | 0.75 | | | 5000 | | |
| | 0.05 | 0.05 | 0.25 | 0.75 | | | 5000 | | |
| | 0.1 | 0.05 | 0.25 | 0.75 | | | 5000 | | |
| Imprecise total effect | 0.2 | 0.05 | 0.25 | 0.75 | | | 1000 | | |
| Measurement error | 0.5 | 0.25 | | | | | 5000 | Exposure | |
| | 0.5 | 0.25 | | | | | 5000 | Mediator | |
| Weak instrument bias | 0.5 | 0.25 | | | | | 5000 | | Exposure |
| | 0.5 | 0.25 | | | | | 5000 | | Mediator |
| No confounding* | 0 | 0.05 | 0.25 | 0.75 | | | 5000 | | |
| Multiple mediators | | **Joint** | **M1** | **M2** | **M3** | **M3 via M2** | 5000 | | |
| | 0.45 | 0.56 | 0.11 | 0.18 | 0.12 | 0 | | | |
| | | 0.56 | 0.11 | 0.18 | 0.27 | 0.06 | | | |

In all simulations the effect of the mediator on the outcome is set to 0.2. All simulations undergo 1000 replications.

Confounding is simulated as residual covariance between the exposure, mediator and outcome in all scenarios except * sTable 2: Estimated effect sizes and size of bias for simulated effect of a phenotypically measured continuous mediator explaining the effect between a continuous exposure and continuous outcome (Simulated N=5000)

| True total effect | True proportion mediated | Total effect (SD) | Size of bias (absolute) | Size of bias (relative) | Direct effect (SD) | Size of bias (absolute) | Size of bias (relative) | Indirect effect - difference method (SD) | Size of bias (absolute) | Size of bias (relative) | Proportion mediated - difference (SD) | Size of bias (absolute) | Size of bias (relative) | Indirect effect - product of coefficients (SD) | Size of bias (absolute) | Size of bias (relative) | Proportion mediated - product of coefficients (SD) | Size of bias (absolute) | Size of bias (relative) |
|---|---|---|---|---|---|---|---|---|---|---|---|---|---|---|---|---|---|---|---|
| 0.5 | 0 | 1.1 (0.009) | 0.60 | 1.20 | 0.833 (0.007) | 0.33 | 0.67 | 0.267 (0.007) | 0.27 | | 0.243 (0.006) | 0.24 | | 0.267 (0.007) | 0.27 | | 0.243 (0.006) | 0.24 | 0.12 |
| 0.5 | -0.5 | 1.1 (0.009) | 0.60 | 1.20 | 1.5 (0.007) | 0.75 | 1.50 | -0.4 (0.008) | -1.15 | 4.60 | -0.364 (0.009) | 0.14 | -0.27 | -0.4 (0.008) | -1.15 | 4.60 | -0.364 (0.009) | 0.14 | 0.07 |
| 0 | 0.05 | 0.6 (0.009) | 0.60 | | 0.333 (0.007) | 0.33 | | 0.267 (0.007) | 0.27 | | 0.445 (0.009) | 0.39 | 7.90 | 0.267 (0.007) | 0.27 | | 0.445 (0.009) | 0.39 | 0.20 |
| 0.2 | 0.05 | 0.8 (0.009) | 0.80 | 4.00 | 0.507 (0.007) | 0.32 | 1.58 | 0.293 (0.007) | 0.28 | 28.33 | 0.367 (0.007) | 0.32 | 6.33 | 0.293 (0.007) | 0.28 | 28.33 | 0.367 (0.007) | 0.32 | 0.16 |
| 0.5 | 0.05 | 1.1 (0.009) | 0.90 | 1.80 | 0.767 (0.007) | 0.29 | 0.58 | 0.333 (0.008) | 0.31 | 12.32 | 0.303 (0.006) | 0.25 | 5.05 | 0.333 (0.008) | 0.31 | 12.32 | 0.303 (0.006) | 0.25 | 0.13 |
| 1 | 0.05 | 1.6 (0.009) | 1.10 | 1.10 | 1.2 (0.007) | 0.25 | 0.25 | 0.4 (0.008) | 0.35 | 7.00 | 0.25 (0.004) | 0.20 | 4.00 | 0.4 (0.008) | 0.35 | 7.00 | 0.25 (0.004) | 0.20 | 0.10 |
| 0 | 0.25 | 0.6 (0.008) | 0.60 | | 0.334 (0.006) | 0.33 | | 0.267 (0.007) | 0.27 | | 0.444 (0.009) | 0.19 | 0.78 | 0.267 (0.007) | 0.27 | | 0.444 (0.009) | 0.19 | 0.10 |
| 0.2 | 0.25 | 0.8 (0.009) | 0.80 | 4.00 | 0.4 (0.007) | 0.25 | 1.25 | 0.399 (0.008) | 0.35 | 6.99 | 0.499 (0.008) | 0.25 | 1.00 | 0.399 (0.008) | 0.35 | 6.99 | 0.499 (0.008) | 0.25 | 0.12 |
| 0.5 | 0.25 | 1.1 (0.009) | 0.90 | 1.80 | 0.5 (0.01) | 0.12 | 0.25 | 0.6 (0.01) | 0.48 | 3.80 | 0.546 (0.008) | 0.30 | 1.18 | 0.6 (0.01) | 0.48 | 3.80 | 0.546 (0.008) | 0.30 | 0.15 |
| 1 | 0.25 | 1.6 (0.009) | 1.10 | 1.10 | 0.667 (0.013) | -0.08 | -0.08 | 0.933 (0.014) | 0.68 | 2.73 | 0.583 (0.008) | 0.33 | 1.33 | 0.933 (0.014) | 0.68 | 2.73 | 0.583 (0.008) | 0.33 | 0.17 |
| 0 | 0.75 | 0.6 (0.009) | 0.60 | | 0.333 (0.007) | 0.33 | | 0.267 (0.007) | 0.27 | | 0.445 (0.009) | -0.31 | -0.41 | 0.267 (0.007) | 0.27 | | 0.445 (0.009) | -0.31 | -0.15 |
| 0.2 | 0.75 | 0.8 (0.009) | 0.80 | 4.00 | 0.134 (0.01) | 0.08 | 0.42 | 0.666 (0.011) | 0.52 | 3.44 | 0.833 (0.012) | 0.08 | 0.11 | 0.666 (0.011) | 0.52 | 3.44 | 0.833 (0.012) | 0.08 | 0.04 |
| 0.5 | 0.75 | 1.1 (0.008) | 0.90 | 1.80 | -0.166 (0.017) | -0.29 | -0.58 | 1.267 (0.017) | 0.89 | 2.38 | 1.151 (0.016) | 0.40 | 0.54 | 1.267 (0.017) | 0.89 | 2.38 | 1.151 (0.016) | 0.40 | 0.20 |
| 1 | 0.75 | 1.6 (0.009) | 1.10 | 1.10 | -0.666 (0.03) | -0.92 | -0.92 | 2.266 (0.03) | 1.52 | 2.02 | 1.416 (0.019) | 0.67 | 0.89 | 2.266 (0.03) | 1.52 | 2.02 | 1.416 (0.019) | 0.67 | 0.33 |

sTable 3: Estimated effect sizes and size of bias for simulated effect of a phenotypically measured continuous mediator explaining the effect between a continuous exposure and continuous outcome (per unit increase in exposure), and a rare binary outcome and common binary outcome on the risk difference scale, with no residual covariance reflecting confounding (Simulated N=5000)

| | **True total effect** | **True proportion mediated** | **Total effect (SD)** | **Size of bias (absolute)** | **Size of bias (relative)** | **Direct effect (SD)** | **Size of bias (absolute)** | **Size of bias (relative)** | **Indirect effect - difference method (SD)** | **Size of bias (absolute)** | **Size of bias (relative)** | **Proportion mediated - difference (SD)** | **Size of bias (absolute)** | **Size of bias (relative)** | **Indirect effect - product of coefficients (SD)** | **Size of bias (absolute)** | **Size of bias (relative)** | **Proportion mediated - product of coefficients (SD)** | **Size of bias (absolute)** | **Size of bias (relative)** |
|---|---|---|---|---|---|---|---|---|---|---|---|---|---|---|---|---|---|---|---|---|
| **Continuous outcome** | 0 | 0.05 | 0 (0.01) | 0.00 | | 0 (0.01) | 0.00 | | 0 (0.01) | 0.00 | | 0 (0.01) | 0.18 | 3.53 | 0 (0.003) | 0.00 | | 0.227 (10.342) | 0.18 | 0.09 |
| | 0 | 0.25 | 0 (0.01) | 0.00 | | 0 (0.01) | 0.00 | | 0 (0.01) | 0.00 | | 0 (0.01) | -0.10 | -0.39 | 0 (0.003) | 0.00 | | 0.152 (3.448) | -0.10 | -0.05 |
| | 0 | 0.75 | 0 (0.01) | 0.00 | | 0 (0.01) | 0.00 | | 0 (0.01) | 0.00 | | 0 (0.01) | -1.18 | -1.57 | 0 (0.003) | 0.00 | | -0.427 (24.509) | -1.18 | -0.59 |
| **Rare binary outcome** | 0 | 0.05 | 0 (0.002) | 0.00 | | 0 (0.002) | 0.00 | | 0 (0.002) | 0.00 | | 0 (0.002) | -0.39 | -7.83 | 0 (0) | 0.00 | | -0.342 (13.323) | -0.39 | -7.83 |
| | 0 | 0.25 | 0 (0.002) | 0.00 | | 0 (0.002) | 0.00 | | 0 (0.002) | 0.00 | | 0 (0.002) | -0.18 | -0.71 | 0 (0) | 0.00 | | 0.073 (3.212) | -0.18 | -0.71 |
| | 0 | 0.75 | 0 (0.002) | 0.00 | | 0 (0.002) | 0.00 | | 0 (0.002) | 0.00 | | 0 (0.002) | -0.48 | -0.65 | 0 (0) | 0.00 | | 0.266 (5.102) | -0.48 | -0.65 |
| **Common binary outcome** | 0 | 0.05 | 0 (0.004) | 0.00 | | 0 (0.004) | 0.00 | | 0 (0.004) | 0.00 | | 0 (0.004) | -0.01 | -0.11 | 0 (0.001) | 0.00 | | 0.044 (3.642) | -0.01 | -0.11 |
| | 0 | 0.25 | 0 (0.004) | 0.00 | | 0 (0.004) | 0.00 | | 0 (0.004) | 0.00 | | 0 (0.004) | -0.13 | -0.50 | 0 (0.001) | 0.00 | | 0.125 (4.933) | -0.13 | -0.50 |
| | 0 | 0.75 | 0 (0.004) | 0.00 | | 0 (0.004) | 0.00 | | 0 (0.004) | 0.00 | | 0 (0.004) | -0.64 | -0.85 | 0 (0.001) | 0.00 | | 0.114 (9.155) | -0.64 | -0.85 |

sTable 4: Estimated effect sizes and size of bias for simulated effect of a continuous mediator explaining the effect between a continuous exposure and continuous outcome using Mendelian Randomization (Simulated N=5000)

| True total effect | True proportion mediated | Total effect from univariate Mendelian randomisation (SD) | Size of bias (absolute) | Size of bias (relative) | Direct effect from multivariable Mendelian randomisation (SD) | Size of bias (absolute) | Size of bias (relative) | Indirect effect from multivariable Mendelian randomisation (SD) | Size of bias (absolute) | Size of bias (relative) | Proportion mediated from multivariable Mendelian randomisation (SD) | Size of bias (absolute) | Size of bias (relative) | Indirect effect from two-step Mendelian randomisation (SD) | Size of bias (absolute) | Size of bias (relative) | Proportion mediated from two-step Mendelian randomisation SD) | Size of bias (absolute) | Size of bias (relative) |
|---|---|---|---|---|---|---|---|---|---|---|---|---|---|---|---|---|---|---|---|
| 0.5 | 0 | 0.499 (0.017) | 0.00 | 0.00 | 0.5 (0.014) | 0.00 | 0.00 | 0 (0.004) | 0.00 | | 0 (0.008) | 0.00 | | 0 (0.004) | 0.00 | | 0.004 (0) | 0.00 | |
| 0.5 | -0.5 | 0.499 (0.017) | 0.00 | 0.00 | 0.75 (0.023) | 0.00 | 0.00 | -0.25 (0.018) | 0.00 | 0.00 | -0.501 (0.043) | 0.00 | 0.00 | -0.25 (0.018) | 0.00 | 0.00 | -0.501 (0.043) | 0.00 | 0.00 |
| 0 | 0.05 | 0 (0.017) | 0.00 | | 0 (0.014) | 0.00 | | 0 (0.004) | 0.00 | | 0.164 (2.176) | 0.11 | 2.28 | 0 (0.004) | 0.00 | | 0.004 (0.164) | 0.11 | 2.28 |
| 0.2 | 0.05 | 0.2 (0.017) | 0.00 | 0.00 | 0.19 (0.014) | 0.00 | 0.00 | 0.01 (0.004) | 0.00 | 0.01 | 0.049 (0.017) | 0.00 | -0.01 | 0.01 (0.004) | 0.00 | 0.01 | 0.004 (0.049) | 0.00 | -0.01 |
| 0.5 | 0.05 | 0.5 (0.018) | 0.00 | 0.00 | 0.475 (0.015) | 0.00 | 0.00 | 0.025 (0.004) | 0.00 | -0.01 | 0.05 (0.008) | 0.00 | -0.01 | 0.025 (0.004) | 0.00 | -0.01 | 0.004 (0.05) | 0.00 | -0.01 |
| 1 | 0.05 | 1 (0.017) | 0.00 | 0.00 | 0.95 (0.014) | 0.00 | 0.00 | 0.05 (0.005) | 0.00 | 0.00 | 0.05 (0.005) | 0.00 | 0.00 | 0.05 (0.005) | 0.00 | 0.00 | 0.005 (0.05) | 0.00 | 0.00 |
| 0 | 0.25 | 0 (0.018) | 0.00 | | 0 (0.014) | 0.00 | | 0 (0.004) | 0.00 | | 0.111 (3.554) | -0.14 | -0.56 | 0 (0.004) | 0.00 | | 0.004 (0.111) | -0.14 | -0.56 |
| 0.2 | 0.25 | 0.199 (0.017) | 0.00 | -0.01 | 0.149 (0.014) | 0.00 | 0.00 | 0.049 (0.005) | 0.00 | -0.01 | 0.249 (0.022) | 0.00 | 0.00 | 0.049 (0.005) | 0.00 | -0.01 | 0.005 (0.249) | 0.00 | 0.00 |
| 0.5 | 0.25 | 0.5 (0.017) | 0.00 | 0.00 | 0.375 (0.017) | 0.00 | 0.00 | 0.125 (0.01) | 0.00 | 0.00 | 0.25 (0.019) | 0.00 | 0.00 | 0.125 (0.01) | 0.00 | 0.00 | 0.01 (0.25) | 0.00 | 0.00 |
| 1 | 0.25 | 1.001 (0.017) | 0.00 | 0.00 | 0.751 (0.023) | 0.00 | 0.00 | 0.249 (0.018) | 0.00 | 0.00 | 0.249 (0.018) | 0.00 | 0.00 | 0.249 (0.018) | 0.00 | 0.00 | 0.018 (0.249) | 0.00 | 0.00 |
| 0 | 0.75 | 0 (0.017) | 0.00 | | 0 (0.014) | 0.00 | | 0 (0.004) | 0.00 | | -0.046 (4.949) | -0.80 | -1.06 | 0 (0.004) | 0.00 | | 0.004 (-0.046) | -0.80 | -1.06 |
| 0.2 | 0.75 | 0.2 (0.018) | 0.00 | 0.00 | 0.05 (0.018) | 0.00 | 0.00 | 0.15 (0.011) | 0.00 | 0.00 | 0.754 (0.074) | 0.00 | 0.01 | 0.15 (0.011) | 0.00 | 0.00 | 0.011 (0.754) | 0.00 | 0.01 |
| 0.5 | 0.75 | 0.501 (0.017) | 0.00 | 0.00 | 0.126 (0.029) | 0.00 | 0.01 | 0.374 (0.026) | 0.00 | 0.00 | 0.748 (0.056) | 0.00 | 0.00 | 0.374 (0.026) | 0.00 | 0.00 | 0.026 (0.748) | 0.00 | 0.00 |
| 1 | 0.75 | 1 (0.017) | 0.00 | 0.00 | 0.249 (0.057) | 0.00 | 0.00 | 0.751 (0.055) | 0.00 | 0.00 | 0.751 (0.056) | 0.00 | 0.00 | 0.751 (0.055) | 0.00 | 0.00 | 0.055 (0.751) | 0.00 | 0.00 |

sTable 5: Estimated effect sizes and size of bias for simulated effect of a continuous mediator explaining the effect between a continuous exposure and continuous outcome (per unit increase in exposure), and a rare binary outcome and common binary outcome on the risk difference scale using Mendelian Randomization, where no residual covariance is included reflecting confounding (Simulated N=5000)

| | True total effect | True proportion mediated | Total effect from univariate Mendelian randomisation (SD) | Size of bias (absolute) | Size of bias (relative) | Direct effect from multivariable Mendelian randomisation (SD) | Size of bias (absolute) | Size of bias (relative) | Indirect effect from multivariable Mendelian randomisation (SD) | Size of bias (absolute) | Size of bias (relative) | Proportion mediated from multivariable Mendelian randomisation (SD) | Size of bias (absolute) | Size of bias (relative) | Indirect effect from two-step Mendelian randomisation (SD) | Size of bias (absolute) | Size of bias (relative) | Proportion mediated from two-step Mendelian randomisation (SD) | Size of bias (absolute) | Size of bias (relative) |
|---|---|---|---|---|---|---|---|---|---|---|---|---|---|---|---|---|---|---|---|---|
| **Continuous outcome** | 0 | 0.05 | 0 (0.003) | 0.00 | | 0.227 (10.342) | 0.00 | | 0 (0.015) | 0.00 | | 0.811 (24.209) | 0.76 | 15.22 | 0 (0.004) | 0.00 | | 0.811 (24.209) | 0.76 | 15.22 |
| | 0 | 0.25 | 0 (0.003) | 0.00 | | 0.152 (3.448) | 0.00 | | 0 (0.015) | 0.00 | | -0.672 (31.009) | -0.92 | -3.69 | 0 (0.004) | 0.00 | | -0.672 (31.009) | -0.92 | -3.69 |
| | 0 | 0.75 | 0 (0.003) | 0.00 | | -0.427 (24.509) | 0.00 | | 0 (0.015) | 0.00 | | -0.022 (5.39) | -0.77 | -1.03 | 0 (0.004) | 0.00 | | -0.022 (5.39) | -0.77 | -1.03 |
| **Rare binary outcome** | 0 | 0.05 | 0 (0) | 0.00 | | -0.342 (13.323) | 0.00 | | 0 (0.003) | 0.00 | | 0 (0.003) | 1.29 | 25.90 | 0 (0) | 0.00 | | 1.345 (43.052) | 1.29 | 25.90 |
| | 0 | 0.25 | 0 (0) | 0.00 | | 0.073 (3.212) | 0.00 | | 0 (0.003) | 0.00 | | 0 (0.003) | -0.21 | -0.85 | 0 (0) | 0.00 | | 0.039 (1.697) | -0.21 | -0.85 |
| | 0 | 0.75 | 0 (0) | 0.00 | | 0.266 (5.102) | 0.00 | | 0 (0.003) | 0.00 | | 0 (0.003) | -0.81 | -1.08 | 0 (0) | 0.00 | | -0.059 (3.035) | -0.81 | -1.08 |
| **Common binary outcome** | 0 | 0.05 | 0 (0.001) | 0.00 | | 0.044 (3.642) | 0.00 | | 0 (0.006) | 0.00 | | 0 (0.006) | 0.78 | 15.57 | 0 (0.001) | 0.00 | | 0.829 (30.515) | 0.78 | 15.57 |
| | 0 | 0.25 | 0 (0.001) | 0.00 | | 0.125 (4.933) | 0.00 | | 0 (0.006) | 0.00 | | 0 (0.006) | -0.24 | -0.96 | 0 (0.001) | 0.00 | | 0.01 (4.135) | -0.24 | -0.96 |
| | 0 | 0.75 | 0 (0.001) | 0.00 | | 0.114 (9.155) | 0.00 | | 0 (0.006) | 0.00 | | 0 (0.006) | -0.19 | -0.25 | 0 (0.001) | 0.00 | | 0.565 (59.327) | -0.19 | -0.25 |

sTable 6: Estimated effect sizes and size of bias for simulated effect of a phenotypically measured continuous mediator explaining the effect between a continuous exposure and rare binary outcome on the risk difference scale (Simulated N=5000)

| True total effect | True proportion mediated | Total effect from univariate Mendelian randomisation (SD) | Size of bias (absolute) | Size of bias (relative) | Direct effect from multivariable Mendelian randomisation (SD) | Size of bias (absolute) | Size of bias (relative) | Indirect effect from multivariable Mendelian randomisation (SD) | Size of bias (absolute) | Size of bias (relative) | Proportion mediated from multivariable Mendelian randomisation (SD) | Size of bias (absolute) | Size of bias (relative) | Indirect effect from two-step Mendelian randomisation (SD) | Size of bias (absolute) | Size of bias (relative) | Proportion mediated from two-step Mendelian randomisation SD) | Size of bias (absolute) | Size of bias (relative) |
|---|---|---|---|---|---|---|---|---|---|---|---|---|---|---|---|---|---|---|---|
| 0.025 | 0 | 0.064 (0.001) | 0.04 | 1.54 | 0.048 (0.002) | 0.02 | 0.92 | 0.015 (0.001) | 0.02 | | 0.244 (0.019) | 0.24 | | 0.015 (0.001) | 0.02 | | 0.244 (0.019) | 0.24 | |
| 0.025 | -0.5 | 0.064 (0.001) | 0.04 | 1.54 | 0.087 (0.002) | 0.05 | 1.31 | -0.023 (0.002) | -0.06 | 4.86 | -0.365 (0.029) | 0.13 | -0.27 | -0.023 (0.002) | -0.06 | 4.86 | -0.365 (0.029) | 0.13 | -0.27 |
| 0 | 0.05 | 0.051 (0.002) | 0.05 | | 0.028 (0.002) | 0.03 | | 0.023 (0.001) | 0.02 | | 0.447 (0.029) | 0.40 | 7.93 | 0.023 (0.001) | 0.02 | | 0.447 (0.029) | 0.40 | 7.93 |
| 0.1 | 0.05 | 0.058 (0.001) | -0.04 | -0.42 | 0.037 (0.002) | -0.06 | -0.61 | 0.021 (0.001) | 0.02 | 3.24 | 0.367 (0.025) | 0.32 | 6.35 | 0.021 (0.001) | 0.02 | 3.24 | 0.367 (0.025) | 0.32 | 6.35 |
| 0.025 | 0.05 | 0.064 (0.001) | 0.04 | 1.54 | 0.044 (0.002) | 0.02 | 0.86 | 0.019 (0.001) | 0.02 | 14.43 | 0.304 (0.025) | 0.25 | 5.07 | 0.019 (0.001) | 0.02 | 14.43 | 0.304 (0.025) | 0.25 | 5.07 |
| 0.05 | 0.05 | 0.068 (0.001) | 0.02 | 0.36 | 0.051 (0.002) | 0.00 | 0.07 | 0.017 (0.002) | 0.01 | 5.83 | 0.251 (0.026) | 0.20 | 4.02 | 0.017 (0.002) | 0.01 | 5.83 | 0.251 (0.026) | 0.20 | 4.02 |
| 0 | 0.25 | 0.051 (0.002) | 0.05 | | 0.028 (0.002) | 0.03 | | 0.023 (0.001) | 0.02 | | 0.445 (0.028) | 0.19 | 0.78 | 0.023 (0.001) | 0.02 | | 0.445 (0.028) | 0.19 | 0.78 |
| 0.1 | 0.25 | 0.058 (0.001) | -0.04 | -0.21 | 0.029 (0.002) | -0.05 | -0.61 | 0.029 (0.002) | 0.00 | 0.16 | 0.5 (0.034) | 0.25 | 1.00 | 0.029 (0.002) | 0.00 | 0.16 | 0.5 (0.034) | 0.25 | 1.00 |
| 0.025 | 0.25 | 0.064 (0.001) | 0.04 | 1.55 | 0.029 (0.003) | 0.01 | 0.55 | 0.035 (0.002) | 0.03 | 4.54 | 0.545 (0.043) | 0.29 | 1.18 | 0.035 (0.002) | 0.03 | 4.54 | 0.545 (0.043) | 0.29 | 1.18 |
| 0.05 | 0.25 | 0.068 (0.001) | 0.02 | 0.36 | 0.029 (0.004) | -0.01 | -0.24 | 0.039 (0.004) | 0.03 | 2.16 | 0.581 (0.061) | 0.33 | 1.32 | 0.039 (0.004) | 0.03 | 2.16 | 0.581 (0.061) | 0.33 | 1.32 |
| 0 | 0.75 | 0.051 (0.002) | 0.05 | | 0.028 (0.002) | 0.03 | | 0.023 (0.001) | 0.02 | | 0.447 (0.028) | -0.30 | -0.40 | 0.023 (0.001) | 0.02 | | 0.447 (0.028) | -0.30 | -0.40 |
| 0.1 | 0.75 | 0.058 (0.001) | -0.04 | -0.21 | 0.01 (0.003) | -0.02 | -0.61 | 0.048 (0.002) | -0.03 | -0.36 | 0.833 (0.054) | 0.08 | 0.11 | 0.048 (0.002) | -0.03 | -0.36 | 0.833 (0.054) | 0.08 | 0.11 |
| 0.025 | 0.75 | 0.064 (0.001) | 0.04 | 1.54 | -0.01 (0.006) | -0.02 | -2.55 | 0.073 (0.005) | 0.05 | 2.91 | 1.153 (0.091) | 0.40 | 0.54 | 0.073 (0.005) | 0.05 | 2.91 | 1.153 (0.091) | 0.40 | 0.54 |
| 0.05 | 0.75 | 0.068 (0.001) | 0.02 | 0.36 | -0.028 (0.01) | -0.04 | -3.26 | 0.096 (0.009) | 0.06 | 1.57 | 1.416 (0.143) | 0.67 | 0.89 | 0.096 (0.009) | 0.06 | 1.57 | 1.416 (0.143) | 0.67 | 0.89 |

sTable 7: Estimated effect sizes and size of bias for simulated effect of a phenotypically measured continuous mediator explaining the effect between a continuous exposure and common binary outcome on the risk difference scale (Simulated N=5000)

| True total effect | True proportion mediated | Total effect (SD) | Size of bias (absolute) | Size of bias (relative) | Direct effect (SD) | Size of bias (absolute) | Size of bias (relative) | Indirect effect - difference method (SD) | Size of bias (absolute) | Size of bias (relative) | Proportion mediated - difference (SD) | Size of bias (absolute) | Size of bias (relative) | Indirect effect - product of coefficients (SD) | Size of bias (absolute) | Size of bias (relative) | Proportion mediated - product of coefficients (SD) | Size of bias (absolute) | Size of bias (relative) |
|---|---|---|---|---|---|---|---|---|---|---|---|---|---|---|---|---|---|---|---|
| 0.125 | 0 | 0.196 (0.002) | 0.07 | 0.57 | 0.148 (0.003) | 0.02 | 0.19 | 0.048 (0.002) | 0.05 | | 0.243 (0.011) | 0.24 | | 0.048 (0.002) | 0.05 | | 0.243 (0.011) | 0.24 | |
| 0.125 | -0.5 | 0.196 (0.002) | 0.07 | 0.57 | 0.267 (0.003) | 0.08 | 0.43 | -0.071 (0.003) | -0.26 | 4.14 | -0.364 (0.018) | 0.14 | -0.27 | -0.071 (0.003) | -0.26 | 4.14 | -0.364 (0.018) | 0.14 | -0.27 |
| 0 | 0.05 | 0.157 (0.003) | 0.16 | | 0.087 (0.004) | 0.09 | | 0.07 (0.002) | 0.07 | | 0.445 (0.017) | 0.40 | 7.91 | 0.07 (0.002) | 0.07 | | 0.445 (0.017) | 0.40 | 7.91 |
| 0.05 | 0.05 | 0.178 (0.003) | 0.13 | 2.56 | 0.113 (0.004) | 0.07 | 1.38 | 0.065 (0.002) | 0.06 | 25.09 | 0.366 (0.014) | 0.32 | 6.33 | 0.065 (0.002) | 0.06 | 25.09 | 0.366 (0.014) | 0.32 | 6.33 |
| 0.125 | 0.05 | 0.196 (0.002) | 0.07 | 0.57 | 0.137 (0.004) | 0.02 | 0.15 | 0.059 (0.002) | 0.05 | 8.49 | 0.303 (0.014) | 0.25 | 5.05 | 0.059 (0.002) | 0.05 | 8.49 | 0.303 (0.014) | 0.25 | 5.05 |
| 0.25 | 0.05 | 0.21 (0.002) | -0.04 | -0.16 | 0.157 (0.004) | -0.08 | -0.34 | 0.052 (0.003) | 0.04 | 3.19 | 0.25 (0.013) | 0.20 | 4.00 | 0.052 (0.003) | 0.04 | 3.19 | 0.25 (0.013) | 0.20 | 4.00 |
| 0 | 0.25 | 0.157 (0.003) | 0.16 | | 0.087 (0.004) | 0.09 | | 0.07 (0.002) | 0.07 | | 0.444 (0.017) | 0.19 | 0.78 | 0.07 (0.002) | 0.07 | | 0.444 (0.017) | 0.19 | 0.78 |
| 0.05 | 0.25 | 0.178 (0.003) | 0.13 | 2.56 | 0.089 (0.004) | 0.05 | 1.37 | 0.089 (0.003) | 0.08 | 6.12 | 0.5 (0.018) | 0.25 | 1.00 | 0.089 (0.003) | 0.08 | 6.12 | 0.5 (0.018) | 0.25 | 1.00 |
| 0.125 | 0.25 | 0.196 (0.002) | 0.07 | 0.57 | 0.089 (0.005) | 0.00 | -0.05 | 0.107 (0.004) | 0.08 | 2.42 | 0.545 (0.023) | 0.29 | 1.18 | 0.107 (0.004) | 0.08 | 2.42 | 0.545 (0.023) | 0.29 | 1.18 |
| 0.25 | 0.25 | 0.21 (0.002) | -0.04 | -0.16 | 0.087 (0.007) | -0.10 | -0.53 | 0.122 (0.006) | 0.06 | 0.96 | 0.583 (0.03) | 0.33 | 1.33 | 0.122 (0.006) | 0.06 | 0.96 | 0.583 (0.03) | 0.33 | 1.33 |
| 0 | 0.75 | 0.157 (0.003) | 0.16 | | 0.087 (0.004) | 0.09 | | 0.07 (0.002) | 0.07 | | 0.444 (0.016) | -0.31 | -0.41 | 0.07 (0.002) | 0.07 | | 0.444 (0.016) | -0.31 | -0.41 |
| 0.05 | 0.75 | 0.178 (0.003) | 0.13 | 2.56 | 0.03 (0.006) | 0.02 | 1.36 | 0.148 (0.004) | 0.11 | 2.96 | 0.834 (0.03) | 0.08 | 0.11 | 0.148 (0.004) | 0.11 | 2.96 | 0.834 (0.03) | 0.08 | 0.11 |
| 0.125 | 0.75 | 0.196 (0.002) | 0.07 | 0.57 | -0.029 (0.009) | -0.06 | -1.94 | 0.225 (0.008) | 0.13 | 1.40 | 1.149 (0.047) | 0.40 | 0.53 | 0.225 (0.008) | 0.13 | 1.40 | 1.149 (0.047) | 0.40 | 0.53 |
| 0.25 | 0.75 | 0.21 (0.002) | -0.04 | -0.16 | -0.087 (0.015) | -0.15 | -2.39 | 0.297 (0.014) | 0.11 | 0.58 | 1.415 (0.073) | 0.66 | 0.89 | 0.297 (0.014) | 0.11 | 0.58 | 1.415 (0.073) | 0.66 | 0.89 |

sTable 8: Estimated effect sizes and size of bias for simulated effect of a phenotypically measured continuous mediator explaining the effect between a continuous exposure and rare binary outcome on the risk difference scale, where simulated total effects are small (Simulated N=5000)

| True total effect | True proportion mediated | Total effect from univariate Mendelian randomisation (SD) | Size of bias (absolute) | Size of bias (relative) | Direct effect from multivariable Mendelian randomisation (SD) | Size of bias (absolute) | Size of bias (relative) | Indirect effect from multivariable Mendelian randomisation (SD) | Size of bias (absolute) | Size of bias (relative) | Proportion mediated from multivariable Mendelian randomisation (SD) | Size of bias (absolute) | Size of bias (relative) | Indirect effect from two-step Mendelian randomisation (SD) | Size of bias (absolute) | Size of bias (relative) | Proportion mediated from two-step Mendelian randomisation SD) | Size of bias (absolute) | Size of bias (relative) |
|---|---|---|---|---|---|---|---|---|---|---|---|---|---|---|---|---|---|---|---|
| 0.0005 | 0.05 | 0.051 (0.002) | 0.05 | 101.47 | 0.029 (0.002) | 0.03 | 59.32 | 0.023 (0.001) | 0.02 | 884.28 | 0.441 (0.029) | 0.39 | 7.83 | 0.023 (0.001) | 0.02 | 884.28 | 0.441 (0.029) | 0.39 | 7.83 |
| 0.0025 | 0.05 | 0.053 (0.002) | 0.05 | 20.17 | 0.031 (0.002) | 0.03 | 11.90 | 0.022 (0.001) | 0.02 | 159.28 | 0.422 (0.027) | 0.37 | 7.43 | 0.022 (0.001) | 0.02 | 159.28 | 0.422 (0.027) | 0.37 | 7.43 |
| 0.0005 | 0.05 | 0.055 (0.002) | 0.05 | 108.35 | 0.033 (0.002) | 0.03 | 68.00 | 0.022 (0.001) | 0.02 | 857.04 | 0.401 (0.028) | 0.35 | 7.02 | 0.022 (0.001) | 0.02 | 857.04 | 0.401 (0.028) | 0.35 | 7.02 |
| 0.0005 | 0.25 | 0.051 (0.002) | 0.05 | 101.63 | 0.028 (0.002) | 0.03 | 74.49 | 0.023 (0.001) | 0.02 | 181.05 | 0.449 (0.029) | 0.20 | 0.80 | 0.023 (0.001) | 0.02 | 181.05 | 0.449 (0.029) | 0.20 | 0.80 |
| 0.0025 | 0.25 | 0.053 (0.002) | 0.05 | 20.17 | 0.028 (0.002) | 0.03 | 14.19 | 0.024 (0.001) | 0.02 | 36.10 | 0.462 (0.03) | 0.21 | 0.85 | 0.024 (0.001) | 0.02 | 36.10 | 0.462 (0.03) | 0.21 | 0.85 |
| 0.0005 | 0.25 | 0.055 (0.001) | 0.05 | 108.41 | 0.029 (0.002) | 0.03 | 75.36 | 0.026 (0.001) | 0.03 | 205.57 | 0.477 (0.031) | 0.23 | 0.91 | 0.026 (0.001) | 0.03 | 205.57 | 0.477 (0.031) | 0.23 | 0.91 |
| 0.0005 | 0.75 | 0.051 (0.002) | 0.05 | 101.74 | 0.027 (0.002) | 0.03 | 217.47 | 0.024 (0.001) | 0.02 | 63.83 | 0.469 (0.031) | -0.28 | -0.37 | 0.024 (0.001) | 0.02 | 63.83 | 0.469 (0.031) | -0.28 | -0.37 |
| 0.0025 | 0.75 | 0.053 (0.002) | 0.05 | 20.14 | 0.023 (0.003) | 0.02 | 35.87 | 0.03 (0.002) | 0.03 | 15.57 | 0.565 (0.038) | -0.19 | -0.25 | 0.03 (0.002) | 0.03 | 15.57 | 0.565 (0.038) | -0.19 | -0.25 |
| 0.0005 | 0.75 | 0.055 (0.001) | 0.05 | 108.28 | 0.018 (0.003) | 0.02 | 143.66 | 0.037 (0.002) | 0.04 | 97.15 | 0.67 (0.042) | -0.08 | -0.11 | 0.037 (0.002) | 0.04 | 97.15 | 0.67 (0.042) | -0.08 | -0.11 |

sTable 9: Estimated effect sizes and size of bias for simulated effect of a phenotypically measured continuous mediator explaining the effect between a continuous exposure and common binary outcome on the risk difference scale, where true total effects are small (Simulated N=5000)

| True total effect | True proportion mediated | Total effect from univariate Mendelian randomisation (SD) | Size of bias (absolute) | Size of bias (relative) | Direct effect from multivariable Mendelian randomisation (SD) | Size of bias (absolute) | Size of bias (relative) | Indirect effect from multivariable Mendelian randomisation (SD) | Size of bias (absolute) | Size of bias (relative) | Proportion mediated from multivariable Mendelian randomisation (SD) | Size of bias (absolute) | Size of bias (relative) | Indirect effect from two-step Mendelian randomisation (SD) | Size of bias (absolute) | Size of bias (relative) | Proportion mediated from two-step Mendelian randomisation SD) | Size of bias (absolute) | Size of bias (relative) |
|---|---|---|---|---|---|---|---|---|---|---|---|---|---|---|---|---|---|---|---|
| 0.0025 | 0.05 | 0.158 (0.003) | 0.16 | 62.22 | 0.089 (0.004) | 0.09 | 36.27 | 0.07 (0.002) | 0.07 | 537.24 | 0.44 (0.017) | 0.39 | 7.80 | 0.07 (0.002) | 0.07 | 537.24 | 0.44 (0.017) | 0.39 | 7.80 |
| 0.0125 | 0.05 | 0.163 (0.003) | 0.15 | 12.05 | 0.095 (0.004) | 0.08 | 6.96 | 0.069 (0.002) | 0.06 | 90.70 | 0.421 (0.016) | 0.37 | 7.41 | 0.069 (0.002) | 0.06 | 90.70 | 0.421 (0.016) | 0.37 | 7.41 |
| 0.025 | 0.05 | 0.169 (0.003) | 0.14 | 5.74 | 0.101 (0.004) | 0.08 | 3.26 | 0.067 (0.002) | 0.04 | 34.97 | 0.4 (0.015) | 0.35 | 7.01 | 0.067 (0.002) | 0.04 | 34.97 | 0.4 (0.015) | 0.35 | 7.01 |
| 0.0025 | 0.25 | 0.158 (0.003) | 0.16 | 62.22 | 0.087 (0.004) | 0.09 | 45.56 | 0.071 (0.002) | 0.07 | 110.19 | 0.448 (0.016) | 0.20 | 0.79 | 0.071 (0.002) | 0.07 | 110.19 | 0.448 (0.016) | 0.20 | 0.79 |
| 0.0125 | 0.25 | 0.163 (0.003) | 0.15 | 12.06 | 0.088 (0.004) | 0.08 | 8.38 | 0.075 (0.002) | 0.07 | 21.08 | 0.461 (0.017) | 0.21 | 0.85 | 0.075 (0.002) | 0.07 | 21.08 | 0.461 (0.017) | 0.21 | 0.85 |
| 0.025 | 0.25 | 0.169 (0.003) | 0.14 | 5.75 | 0.088 (0.004) | 0.07 | 3.72 | 0.08 (0.002) | 0.06 | 9.84 | 0.476 (0.017) | 0.23 | 0.90 | 0.08 (0.002) | 0.06 | 9.84 | 0.476 (0.017) | 0.23 | 0.90 |
| 0.0025 | 0.75 | 0.158 (0.003) | 0.16 | 62.22 | 0.084 (0.004) | 0.08 | 133.04 | 0.074 (0.002) | 0.07 | 39.28 | 0.47 (0.017) | -0.28 | -0.37 | 0.074 (0.002) | 0.07 | 39.28 | 0.47 (0.017) | -0.28 | -0.37 |
| 0.0125 | 0.75 | 0.163 (0.003) | 0.15 | 12.05 | 0.071 (0.004) | 0.07 | 21.75 | 0.092 (0.003) | 0.09 | 9.48 | 0.564 (0.02) | -0.19 | -0.25 | 0.092 (0.003) | 0.09 | 9.48 | 0.564 (0.02) | -0.19 | -0.25 |
| 0.025 | 0.75 | 0.169 (0.003) | 0.14 | 5.75 | 0.056 (0.005) | 0.05 | 8.01 | 0.112 (0.003) | 0.11 | 5.66 | 0.666 (0.023) | -0.08 | -0.11 | 0.112 (0.003) | 0.11 | 5.66 | 0.666 (0.023) | -0.08 | -0.11 |

sTable 10: Estimated effect sizes and size of bias for simulated effect of a continuous mediator explaining the effect between a continuous exposure and a rare binary outcome on the risk difference scale using Mendelian randomization (Simulated N=5000)

| True total effect | True proportion mediated | Total effect from univariate Mendelian randomisation (SD) | Size of bias (absolute) | Size of bias (relative) | Direct effect from multivariable Mendelian randomisation (SD) | Size of bias (absolute) | Size of bias (relative) | Indirect effect from multivariable Mendelian randomisation (SD) | Size of bias (absolute) | Size of bias (relative) | Proportion mediated from multivariable Mendelian randomisation (SD) | Size of bias (absolute) | Size of bias (relative) | Indirect effect from two-step Mendelian randomisation (SD) | Size of bias (absolute) | Size of bias (relative) | Proportion mediated from two-step Mendelian randomisation SD) | Size of bias (absolute) | Size of bias (relative) |
|---|---|---|---|---|---|---|---|---|---|---|---|---|---|---|---|---|---|---|---|
| 0.025 | 0 | 0.029 (0.003) | 0.00 | 0.15 | 0.029 (0.003) | 0.00 | 0.15 | 0 (0) | 0.00 | | -0.001 (0.008) | 0.00 | | 0 (0) | 0.00 | | 0 (-0.001) | 0.00 | |
| 0.025 | -0.5 | 0.029 (0.003) | 0.00 | 0.15 | 0.043 (0.004) | 0.01 | 0.15 | -0.014 (0.003) | 0.00 | 0.16 | -0.509 (0.133) | -0.01 | 0.02 | -0.014 (0.003) | 0.00 | 0.16 | -0.509 (0.133) | -0.01 | 0.02 |
| 0 | 0.05 | 0 (0.003) | 0.00 | | 0 (0.003) | 0.00 | | 0 (0) | 0.00 | | -0.038 (2.344) | -0.09 | -1.77 | 0 (0) | 0.00 | | 0 (-0.038) | -0.09 | -1.77 |
| 0.1 | 0.05 | 0.014 (0.003) | -0.09 | -0.86 | 0.014 (0.003) | -0.08 | -0.86 | 0.001 (0) | 0.00 | -0.85 | 0.051 (0.023) | 0.00 | 0.02 | 0.001 (0) | 0.00 | -0.85 | 0 (0.051) | 0.00 | 0.02 |
| 0.025 | 0.05 | 0.029 (0.003) | 0.00 | 0.15 | 0.027 (0.003) | 0.00 | 0.15 | 0.001 (0) | 0.00 | 0.15 | 0.05 (0.015) | 0.00 | 0.01 | 0.001 (0) | 0.00 | 0.15 | 0 (0.05) | 0.00 | 0.01 |
| 0.05 | 0.05 | 0.043 (0.003) | -0.01 | -0.15 | 0.04 (0.003) | -0.01 | -0.15 | 0.002 (0.001) | 0.00 | -0.14 | 0.051 (0.017) | 0.00 | 0.02 | 0.002 (0.001) | 0.00 | -0.14 | 0.001 (0.051) | 0.00 | 0.02 |
| 0 | 0.25 | 0 (0.003) | 0.00 | | 0 (0.003) | 0.00 | | 0 (0) | 0.00 | | -0.08 (3.313) | -0.33 | -1.32 | 0 (0) | 0.00 | | 0 (-0.08) | -0.33 | -1.32 |
| 0.1 | 0.25 | 0.014 (0.003) | -0.09 | -0.43 | 0.011 (0.003) | -0.06 | -0.86 | 0.004 (0.001) | -0.02 | -0.86 | 0.26 (0.078) | 0.01 | 0.04 | 0.004 (0.001) | -0.02 | -0.86 | 0.001 (0.26) | 0.01 | 0.04 |
| 0.025 | 0.25 | 0.029 (0.003) | 0.00 | 0.16 | 0.022 (0.003) | 0.00 | 0.17 | 0.007 (0.002) | 0.00 | 0.15 | 0.25 (0.066) | 0.00 | 0.00 | 0.007 (0.002) | 0.00 | 0.15 | 0.002 (0.25) | 0.00 | 0.00 |
| 0.05 | 0.25 | 0.043 (0.003) | -0.01 | -0.15 | 0.032 (0.004) | -0.01 | -0.14 | 0.011 (0.004) | 0.00 | -0.16 | 0.248 (0.085) | 0.00 | -0.01 | 0.011 (0.004) | 0.00 | -0.16 | 0.004 (0.248) | 0.00 | -0.01 |
| 0 | 0.75 | 0 (0.003) | 0.00 | | 0 (0.003) | 0.00 | | 0 (0) | 0.00 | | 0.027 (2.198) | -0.72 | -0.96 | 0 (0) | 0.00 | | 0 (0.027) | -0.72 | -0.96 |
| 0.1 | 0.75 | 0.014 (0.003) | -0.09 | -0.43 | 0.004 (0.004) | -0.02 | -0.85 | 0.011 (0.002) | -0.06 | -0.86 | 0.782 (0.237) | 0.03 | 0.04 | 0.011 (0.002) | -0.06 | -0.86 | 0.002 (0.782) | 0.03 | 0.04 |
| 0.025 | 0.75 | 0.029 (0.003) | 0.00 | 0.16 | 0.007 (0.006) | 0.00 | 0.16 | 0.022 (0.005) | 0.00 | 0.16 | 0.757 (0.202) | 0.01 | 0.01 | 0.022 (0.005) | 0.00 | 0.16 | 0.005 (0.757) | 0.01 | 0.01 |
| 0.05 | 0.75 | 0.043 (0.003) | -0.01 | -0.15 | 0.011 (0.011) | 0.00 | -0.15 | 0.032 (0.01) | -0.01 | -0.15 | 0.753 (0.247) | 0.00 | 0.00 | 0.032 (0.01) | -0.01 | -0.15 | 0.01 (0.753) | 0.00 | 0.00 |

sTable 11: Estimated effect sizes and size of bias for simulated effect of a continuous mediator explaining the effect between a continuous exposure and a common binary outcome on the risk difference scale using Mendelian randomization (Simulated N=5000)

| True total effect | True proportion mediated | Total effect from univariate Mendelian randomisation (SD) | Size of bias (absolute) | Size of bias (relative) | Direct effect from multivariable Mendelian randomisation (SD) | Size of bias (absolute) | Size of bias (relative) | Indirect effect from multivariable Mendelian randomisation (SD) | Size of bias (absolute) | Size of bias (relative) | Proportion mediated from multivariable Mendelian randomisation (SD) | Size of bias (absolute) | Size of bias (relative) | Indirect effect from two-step Mendelian randomisation (SD) | Size of bias (absolute) | Size of bias (relative) | Proportion mediated from two-step Mendelian randomisation SD) | Size of bias (absolute) | Size of bias (relative) |
|---|---|---|---|---|---|---|---|---|---|---|---|---|---|---|---|---|---|---|---|
| 0.125 | 0 | 0.089 (0.005) | -0.04 | -0.29 | 0.089 (0.005) | -0.04 | -0.29 | 0 (0.001) | 0.00 | | -0.001 (0.008) | 0.00 | | 0 (0.001) | 0.00 | | -0.001 (0.008) | 0.00 | |
| 0.125 | -0.5 | 0.089 (0.005) | -0.04 | -0.29 | 0.133 (0.007) | -0.05 | -0.29 | -0.045 (0.006) | 0.02 | -0.29 | -0.503 (0.079) | 0.00 | 0.01 | -0.045 (0.006) | 0.02 | -0.29 | -0.503 (0.079) | 0.00 | 0.01 |
| 0 | 0.05 | 0 (0.006) | 0.00 | | 0 (0.006) | 0.00 | | 0 (0.001) | 0.00 | | -1.207 (38.778) | -1.26 | -25.14 | 0 (0.001) | 0.00 | | -1.207 (38.778) | -1.26 | -25.14 |
| 0.05 | 0.05 | 0.045 (0.006) | -0.01 | -0.10 | 0.043 (0.005) | 0.00 | -0.10 | 0.002 (0.001) | 0.00 | -0.10 | 0.05 (0.019) | 0.00 | -0.01 | 0.002 (0.001) | 0.00 | -0.10 | 0.05 (0.019) | 0.00 | -0.01 |
| 0.125 | 0.05 | 0.089 (0.005) | -0.04 | -0.29 | 0.085 (0.005) | -0.03 | -0.29 | 0.004 (0.001) | 0.00 | -0.29 | 0.05 (0.01) | 0.00 | -0.01 | 0.004 (0.001) | 0.00 | -0.29 | 0.05 (0.01) | 0.00 | -0.01 |
| 0.25 | 0.05 | 0.131 (0.004) | -0.12 | -0.48 | 0.125 (0.004) | -0.11 | -0.48 | 0.007 (0.001) | -0.01 | -0.48 | 0.05 (0.009) | 0.00 | 0.00 | 0.007 (0.001) | -0.01 | -0.48 | 0.05 (0.009) | 0.00 | 0.00 |
| 0 | 0.25 | 0 (0.006) | 0.00 | | 0 (0.006) | 0.00 | | 0 (0.001) | 0.00 | | 1.384 (49.314) | 1.13 | 4.54 | 0 (0.001) | 0.00 | | 1.384 (49.314) | 1.13 | 4.54 |
| 0.05 | 0.25 | 0.044 (0.005) | -0.01 | -0.12 | 0.033 (0.005) | 0.00 | -0.12 | 0.011 (0.002) | 0.00 | -0.12 | 0.252 (0.04) | 0.00 | 0.01 | 0.011 (0.002) | 0.00 | -0.12 | 0.252 (0.04) | 0.00 | 0.01 |
| 0.125 | 0.25 | 0.089 (0.005) | -0.04 | -0.29 | 0.067 (0.006) | -0.03 | -0.28 | 0.022 (0.003) | -0.01 | -0.29 | 0.248 (0.037) | 0.00 | -0.01 | 0.022 (0.003) | -0.01 | -0.29 | 0.248 (0.037) | 0.00 | -0.01 |
| 0.25 | 0.25 | 0.131 (0.004) | -0.12 | -0.48 | 0.098 (0.007) | -0.09 | -0.48 | 0.033 (0.006) | -0.03 | -0.47 | 0.25 (0.044) | 0.00 | 0.00 | 0.033 (0.006) | -0.03 | -0.47 | 0.25 (0.044) | 0.00 | 0.00 |
| 0 | 0.75 | 0 (0.006) | 0.00 | | 0 (0.005) | 0.00 | | 0 (0.001) | 0.00 | | -0.065 (9.881) | -0.81 | -1.09 | 0 (0.001) | 0.00 | | -0.065 (9.881) | -0.81 | -1.09 |
| 0.05 | 0.75 | 0.044 (0.006) | -0.01 | -0.11 | 0.011 (0.006) | 0.00 | -0.12 | 0.033 (0.004) | 0.00 | -0.11 | 0.764 (0.13) | 0.01 | 0.02 | 0.033 (0.004) | 0.00 | -0.11 | 0.764 (0.13) | 0.01 | 0.02 |
| 0.125 | 0.75 | 0.089 (0.005) | -0.04 | -0.29 | 0.023 (0.01) | -0.01 | -0.26 | 0.066 (0.009) | -0.03 | -0.29 | 0.744 (0.109) | -0.01 | -0.01 | 0.066 (0.009) | -0.03 | -0.29 | 0.744 (0.109) | -0.01 | -0.01 |
| 0.25 | 0.75 | 0.131 (0.004) | -0.12 | -0.48 | 0.033 (0.017) | -0.03 | -0.47 | 0.098 (0.017) | -0.09 | -0.48 | 0.748 (0.131) | 0.00 | 0.00 | 0.098 (0.017) | -0.09 | -0.48 | 0.748 (0.131) | 0.00 | 0.00 |

sTable 12: Estimated effect sizes and size of bias for simulated effect of a continuous mediator explaining the effect between a continuous exposure and rare binary outcome on the risk difference scale using Mendelian randomization, where simulated total effects are small (Simulated N=5000)

| True total effect | True proportion mediated | Total effect from univariate Mendelian randomisation (SD) | Size of bias (absolute) | Size of bias (relative) | Direct effect from multivariable Mendelian randomisation (SD) | Size of bias (absolute) | Size of bias (relative) | Indirect effect from multivariable Mendelian randomisation (SD) | Size of bias (absolute) | Size of bias (relative) | Proportion mediated from multivariable Mendelian randomisation (SD) | Size of bias (absolute) | Size of bias (relative) | Indirect effect from two-step Mendelian randomisation (SD) | Size of bias (absolute) | Size of bias (relative) | Proportion mediated from two-step Mendelian randomisation SD) | Size of bias (absolute) | Size of bias (relative) |
|---|---|---|---|---|---|---|---|---|---|---|---|---|---|---|---|---|---|---|---|
| 0.0005 | 0.05 | 0.001 (0.003) | 0.00 | 0.39 | 0.001 (0.003) | 0.00 | 0.38 | 0 (0) | 0.00 | 0.56 | -0.021 (5.936) | -0.07 | -1.42 | 0 (0) | 0.00 | 0.56 | -0.021 (5.936) | -0.07 | -1.42 |
| 0.0025 | 0.05 | 0.004 (0.003) | 0.00 | 0.64 | 0.004 (0.003) | 0.00 | 0.64 | 0 (0) | 0.00 | 0.71 | 0.05 (0.682) | 0.00 | -0.01 | 0 (0) | 0.00 | 0.71 | 0.05 (0.682) | 0.00 | -0.01 |
| 0.0005 | 0.05 | 0.008 (0.003) | 0.01 | 14.51 | 0.007 (0.003) | 0.01 | 14.51 | 0 (0) | 0.00 | 14.48 | 0.05 (0.169) | 0.00 | -0.01 | 0 (0) | 0.00 | 14.48 | 0.05 (0.169) | 0.00 | -0.01 |
| 0.0005 | 0.25 | 0.001 (0.003) | 0.00 | 0.70 | 0.001 (0.003) | 0.00 | 0.71 | 0 (0) | 0.00 | 0.65 | 0.055 (1.544) | -0.20 | -0.78 | 0 (0) | 0.00 | 0.65 | 0.055 (1.544) | -0.20 | -0.78 |
| 0.0025 | 0.25 | 0.004 (0.003) | 0.00 | 0.61 | 0.003 (0.003) | 0.00 | 0.61 | 0.001 (0) | 0.00 | 0.62 | 1.169 (30.416) | 0.92 | 3.68 | 0.001 (0) | 0.00 | 0.62 | 1.169 (30.416) | 0.92 | 3.68 |
| 0.0005 | 0.25 | 0.008 (0.003) | 0.01 | 14.38 | 0.006 (0.003) | 0.01 | 14.35 | 0.002 (0) | 0.00 | 14.48 | 0.275 (0.864) | 0.02 | 0.10 | 0.002 (0) | 0.00 | 14.48 | 0.275 (0.864) | 0.02 | 0.10 |
| 0.0005 | 0.75 | 0.001 (0.003) | 0.00 | 0.96 | 0 (0.003) | 0.00 | 1.86 | 0.001 (0) | 0.00 | 0.66 | -0.061 (7.019) | -0.81 | -1.08 | 0.001 (0) | 0.00 | 0.66 | -0.061 (7.019) | -0.81 | -1.08 |
| 0.0025 | 0.75 | 0.004 (0.003) | 0.00 | 0.64 | 0.001 (0.003) | 0.00 | 0.67 | 0.003 (0.001) | 0.00 | 0.63 | 0.492 (18.682) | -0.26 | -0.34 | 0.003 (0.001) | 0.00 | 0.63 | 0.492 (18.682) | -0.26 | -0.34 |
| 0.0005 | 0.75 | 0.008 (0.003) | 0.01 | 14.34 | 0.002 (0.003) | 0.00 | 13.27 | 0.006 (0.001) | 0.01 | 14.70 | 1.047 (2.14) | 0.30 | 0.40 | 0.006 (0.001) | 0.01 | 14.70 | 1.047 (2.14) | 0.30 | 0.40 |

sTable 13: Estimated effect sizes and size of bias for simulated effect of a continuous mediator explaining the effect between a continuous exposure and common binary outcome on the risk difference scale using Mendelian randomization, where simulated total effects are small (Simulated N=5000)

| True total effect | True proportion mediated | Total effect from univariate Mendelian randomisation (SD) | Size of bias (absolute) | Size of bias (relative) | Direct effect from multivariable Mendelian randomisation (SD) | Size of bias (absolute) | Size of bias (relative) | Indirect effect from multivariable Mendelian randomisation (SD) | Size of bias (absolute) | Size of bias (relative) | Proportion mediated from multivariable Mendelian randomisation (SD) | Size of bias (absolute) | Size of bias (relative) | Indirect effect from two-step Mendelian randomisation (SD) | Size of bias (absolute) | Size of bias (relative) | Proportion mediated from two-step Mendelian randomisation SD) | Size of bias (absolute) | Size of bias (relative) |
|---|---|---|---|---|---|---|---|---|---|---|---|---|---|---|---|---|---|---|---|
| 0.0025 | 0.05 | 0.002 (0.006) | 0.00 | -0.02 | 0.002 (0.006) | 0.00 | -0.02 | 0 (0.001) | 0.00 | -0.05 | 0.115 (2.813) | 0.07 | 1.31 | 0 (0.001) | 0.00 | -0.05 | 0.115 (2.813) | 0.07 | 1.31 |
| 0.0125 | 0.05 | 0.013 (0.006) | 0.00 | 0.02 | 0.012 (0.006) | 0.00 | 0.02 | 0.001 (0.001) | 0.00 | 0.04 | 0.047 (0.533) | 0.00 | -0.06 | 0.001 (0.001) | 0.00 | 0.04 | 0.047 (0.533) | 0.00 | -0.06 |
| 0.025 | 0.05 | 0.024 (0.006) | 0.00 | -0.04 | 0.023 (0.005) | 0.00 | -0.04 | 0.001 (0.001) | 0.00 | -0.05 | 0.046 (0.044) | 0.00 | -0.08 | 0.001 (0.001) | 0.00 | -0.05 | 0.046 (0.044) | 0.00 | -0.08 |
| 0.0025 | 0.25 | 0.003 (0.006) | 0.00 | 0.04 | 0.002 (0.005) | 0.00 | 0.05 | 0.001 (0.001) | 0.00 | 0.01 | 0.243 (5.842) | -0.01 | -0.03 | 0.001 (0.001) | 0.00 | 0.01 | 0.243 (5.842) | -0.01 | -0.03 |
| 0.0125 | 0.25 | 0.013 (0.006) | 0.00 | 0.02 | 0.01 (0.006) | 0.00 | 0.02 | 0.003 (0.001) | 0.00 | 0.00 | 0.287 (1.014) | 0.04 | 0.15 | 0.003 (0.001) | 0.00 | 0.00 | 0.287 (1.014) | 0.04 | 0.15 |
| 0.025 | 0.25 | 0.024 (0.006) | 0.00 | -0.04 | 0.018 (0.005) | 0.00 | -0.04 | 0.006 (0.001) | 0.00 | -0.05 | 0.26 (0.075) | 0.01 | 0.04 | 0.006 (0.001) | 0.00 | -0.05 | 0.26 (0.075) | 0.01 | 0.04 |
| 0.0025 | 0.75 | 0.003 (0.006) | 0.00 | 0.01 | 0.001 (0.006) | 0.00 | -0.03 | 0.002 (0.001) | 0.00 | 0.02 | 1.458 (28.614) | 0.71 | 0.94 | 0.002 (0.001) | 0.00 | 0.02 | 1.458 (28.614) | 0.71 | 0.94 |
| 0.0125 | 0.75 | 0.013 (0.006) | 0.00 | 0.02 | 0.003 (0.006) | 0.00 | 0.06 | 0.009 (0.001) | 0.00 | 0.01 | 0.877 (4.495) | 0.13 | 0.17 | 0.009 (0.001) | 0.00 | 0.01 | 0.877 (4.495) | 0.13 | 0.17 |
| 0.025 | 0.75 | 0.024 (0.006) | 0.00 | -0.04 | 0.006 (0.006) | 0.00 | -0.02 | 0.018 (0.002) | 0.00 | -0.04 | 0.792 (0.231) | 0.04 | 0.06 | 0.018 (0.002) | 0.00 | -0.04 | 0.792 (0.231) | 0.04 | 0.06 |

sTable 14: Estimated effect sizes and size of bias for simulated effect of a continuous mediator explaining the effect between a continuous exposure and a rare binary outcome on the log odds ratio scale using Mendelian randomization (Simulated N=5000)

| True total effect | True proportion mediated | Total effect from univariate Mendelian randomisation (SD) | Size of bias (absolute) | Size of bias (relative) | Direct effect from multivariable Mendelian randomisation (SD) | Size of bias (absolute) | Size of bias (relative) | Indirect effect from multivariable Mendelian randomisation (SD) | Size of bias (absolute) | Size of bias (relative) | Proportion mediated from multivariable Mendelian randomisation (SD) | Size of bias (absolute) | Size of bias (relative) | Indirect effect from two-step Mendelian randomisation (SD) | Size of bias (absolute) | Size of bias (relative) | Proportion mediated from two-step Mendelian randomisation SD) | Size of bias (absolute) | Size of bias (relative) |
|---|---|---|---|---|---|---|---|---|---|---|---|---|---|---|---|---|---|---|---|
| 0.5 | 0 | 0.617 (0.063) | 0.12 | 0.23 | 0.62 (0.062) | 0.12 | 0.24 | -0.003 (0.006) | 0.00 | | -0.004 (0.01) | 0.00 | | 0 (0.005) | 0.00 | | -0.001 (0.008) | 0.00 | |
| 0 | 0.05 | -0.003 (0.063) | 0.00 | | -0.003 (0.062) | 0.00 | | 0 (0.007) | 0.00 | | -0.049 (2.294) | -0.10 | -1.99 | 0 (0.007) | 0.00 | | -0.039 (2.356) | -0.09 | -1.78 |
| 0.2 | 0.05 | 0.306 (0.061) | 0.11 | 0.53 | 0.292 (0.06) | 0.10 | 0.54 | 0.014 (0.007) | 0.00 | 0.39 | 0.046 (0.024) | 0.00 | -0.08 | 0.016 (0.007) | 0.01 | 0.56 | 0.051 (0.023) | 0.00 | 0.03 |
| 0.5 | 0.05 | 0.621 (0.065) | 0.12 | 0.24 | 0.592 (0.064) | 0.12 | 0.25 | 0.028 (0.009) | 0.00 | 0.14 | 0.046 (0.014) | 0.00 | -0.08 | 0.031 (0.01) | 0.01 | 0.25 | 0.051 (0.016) | 0.00 | 0.01 |
| 1 | 0.05 | 0.946 (0.065) | -0.05 | -0.05 | 0.901 (0.066) | -0.05 | -0.05 | 0.045 (0.015) | -0.01 | -0.10 | 0.048 (0.016) | 0.00 | -0.05 | 0.048 (0.016) | 0.00 | -0.04 | 0.051 (0.017) | 0.00 | 0.02 |
| 0 | 0.25 | 0 (0.063) | 0.00 | | 0 (0.061) | 0.00 | | 0 (0.007) | 0.00 | | -0.048 (3.005) | -0.30 | -1.19 | 0 (0.007) | 0.00 | | -0.08 (3.316) | -0.33 | -1.32 |
| 0.2 | 0.25 | 0.305 (0.062) | 0.10 | 0.52 | 0.23 (0.062) | 0.08 | 0.53 | 0.075 (0.016) | 0.02 | 0.50 | 0.256 (0.078) | 0.01 | 0.02 | 0.077 (0.017) | 0.03 | 0.53 | 0.261 (0.079) | 0.01 | 0.05 |
| 0.5 | 0.25 | 0.625 (0.065) | 0.12 | 0.25 | 0.472 (0.073) | 0.10 | 0.26 | 0.153 (0.037) | 0.03 | 0.22 | 0.247 (0.065) | 0.00 | -0.01 | 0.156 (0.038) | 0.03 | 0.25 | 0.251 (0.067) | 0.00 | 0.01 |
| 1 | 0.25 | 0.947 (0.067) | -0.05 | -0.05 | 0.715 (0.102) | -0.04 | -0.05 | 0.232 (0.077) | -0.02 | -0.07 | 0.246 (0.083) | 0.00 | -0.02 | 0.235 (0.078) | -0.02 | -0.06 | 0.249 (0.085) | 0.00 | 0.00 |
| 0 | 0.75 | -0.003 (0.065) | 0.00 | | -0.003 (0.063) | 0.00 | | 0 (0.007) | 0.00 | | 0.042 (2.245) | -0.71 | -0.94 | 0 (0.007) | 0.00 | | 0.028 (2.217) | -0.72 | -0.96 |
| 0.2 | 0.75 | 0.304 (0.063) | 0.10 | 0.52 | 0.077 (0.075) | 0.03 | 0.55 | 0.227 (0.045) | 0.08 | 0.51 | 0.781 (0.237) | 0.03 | 0.04 | 0.229 (0.046) | 0.08 | 0.52 | 0.786 (0.239) | 0.04 | 0.05 |
| 0.5 | 0.75 | 0.622 (0.061) | 0.12 | 0.24 | 0.156 (0.13) | 0.03 | 0.25 | 0.465 (0.115) | 0.09 | 0.24 | 0.756 (0.202) | 0.01 | 0.01 | 0.468 (0.116) | 0.09 | 0.25 | 0.761 (0.204) | 0.01 | 0.01 |
| 1 | 0.75 | 0.945 (0.064) | -0.05 | -0.05 | 0.234 (0.235) | -0.02 | -0.06 | 0.711 (0.227) | -0.04 | -0.05 | 0.756 (0.248) | 0.01 | 0.01 | 0.714 (0.229) | -0.04 | -0.05 | 0.759 (0.25) | 0.01 | 0.01 |

sTable 15: Estimated effect sizes and size of bias for simulated effect of a continuous mediator explaining the effect between a continuous exposure and a common binary outcome on the log odds ratio scale using Mendelian randomization (Simulated N=5000)

| True total effect | True proportion mediated | Total effect from univariate Mendelian randomisation (SD) | Size of bias (absolute) | Size of bias (relative) | Direct effect from multivariable Mendelian randomisation (SD) | Size of bias (absolute) | Size of bias (relative) | Indirect effect from multivariable Mendelian randomisation (SD) | Size of bias (absolute) | Size of bias (relative) | Proportion mediated from multivariable Mendelian randomisation (SD) | Size of bias (absolute) | Size of bias (relative) | Indirect effect from two-step Mendelian randomisation (SD) | Size of bias (absolute) | Size of bias (relative) | Proportion mediated from two-step Mendelian randomisation SD) | Size of bias (absolute) | Size of bias (relative) |
|---|---|---|---|---|---|---|---|---|---|---|---|---|---|---|---|---|---|---|---|
| 0.5 | 0 | 0.496 (0.032) | 0.00 | -0.01 | 0.5 (0.03) | 0.00 | 0.00 | -0.004 (0.004) | 0.00 | | -0.008 (0.009) | -0.01 | -0.02 | 0 (0.004) | 0.00 | | -0.001 (0.008) | 0.00 | 0.00 |
| 0 | 0.05 | -0.001 (0.033) | 0.00 | | -0.001 (0.03) | 0.00 | | 0 (0.005) | 0.00 | | -1.244 (38.267) | -1.29 | | 0 (0.006) | 0.00 | | -1.226 (39.475) | -1.28 | |
| 0.2 | 0.05 | 0.242 (0.031) | 0.04 | 0.21 | 0.232 (0.029) | 0.04 | 0.22 | 0.01 (0.005) | 0.00 | -0.05 | 0.039 (0.019) | -0.01 | -0.06 | 0.012 (0.005) | 0.00 | 0.23 | 0.05 (0.019) | 0.00 | 0.00 |
| 0.5 | 0.05 | 0.497 (0.03) | 0.00 | -0.01 | 0.476 (0.029) | 0.00 | 0.00 | 0.021 (0.005) | 0.00 | -0.16 | 0.042 (0.009) | -0.01 | -0.02 | 0.025 (0.006) | 0.00 | 0.00 | 0.05 (0.01) | 0.00 | 0.00 |
| 1 | 0.05 | 0.775 (0.032) | -0.23 | -0.23 | 0.74 (0.032) | -0.21 | -0.22 | 0.035 (0.006) | -0.01 | -0.30 | 0.045 (0.008) | 0.00 | 0.00 | 0.039 (0.007) | -0.01 | -0.22 | 0.05 (0.009) | 0.00 | 0.00 |
| 0 | 0.25 | 0.001 (0.034) | 0.00 | | 0.001 (0.031) | 0.00 | | 0 (0.006) | 0.00 | | 1.345 (48.216) | 1.10 | | 0 (0.006) | 0.00 | | 1.423 (50.673) | 1.17 | |
| 0.2 | 0.25 | 0.238 (0.03) | 0.04 | 0.19 | 0.181 (0.029) | 0.03 | 0.21 | 0.058 (0.008) | 0.01 | 0.15 | 0.244 (0.039) | -0.01 | -0.04 | 0.06 (0.008) | 0.01 | 0.20 | 0.255 (0.041) | 0.01 | 0.04 |
| 0.5 | 0.25 | 0.498 (0.031) | 0.00 | 0.00 | 0.377 (0.034) | 0.00 | 0.01 | 0.12 (0.017) | 0.00 | -0.04 | 0.243 (0.036) | -0.01 | -0.02 | 0.124 (0.018) | 0.00 | -0.01 | 0.25 (0.038) | 0.00 | 0.00 |
| 1 | 0.25 | 0.775 (0.032) | -0.22 | -0.22 | 0.584 (0.044) | -0.17 | -0.22 | 0.191 (0.033) | -0.06 | -0.23 | 0.247 (0.043) | 0.00 | 0.00 | 0.195 (0.034) | -0.06 | -0.22 | 0.252 (0.045) | 0.00 | 0.00 |
| 0 | 0.75 | 0.001 (0.032) | 0.00 | | 0.001 (0.03) | 0.00 | | 0 (0.005) | 0.00 | | -0.048 (9.389) | -0.80 | | 0 (0.005) | 0.00 | | -0.066 (10.008) | -0.82 | |
| 0.2 | 0.75 | 0.239 (0.031) | 0.04 | 0.19 | 0.06 (0.035) | 0.01 | 0.19 | 0.179 (0.021) | 0.03 | 0.19 | 0.762 (0.131) | 0.01 | 0.24 | 0.182 (0.022) | 0.03 | 0.21 | 0.773 (0.133) | 0.02 | 0.46 |
| 0.5 | 0.75 | 0.497 (0.031) | 0.00 | -0.01 | 0.129 (0.057) | 0.00 | 0.03 | 0.368 (0.05) | -0.01 | -0.02 | 0.743 (0.109) | -0.01 | -0.06 | 0.372 (0.051) | 0.00 | -0.01 | 0.75 (0.111) | 0.00 | 0.00 |
| 1 | 0.75 | 0.774 (0.031) | -0.23 | -0.23 | 0.196 (0.103) | -0.05 | -0.22 | 0.579 (0.099) | -0.17 | -0.23 | 0.748 (0.131) | 0.00 | -0.01 | 0.582 (0.101) | -0.17 | -0.22 | 0.753 (0.133) | 0.00 | 0.01 |

sTable 16: Estimated effect sizes and size of bias for simulated effect of a continuous mediator explaining the effect between a continuous exposure and a rare binary outcome on the odds ratio scale using Mendelian randomization (Simulated N=5000)

| True total effect | True proportion mediated | Total effect from univariate Mendelian randomisation (SD) | Size of bias (absolute) | Size of bias (relative) | Direct effect from multivariable Mendelian randomisation (SD) | Size of bias (absolute) | Size of bias (relative) | Indirect effect from multivariable Mendelian randomisation (SD) | Size of bias (absolute) | Size of bias (relative) | Proportion mediated from multivariable Mendelian randomisation (SD) | Size of bias (absolute) | Size of bias (relative) | Indirect effect from two-step Mendelian randomisation (SD) | Size of bias (absolute) | Size of bias (relative) | Proportion mediated from two-step Mendelian randomisation SD) | Size of bias (absolute) | Size of bias (relative) |
|---|---|---|---|---|---|---|---|---|---|---|---|---|---|---|---|---|---|---|---|
| 1.65 | 0 | 1.857 (0.116) | 0.21 | 0.13 | 1.862 (0.115) | 0.21 | 0.13 | -0.005 (0.011) | 0.00 | | -0.003 (0.006) | 0.00 | | -0.001 (0.025) | 0.00 | | 0.025 (-0.001) | 0.00 | |
| 1.00 | 0.05 | 0.999 (0.063) | 0.00 | 0.00 | 0.999 (0.062) | 0.05 | 0.05 | 0 (0.007) | -0.05 | -1.00 | 0 (0.007) | -0.05 | -1.00 | 0 (0.029) | -0.05 | -1.01 | 0.029 (-0.001) | -0.05 | -1.02 |
| 1.22 | 0.05 | 1.361 (0.084) | 0.14 | 0.11 | 1.342 (0.081) | 0.18 | 0.16 | 0.019 (0.01) | -0.04 | -0.69 | 0.014 (0.007) | -0.04 | -0.72 | 0.069 (0.028) | 0.01 | 0.13 | 0.028 (0.05) | 0.00 | 0.01 |
| 1.65 | 0.05 | 1.864 (0.121) | 0.22 | 0.13 | 1.812 (0.117) | 0.25 | 0.16 | 0.052 (0.017) | -0.03 | -0.36 | 0.028 (0.009) | -0.02 | -0.44 | 0.161 (0.029) | 0.08 | 0.95 | 0.029 (0.086) | 0.04 | 0.72 |
| 2.72 | 0.05 | 2.58 (0.168) | -0.14 | -0.05 | 2.467 (0.163) | -0.12 | -0.04 | 0.113 (0.037) | -0.02 | -0.17 | 0.044 (0.014) | -0.01 | -0.12 | 0.303 (0.031) | 0.17 | 1.23 | 0.031 (0.118) | 0.07 | 1.36 |
| 1.00 | 0.25 | 1.002 (0.063) | 0.00 | 0.00 | 1.002 (0.061) | 0.25 | 0.34 | 0 (0.007) | -0.25 | -1.00 | 0 (0.007) | -0.25 | -1.00 | -0.001 (0.029) | -0.25 | -1.00 | 0.029 (-0.002) | -0.25 | -1.01 |
| 1.22 | 0.25 | 1.359 (0.084) | 0.14 | 0.11 | 1.261 (0.078) | 0.34 | 0.38 | 0.098 (0.022) | -0.21 | -0.68 | 0.072 (0.015) | -0.18 | -0.71 | 0.339 (0.034) | 0.03 | 0.11 | 0.034 (0.25) | 0.00 | 0.00 |
| 1.65 | 0.25 | 1.872 (0.122) | 0.22 | 0.14 | 1.607 (0.117) | 0.37 | 0.30 | 0.265 (0.063) | -0.15 | -0.36 | 0.141 (0.032) | -0.11 | -0.43 | 0.803 (0.055) | 0.39 | 0.95 | 0.055 (0.431) | 0.18 | 0.72 |
| 2.72 | 0.25 | 2.583 (0.173) | -0.14 | -0.05 | 2.055 (0.209) | 0.02 | 0.01 | 0.528 (0.162) | -0.15 | -0.22 | 0.205 (0.061) | -0.05 | -0.18 | 1.512 (0.098) | 0.83 | 1.23 | 0.098 (0.588) | 0.34 | 1.35 |
| 1.00 | 0.75 | 0.999 (0.065) | 0.00 | 0.00 | 0.999 (0.063) | 0.75 | 3.00 | 0 (0.007) | -0.75 | -1.00 | 0 (0.007) | -0.75 | -1.00 | 0 (0.028) | -0.75 | -1.00 | 0.028 (0) | -0.75 | -1.00 |
| 1.22 | 0.75 | 1.359 (0.086) | 0.14 | 0.11 | 1.083 (0.081) | 0.78 | 2.55 | 0.275 (0.053) | -0.64 | -0.70 | 0.202 (0.036) | -0.55 | -0.73 | 1.019 (0.067) | 0.10 | 0.11 | 0.067 (0.752) | 0.00 | 0.00 |
| 1.65 | 0.75 | 1.865 (0.113) | 0.22 | 0.13 | 1.179 (0.156) | 0.77 | 1.86 | 0.686 (0.143) | -0.55 | -0.44 | 0.368 (0.073) | -0.38 | -0.51 | 2.412 (0.151) | 1.18 | 0.95 | 0.151 (1.298) | 0.55 | 0.73 |
| 2.72 | 0.75 | 2.578 (0.165) | -0.14 | -0.05 | 1.299 (0.304) | 0.62 | 0.91 | 1.28 (0.308) | -0.76 | -0.37 | 0.496 (0.115) | -0.25 | -0.34 | 4.545 (0.279) | 2.51 | 1.23 | 0.279 (1.77) | 1.02 | 1.36 |

sTable 17: Estimated effect sizes and size of bias for simulated effect of a continuous mediator explaining the effect between a continuous exposure and a common binary outcome on the odds ratio scale using Mendelian randomization (Simulated N=5000)

| True total effect | True proportion mediated | Total effect from univariate Mendelian randomisation (SD) | Size of bias | Direct effect from multivariable Mendelian randomisation (SD) | Size of bias | Indirect effect from multivariable Mendelian randomisation (SD) | Size of bias | Proportion mediated from multivariable Mendelian randomisation (SD) | Size of bias | Indirect effect from two-step Mendelian randomisation (SD) | Size of bias | Proportion mediated from two-step Mendelian randomisation SD) | Size of bias |
|---|---|---|---|---|---|---|---|---|---|---|---|---|---|
| 1.65 | 0 | 1.643 (0.052) | -0.01 | 1.65 (0.05) | 0.00 | -0.006 (0.007) | -0.01 | -0.004 (0.004) | 0.00 | -0.001 (0.024) | 0.00 | -0.001 (0.015) | 0.00 |
| 1.00 | 0.05 | 1 (0.033) | 0.00 | 1 (0.03) | 0.05 | 0 (0.005) | -0.05 | 0 (0.005) | -0.05 | 0 (0.027) | -0.05 | -0.001 (0.027) | -0.05 |
| 1.22 | 0.05 | 1.274 (0.039) | 0.05 | 1.262 (0.036) | 0.10 | 0.012 (0.006) | -0.05 | 0.009 (0.005) | -0.04 | 0.064 (0.025) | 0.00 | 0.05 (0.019) | 0.00 |
| 1.65 | 0.05 | 1.645 (0.049) | 0.00 | 1.611 (0.046) | 0.04 | 0.034 (0.008) | -0.05 | 0.021 (0.005) | -0.03 | 0.152 (0.026) | 0.07 | 0.093 (0.015) | 0.04 |
| 2.72 | 0.05 | 2.171 (0.07) | -0.55 | 2.096 (0.067) | -0.49 | 0.075 (0.013) | -0.06 | 0.034 (0.006) | -0.02 | 0.292 (0.024) | 0.16 | 0.134 (0.011) | 0.08 |
| 1.00 | 0.25 | 1.001 (0.034) | 0.00 | 1.001 (0.031) | 0.25 | 0 (0.006) | -0.25 | 0 (0.006) | -0.25 | -0.001 (0.027) | -0.25 | -0.001 (0.027) | -0.25 |
| 1.22 | 0.25 | 1.27 (0.038) | 0.05 | 1.199 (0.035) | 0.28 | 0.071 (0.01) | -0.23 | 0.056 (0.007) | -0.19 | 0.317 (0.027) | 0.01 | 0.249 (0.019) | 0.00 |
| 1.65 | 0.25 | 1.646 (0.051) | 0.00 | 1.459 (0.05) | 0.69 | 0.187 (0.025) | -0.07 | 0.113 (0.015) | -0.14 | 0.763 (0.032) | 0.51 | 0.464 (0.02) | 0.21 |
| 2.72 | 0.25 | 2.173 (0.069) | -0.55 | 1.795 (0.08) | 1.01 | 0.377 (0.06) | 0.11 | 0.174 (0.027) | -0.08 | 1.463 (0.046) | 1.20 | 0.674 (0.027) | 0.42 |
| 1.00 | 0.75 | 1.001 (0.032) | 0.00 | 1.001 (0.03) | 0.75 | 0 (0.005) | -0.75 | 0 (0.005) | -0.75 | 0 (0.026) | -0.75 | 0 (0.026) | -0.75 |
| 1.22 | 0.75 | 1.27 (0.039) | 0.05 | 1.062 (0.038) | 0.76 | 0.208 (0.024) | -0.71 | 0.164 (0.018) | -0.59 | 0.956 (0.037) | 0.04 | 0.753 (0.029) | 0.00 |
| 1.65 | 0.75 | 1.645 (0.051) | 0.00 | 1.14 (0.066) | 0.88 | 0.505 (0.059) | -0.26 | 0.307 (0.035) | -0.44 | 2.288 (0.067) | 1.52 | 1.392 (0.053) | 0.64 |
| 2.72 | 0.75 | 2.170 (0.068) | -0.55 | 1.222 (0.126) | 0.96 | 0.948 (0.126) | 0.16 | 0.437 (0.056) | -0.31 | 4.382 (0.12) | 3.59 | 2.021 (0.08) | 1.27 |

sTable 18: Estimated effect sizes and size of bias for simulated effect of a phenotypically measured continuous mediator explaining the effect between a continuous exposure and continuous outcome (per unit increase in exposure), and a rare binary outcome and common binary outcome on the risk difference scale, where measurement error is introduced in either the exposure or mediator (Simulated N=5000)

| | **Outcome** | **True total effect** | **True proportion mediated** | **Total effect (SD)** | **Size of bias (absolute)** | **Size of bias (relative)** | **Direct effect (SD)** | **Size of bias (absolute)** | **Size of bias (relative)** | **Indirect effect - difference method (SD)** | **Size of bias (absolute)** | **Size of bias (relative)** | **Proportion mediated - difference (SD)** | **Size of bias (absolute)** | **Size of bias (relative)** | **Indirect effect - product of coefficients (SD)** | **Size of bias (absolute)** | **Size of bias (relative)** | **Proportion mediated - product of coefficients (SD)** | **Size of bias (absolute)** | **Size of bias (relative)** |
|---|---|---|---|---|---|---|---|---|---|---|---|---|---|---|---|---|---|---|---|---|---|
| **Measurement error in the exposure** | Continuous | 0.5 | 0.25 | 0.366 (0.009) | 0.17 | 0.33 | 0.078 (0.004) | -0.30 | -0.59 | 0.288 (0.008) | 0.16 | 1.30 | 0.786 (0.011) | 0.54 | 2.14 | 0.288 (0.008) | 0.16 | 1.30 | 0.786 (0.011) | 0.54 | 0.27 |
| | Rare binary | 0.025 | 0.25 | 0.021 (0.001) | 0.00 | -0.15 | 0.005 (0.001) | -0.01 | -0.76 | 0.017 (0.001) | 0.01 | 1.66 | 0.787 (0.049) | 0.54 | 2.15 | 0.017 (0.001) | 0.01 | 1.66 | 0.787 (0.049) | 0.54 | 2.15 |
| | Common binary | 0.125 | 0.25 | 0.065 (0.002) | -0.06 | -0.48 | 0.014 (0.002) | -0.08 | -0.85 | 0.051 (0.001) | 0.02 | 0.64 | 0.785 (0.027) | 0.54 | 2.14 | 0.051 (0.001) | 0.02 | 0.64 | 0.785 (0.027) | 0.54 | 2.14 |
| **Measurement error in the mediator** | Continuous | 0.5 | 0.25 | 1.1 (0.009) | 0.60 | 1.20 | 0.936 (0.01) | 0.56 | 1.12 | 0.164 (0.006) | 0.04 | 0.31 | 0.149 (0.006) | -0.10 | -0.40 | 0.164 (0.006) | 0.04 | 0.31 | 0.149 (0.006) | -0.10 | -0.05 |
| | Rare binary | 0.025 | 0.25 | 0.064 (0.001) | 0.04 | 1.54 | 0.054 (0.002) | 0.04 | 1.89 | 0.009 (0.001) | 0.00 | 0.51 | 0.148 (0.021) | -0.10 | -0.41 | 0.009 (0.001) | 0.00 | 0.51 | 0.148 (0.021) | -0.10 | -0.41 |
| | Common binary | 0.125 | 0.25 | 0.196 (0.002) | 0.07 | 0.57 | 0.167 (0.004) | 0.07 | 0.78 | 0.029 (0.002) | 0.00 | -0.07 | 0.148 (0.012) | -0.10 | -0.41 | 0.029 (0.002) | 0.00 | -0.07 | 0.148 (0.012) | -0.10 | -0.41 |

sTable 19: Estimated effect sizes and size of bias for simulated effect of a phenotypically measured continuous mediator explaining the effect between a continuous exposure and continuous outcome (per unit increase in exposure), and a rare binary outcome and common binary outcome on the risk difference scale using Mendelian Randomization, where measurement error is introduced in either the exposure or mediator (Simulated N=5000)

| | Outcome | True total effect | True proportion mediated | Total effect from uni-variate Mendelian rando-misation (SD) | Size of bias (absolute) | Size of bias (relative) | Direct effect from multi-variable Mendelian randomisation (SD) | Size of bias (absolute) | Size of bias (relative) | Indirect effect from multi-variable Mendelian random-isation (SD) | Size of bias (absolute) | Size of bias (relative) | Proportion mediated from multi-variable Mendelian rando-misation (SD) | Size of bias (absolute) | Size of bias (relative) | Indirect effect from two-step Mendelian random-isation (SD) | Size of bias (absolute) | Size of bias (relative) | Proportion mediated from two-step Mendelian random-isation SD) | Size of bias (absolute) | Size of bias (relative) |
|---|---|---|---|---|---|---|---|---|---|---|---|---|---|---|---|---|---|---|---|---|---|
| **Measurement error in the exposure** | Continuous | 0.5 | 0.25 | 0.499 (0.022) | 0.00 | 0.00 | 0.375 (0.021) | 0.00 | 0.00 | 0.124 (0.013) | 0.00 | -0.01 | 0.249 (0.024) | 0.00 | 0.00 | 0.124 (0.013) | 0.00 | -0.01 | 0.013 (0.249) | 0.00 | 0.00 |
| | Rare binary | 0.025 | 0.25 | 0.029 (0.003) | 0.00 | 0.16 | 0.022 (0.003) | 0.00 | 0.16 | 0.007 (0.002) | 0.00 | 0.15 | 0.252 (0.067) | 0.00 | 0.01 | 0.007 (0.002) | 0.00 | 0.15 | 0.002 (0.252) | 0.00 | 0.01 |
| | Common binary | 0.125 | 0.25 | 0.089 (0.006) | -0.04 | -0.29 | 0.067 (0.006) | -0.03 | -0.29 | 0.022 (0.003) | -0.01 | -0.29 | 0.248 (0.039) | 0.00 | -0.01 | 0.022 (0.003) | -0.01 | -0.29 | 0.248 (0.039) | 0.00 | -0.01 |
| **Measurement error in the mediator** | Continuous | 0.5 | 0.25 | 0.499 (0.017) | 0.00 | 0.00 | 0.374 (0.018) | 0.00 | 0.00 | 0.125 (0.012) | 0.00 | 0.00 | 0.251 (0.022) | 0.00 | 0.00 | 0.125 (0.012) | 0.00 | 0.00 | 0.012 (0.251) | 0.00 | 0.00 |
| | Rare binary | 0.025 | 0.25 | 0.029 (0.003) | 0.00 | 0.15 | 0.022 (0.003) | 0.00 | 0.15 | 0.007 (0.002) | 0.00 | 0.16 | 0.255 (0.068) | 0.00 | 0.02 | 0.007 (0.002) | 0.00 | 0.16 | 0.002 (0.255) | 0.00 | 0.02 |
| | Common binary | 0.125 | 0.25 | 0.089 (0.005) | -0.04 | -0.29 | 0.067 (0.006) | -0.03 | -0.29 | 0.022 (0.003) | -0.01 | -0.29 | 0.249 (0.04) | 0.00 | 0.00 | 0.022 (0.003) | -0.01 | -0.29 | 0.249 (0.04) | 0.00 | 0.00 |

sTable 20: Estimated effect sizes and size of bias for simulated effect of a continuous mediator explaining the effect a continuous exposure and continuous outcome (per unit increase in exposure), and a rare binary outcome and common binary outcome on the risk difference scale using Mendelian Randomization, where simulated total effects are imprecise (Simulated N=1000)

| Outcome | True total effect | True proportion mediated | Total effect from univariate Mendelian randomisation (SD) | Size of bias (absolute) | Size of bias (relative) | Direct effect from multivariable Mendelian randomisation (SD) | Size of bias (absolute) | Size of bias (relative) | Indirect effect from multivariable Mendelian randomisation (SD) | Size of bias (absolute) | Size of bias (relative) | Proportion mediated from multivariable Mendelian randomisation (SD) | Size of bias (absolute) | Size of bias (relative) | Indirect effect from two-step Mendelian randomisation (SD) | Size of bias (absolute) | Size of bias (relative) | Proportion mediated from two-step Mendelian randomisation SD) | Size of bias (absolute) | Size of bias (relative) |
|---|---|---|---|---|---|---|---|---|---|---|---|---|---|---|---|---|---|---|---|---|
| | 0.2 | 0.05 | 0.319 (0.058) | 0.00 | 0.00 | 0.334 (0.066) | 0.00 | 0.00 | 0.2 (0.144) | 0.00 | -0.07 | -0.137 (14.419) | -0.19 | -3.73 | 0.009 (0.015) | 0.00 | -0.07 | -0.137 (14.419) | -0.19 | -3.73 |
| Continuous | 0.2 | 0.25 | 0.419 (0.076) | -0.01 | -0.03 | 0.439 (0.088) | -0.01 | -0.05 | 0.194 (0.147) | 0.00 | 0.02 | 0.228 (3.527) | -0.02 | -0.09 | 0.051 (0.03) | 0.00 | 0.02 | 0.228 (3.527) | -0.02 | -0.09 |
| | 0.2 | 0.75 | 0.648 (0.113) | 0.00 | 0.00 | 0.679 (0.131) | 0.00 | 0.03 | 0.2 (0.144) | 0.00 | -0.01 | 0.849 (9.156) | 0.10 | 0.13 | 0.148 (0.076) | 0.00 | -0.01 | 0.849 (9.156) | 0.10 | 0.13 |
| | 0.01 | 0.05 | 0.01 (0.004) | 0.00 | -0.39 | 0.352 (0.154) | 0.00 | -0.39 | 0.006 (0.009) | 0.00 | -0.38 | 0.006 (0.009) | -0.13 | -2.56 | 0 (0.001) | 0.00 | -0.38 | -0.078 (2.288) | -0.13 | -2.56 |
| Rare binary | 0.01 | 0.25 | 0.013 (0.005) | 0.00 | -0.43 | 0.456 (0.216) | 0.00 | -0.46 | 0.006 (0.01) | 0.00 | -0.36 | 0.004 (0.01) | -0.14 | -0.55 | 0.002 (0.002) | 0.00 | -0.36 | 0.113 (2.935) | -0.14 | -0.55 |
| | 0.01 | 0.75 | 0.02 (0.008) | 0.00 | -0.39 | 0.702 (0.314) | 0.00 | -0.38 | 0.006 (0.01) | 0.00 | -0.39 | 0.002 (0.011) | -1.48 | -1.97 | 0.005 (0.005) | 0.00 | -0.39 | -0.731 (42.634) | -1.48 | -1.97 |
| | 0.05 | 0.05 | 0.03 (0.008) | -0.03 | -0.62 | 0.334 (0.098) | -0.03 | -0.62 | 0.019 (0.019) | 0.00 | -0.65 | 0.018 (0.019) | 0.00 | -0.05 | 0.001 (0.002) | 0.00 | -0.65 | 0.047 (2.153) | 0.00 | -0.05 |
| Common binary | 0.05 | 0.25 | 0.04 (0.01) | -0.03 | -0.63 | 0.44 (0.124) | -0.02 | -0.64 | 0.018 (0.019) | -0.01 | -0.62 | 0.013 (0.019) | -1.68 | -6.72 | 0.005 (0.004) | -0.01 | -0.62 | -1.431 (48.68) | -1.68 | -6.72 |
| | 0.05 | 0.75 | 0.062 (0.015) | -0.03 | -0.61 | 0.682 (0.191) | -0.01 | -0.57 | 0.019 (0.019) | -0.02 | -0.62 | 0.005 (0.021) | -0.02 | -0.02 | 0.014 (0.01) | -0.02 | -0.62 | 0.731 (16.786) | -0.02 | -0.02 |

sTable 21: Estimated effect sizes and size of bias for simulated effect of a continuous mediator explaining the effect between a continuous exposure and continuous outcome using Mendelian Randomization, where true simulated total effects are small (Simulated N=5000)

| True total effect | True proportion mediated | Total effect from univariate Mendelian randomisation (SD) | Size of bias (absolute) | Size of bias (relative) | Direct effect from multivariable Mendelian randomisation (SD) | Size of bias (absolute) | Size of bias (relative) | Indirect effect from multivariable Mendelian randomisation (SD) | Size of bias (absolute) | Size of bias (relative) | Proportion mediated from multivariable Mendelian randomisation (SD) | Size of bias (absolute) | Size of bias (relative) | Indirect effect from two-step Mendelian randomisation (SD) | Size of bias (absolute) | Size of bias (relative) | Proportion mediated from two-step Mendelian randomisation SD) | Size of bias (absolute) | Size of bias (relative) |
|---|---|---|---|---|---|---|---|---|---|---|---|---|---|---|---|---|---|---|---|
| 0.01 | 0.05 | 0.01 (0.017) | 0.00 | -0.03 | 0.009 (0.014) | 0.00 | -0.02 | 0 (0.004) | 0.00 | -0.09 | -0.448 (15.516) | -0.50 | -9.97 | 0 (0.004) | 0.00 | -0.09 | -0.448 (15.516) | -0.50 | -9.97 |
| 0.05 | 0.05 | 0.05 (0.017) | 0.00 | 0.01 | 0.048 (0.014) | 0.00 | 0.01 | 0.003 (0.004) | 0.00 | 0.04 | 0.027 (0.113) | -0.02 | -0.46 | 0.003 (0.004) | 0.00 | 0.04 | 0.027 (0.113) | -0.02 | -0.46 |
| 0.1 | 0.05 | 0.1 (0.018) | 0.00 | 0.00 | 0.095 (0.015) | 0.00 | 0.00 | 0.005 (0.004) | 0.00 | -0.02 | 0.044 (0.039) | -0.01 | -0.11 | 0.005 (0.004) | 0.00 | -0.02 | 0.044 (0.039) | -0.01 | -0.11 |
| 0.01 | 0.25 | 0.01 (0.017) | 0.00 | -0.04 | 0.007 (0.014) | 0.00 | -0.05 | 0.002 (0.004) | 0.00 | -0.02 | 0.198 (5.108) | -0.05 | -0.21 | 0.002 (0.004) | 0.00 | -0.02 | 0.198 (5.108) | -0.05 | -0.21 |
| 0.05 | 0.25 | 0.05 (0.018) | 0.00 | -0.01 | 0.037 (0.014) | 0.00 | -0.01 | 0.012 (0.004) | 0.00 | -0.01 | 0.257 (0.124) | 0.01 | 0.03 | 0.012 (0.004) | 0.00 | -0.01 | 0.257 (0.124) | 0.01 | 0.03 |
| 0.1 | 0.25 | 0.099 (0.017) | 0.00 | -0.01 | 0.074 (0.014) | 0.00 | -0.01 | 0.025 (0.004) | 0.00 | -0.02 | 0.251 (0.033) | 0.00 | 0.00 | 0.025 (0.004) | 0.00 | -0.02 | 0.251 (0.033) | 0.00 | 0.00 |
| 0.01 | 0.75 | 0.01 (0.017) | 0.00 | -0.02 | 0.002 (0.014) | 0.00 | -0.04 | 0.007 (0.004) | 0.00 | -0.01 | 2.062 (68.799) | 1.31 | 1.75 | 0.007 (0.004) | 0.00 | -0.01 | 2.062 (68.799) | 1.31 | 1.75 |
| 0.05 | 0.75 | 0.051 (0.017) | 0.00 | 0.02 | 0.013 (0.014) | 0.00 | 0.06 | 0.038 (0.005) | 0.00 | 0.00 | 0.901 (1.63) | 0.15 | 0.20 | 0.038 (0.005) | 0.00 | 0.00 | 0.901 (1.63) | 0.15 | 0.20 |
| 0.1 | 0.75 | 0.1 (0.017) | 0.00 | 0.00 | 0.025 (0.015) | 0.00 | 0.00 | 0.075 (0.007) | 0.00 | 0.00 | 0.767 (0.117) | 0.02 | 0.02 | 0.075 (0.007) | 0.00 | 0.00 | 0.767 (0.117) | 0.02 | 0.02 |

sTable 22: Estimated effect sizes and size of bias for simulated effect of a phenotypically measured continuous mediator explaining the effect between a continuous exposure and continuous outcome (per unit increase in exposure), and a rare binary outcome and common binary outcome on the risk difference scale, where simulated total effects are imprecise (Simulated N=1000)

| Outcome | True total effect | True proportion mediated | Total effect (SD) | Size of bias (absolute) | Size of bias (relative) | Direct effect (SD) | Size of bias (absolute) | Size of bias (relative) | Indirect effect - difference method (SD) | Size of bias (absolute) | Size of bias (relative) | Proportion mediated - difference (SD) | Size of bias (absolute) | Size of bias (relative) | Indirect effect - product of coefficients (SD) | Size of bias (absolute) | Size of bias (relative) | Proportion mediated - product of coefficients (SD) | Size of bias (absolute) | Size of bias (relative) |
|---|---|---|---|---|---|---|---|---|---|---|---|---|---|---|---|---|---|---|---|---|
| Continuous | 0.2 | 0.05 | 0.961 (0.076) | 0.76 | 3.80 | 0.642 (0.095) | 0.45 | 2.26 | 0.961 (0.076) | 0.13 | 12.86 | 0.642 (0.095) | 0.28 | 5.67 | 0.319 (0.058) | 0.13 | 12.86 | 0.334 (0.066) | 0.28 | 0.14 |
| | 0.2 | 0.25 | 0.962 (0.079) | 0.76 | 3.81 | 0.542 (0.109) | 0.39 | 1.96 | 0.962 (0.079) | 0.27 | 5.38 | 0.542 (0.109) | 0.19 | 0.76 | 0.419 (0.076) | 0.27 | 5.38 | 0.439 (0.088) | 0.19 | 0.09 |
| | 0.2 | 0.75 | 0.961 (0.079) | 0.76 | 3.81 | 0.313 (0.137) | 0.26 | 1.31 | 0.961 (0.079) | 0.60 | 3.99 | 0.313 (0.137) | -0.07 | -0.09 | 0.648 (0.113) | 0.60 | 3.99 | 0.679 (0.131) | -0.07 | -0.04 |
| Rare binary | 0.01 | 0.05 | 0.03 (0.005) | 0.02 | 1.98 | 0.02 (0.006) | 0.01 | 1.07 | 0.03 (0.005) | 0.00 | 1.31 | 0.02 (0.006) | 0.30 | 6.04 | 0.01 (0.004) | 0.00 | 1.31 | 0.352 (0.154) | 0.30 | 6.04 |
| | 0.01 | 0.25 | 0.03 (0.005) | 0.02 | 1.97 | 0.017 (0.008) | 0.01 | 1.22 | 0.03 (0.005) | 0.01 | 2.22 | 0.017 (0.008) | 0.21 | 0.82 | 0.013 (0.005) | 0.01 | 2.22 | 0.456 (0.216) | 0.21 | 0.82 |
| | 0.01 | 0.75 | 0.03 (0.005) | 0.02 | 2.00 | 0.01 (0.01) | 0.01 | 2.88 | 0.03 (0.005) | 0.02 | 2.37 | 0.01 (0.01) | -0.05 | -0.06 | 0.02 (0.008) | 0.02 | 2.37 | 0.702 (0.314) | -0.05 | -0.06 |
| Common binary | 0.05 | 0.05 | 0.092 (0.01) | 0.04 | 0.84 | 0.062 (0.013) | 0.01 | 0.30 | 0.092 (0.01) | -0.02 | -6.86 | 0.062 (0.013) | 0.28 | 5.69 | 0.03 (0.008) | -0.02 | -6.86 | 0.334 (0.098) | 0.28 | 5.69 |
| | 0.05 | 0.25 | 0.092 (0.009) | 0.04 | 0.84 | 0.052 (0.014) | 0.01 | 0.39 | 0.092 (0.009) | 0.00 | 0.20 | 0.052 (0.014) | 0.19 | 0.76 | 0.04 (0.01) | 0.00 | 0.20 | 0.44 (0.124) | 0.19 | 0.76 |
| | 0.05 | 0.75 | 0.092 (0.01) | 0.04 | 0.84 | 0.03 (0.019) | 0.02 | 1.41 | 0.092 (0.01) | 0.05 | 1.31 | 0.03 (0.019) | -0.07 | -0.09 | 0.062 (0.015) | 0.05 | 1.31 | 0.682 (0.191) | -0.07 | -0.09 |

sTable 23: Estimated effect sizes and size of bias for simulated effect of a phenotypically measured continuous mediator explaining the effect between a continuous exposure and continuous outcome, where true total effects simulated are small (Simulated N=5000)

| True total effect | True proportion mediated | Total effect (SD) | Size of bias (absolute) | Size of bias (relative) | Direct effect (SD) | Size of bias (absolute) | Size of bias (relative) | Indirect effect - difference method (SD) | Size of bias (absolute) | Size of bias (relative) | Proportion mediated - difference (SD) | Size of bias (absolute) | Size of bias (relative) | Indirect effect - product of coefficients (SD) | Size of bias (absolute) | Size of bias (relative) | Proportion mediated - product of coefficients (SD) | Size of bias (absolute) | Size of bias (relative) |
|---|---|---|---|---|---|---|---|---|---|---|---|---|---|---|---|---|---|---|---|
| 0.01 | 0.05 | 0.61 (0.009) | 0.60 | 60.05 | 0.342 (0.007) | 0.33 | 33.25 | 0.268 (0.007) | 0.26 | 517.96 | 0.44 (0.009) | 0.39 | 7.80 | 0.268 (0.007) | 0.26 | 517.96 | 0.44 (0.009) | 0.39 | 0.19 |
| 0.05 | 0.05 | 0.65 (0.009) | 0.60 | 12.00 | 0.377 (0.007) | 0.33 | 6.58 | 0.273 (0.007) | 0.23 | 90.32 | 0.42 (0.009) | 0.37 | 7.41 | 0.273 (0.007) | 0.23 | 90.32 | 0.42 (0.009) | 0.37 | 0.19 |
| 0.1 | 0.05 | 0.7 (0.009) | 0.60 | 6.00 | 0.42 (0.007) | 0.33 | 3.25 | 0.28 (0.008) | 0.18 | 36.95 | 0.4 (0.008) | 0.35 | 6.99 | 0.28 (0.008) | 0.18 | 36.95 | 0.4 (0.008) | 0.35 | 0.17 |
| 0.01 | 0.25 | 0.61 (0.009) | 0.60 | 59.95 | 0.336 (0.006) | 0.33 | 32.87 | 0.273 (0.007) | 0.27 | 106.31 | 0.448 (0.009) | 0.20 | 0.79 | 0.273 (0.007) | 0.27 | 106.31 | 0.448 (0.009) | 0.20 | 0.10 |
| 0.05 | 0.25 | 0.65 (0.008) | 0.60 | 12.01 | 0.35 (0.007) | 0.31 | 6.26 | 0.3 (0.007) | 0.26 | 21.01 | 0.461 (0.009) | 0.21 | 0.85 | 0.3 (0.007) | 0.26 | 21.01 | 0.461 (0.009) | 0.21 | 0.11 |
| 0.1 | 0.25 | 0.7 (0.009) | 0.60 | 6.00 | 0.367 (0.007) | 0.29 | 2.92 | 0.333 (0.008) | 0.26 | 10.31 | 0.476 (0.008) | 0.23 | 0.90 | 0.333 (0.008) | 0.26 | 10.31 | 0.476 (0.008) | 0.23 | 0.11 |
| 0.01 | 0.75 | 0.61 (0.009) | 0.60 | 59.99 | 0.323 (0.007) | 0.32 | 32.07 | 0.287 (0.008) | 0.28 | 37.90 | 0.47 (0.009) | -0.28 | -0.37 | 0.287 (0.008) | 0.28 | 37.90 | 0.47 (0.009) | -0.28 | -0.14 |
| 0.05 | 0.75 | 0.65 (0.009) | 0.60 | 12.00 | 0.284 (0.007) | 0.27 | 5.42 | 0.366 (0.008) | 0.35 | 9.44 | 0.564 (0.01) | -0.19 | -0.25 | 0.366 (0.008) | 0.35 | 9.44 | 0.564 (0.01) | -0.19 | -0.09 |
| 0.1 | 0.75 | 0.7 (0.009) | 0.60 | 6.00 | 0.233 (0.008) | 0.21 | 2.08 | 0.467 (0.009) | 0.44 | 5.89 | 0.667 (0.01) | -0.08 | -0.11 | 0.467 (0.009) | 0.44 | 5.89 | 0.667 (0.01) | -0.08 | -0.04 |

sTable 24: Estimated indirect effect and proportion mediated by multiple continuous mediators explaining the association between a continuous exposure and continuous outcome in simulation analyses using phenotypic methods and MR methods (Simulated N = 5000)

| | | Total Effect (true value = 0.45) | Direct Effect (true value = 0.20) | Mutually adjusting for all mediators (Difference in coefficients/MVMR) | | | | | | | Considering each mediator independently (Product of coefficients/two-step) | | | | | | |
|---|---|---|---|---|---|---|---|---|---|---|---|---|---|---|---|---|---|
| | | | | M1 | | M2 | | M3 | | Proportion mediated combined (true value = 0.56) | M1 | | M2 | | M3 | | Proportion mediated combined (true value = 0.56 |
| | | | | Indirect effect | Proportion mediated | Indirect effect | Proportion mediated | Indirect effect | Proportion mediated | | Indirect effect | Proportion mediated | Indirect effect | Proportion mediated | Indirect effect | Proportion mediated | |
| **Phenotypic** | Independent mediators | 1.55 (0.02) | 0.26 (0.01) | 0.42 (0.01) | 0.28 (0.01) | 0.42 (0.01) | 0.30 (0.01) | 0.45 (0.01) | 0.35 (0.01) | 0.83 (0.02) | 0.93 (0.02) | 0.60 (0.01) | 1.04 (0.02) | 0.67 (0.01) | 1.21 (0.02) | 0.78 (0.01) | 2.05 |
| | Related mediators | 1.55 (0.02) | 0.26 (0.01) | 0.42 (0.01) | 0.28 (0.01) | 0.25 (0.01) | 0.16 (0.01) | 0.63 (0.02) | 0.48 (0.01) | 0.83 (0.02) | 0.94 (0.02) | 0.60 (0.01) | 1.04 (0.02) | 0.67 (0.01) | 1.37 (0.02) | 0.88 (0.01) | 2.15 |
| **MR** | Independent mediators | 0.45 (0.03) | 0.20 (0.02) | 0.05 (0.01) | 0.11 (0.02) | 0.08 (0.01) | 0.18 (0.01) | 0.12 (0.01) | 0.27 (0.02) | 0.55 (0.02) | 0.05 (0.01) | 0.11 (0.02) | 0.12 (0.01) | 0.18 (0.01) | 0.12 (0.01) | 0.27 (0.01) | 0.56 |
| | Related mediators | 0.45 (0.03) | 0.20 (0.02) | 0.05 (0.01) | 0.11 (0.02) | 0.05 (0.01) | 0.11 (0.01) | 0.15 (0.01) | 0.33 (0.02) | 0.55 (0.02) | 0.05 (0.01) | 0.11 (0.02) | 0.08 (0.01) | 0.18 (0.01) | 0.15 (0.02) | 0.33 (0.04) | 0.62 |

True indirect effect of independent mediators: M1 = 0.05; M2 = 0.08; M3 = 0.12

True indirect effect of related mediators: M1 = 0.05; M2 = 0.05; M3 = 0.12; M2 via M3; 0.03

### Supplementary Figures

#### sFigure 1: Directed acyclic graph illustrating Mendelian randomisation and the instrumental variable assumptions required for valid inference

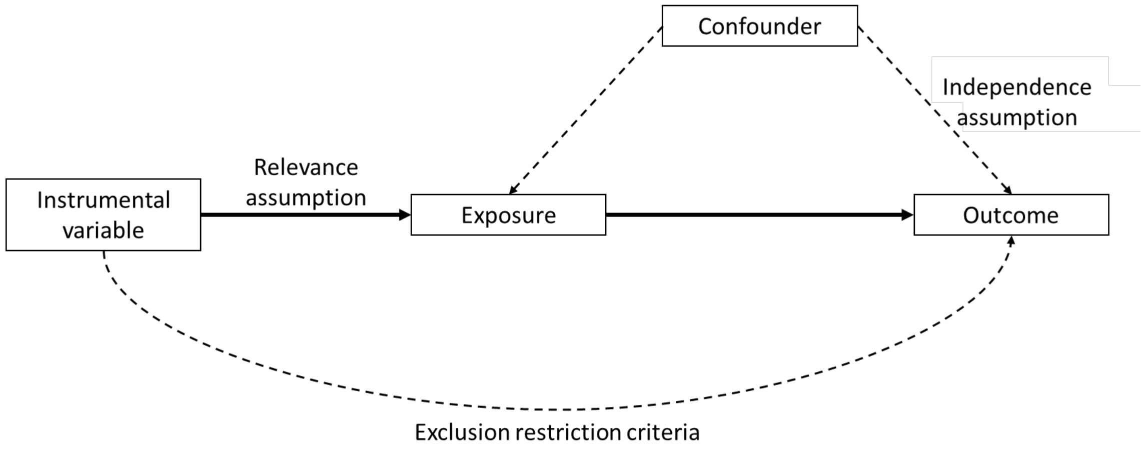

sFigure 2: Directed acyclic graphs depicting simulation scenarios considering the role of multiple mediators where in A) all three mediators are independent and in B) there is covariance between two of the three mediators

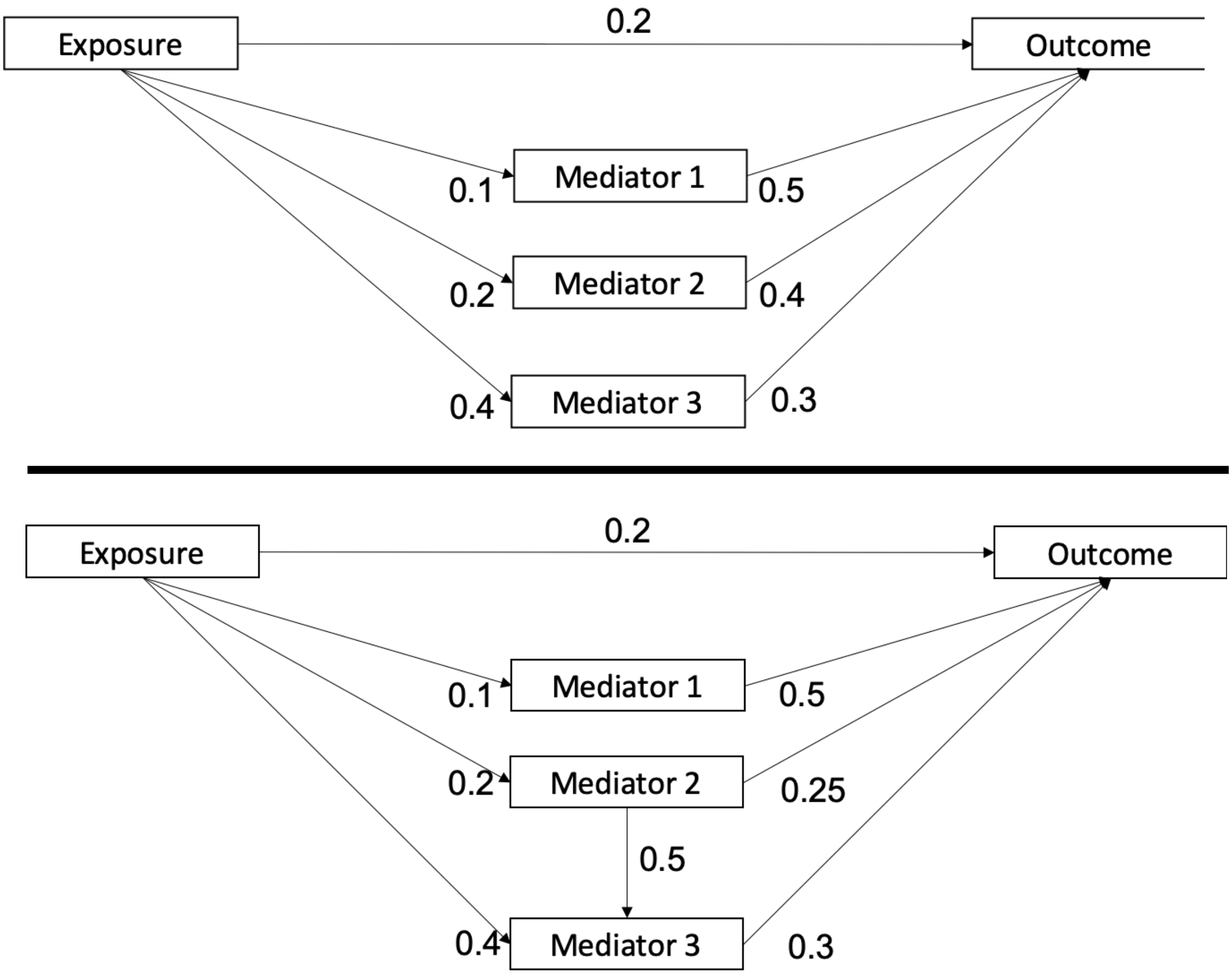

sFigure 3: Directed acyclic graphs depicting how collider bias can be introduced in phenotypic mediation analysis when conditioning on a mediator in the presence of un- or mis- measured mediator-outcome confounders

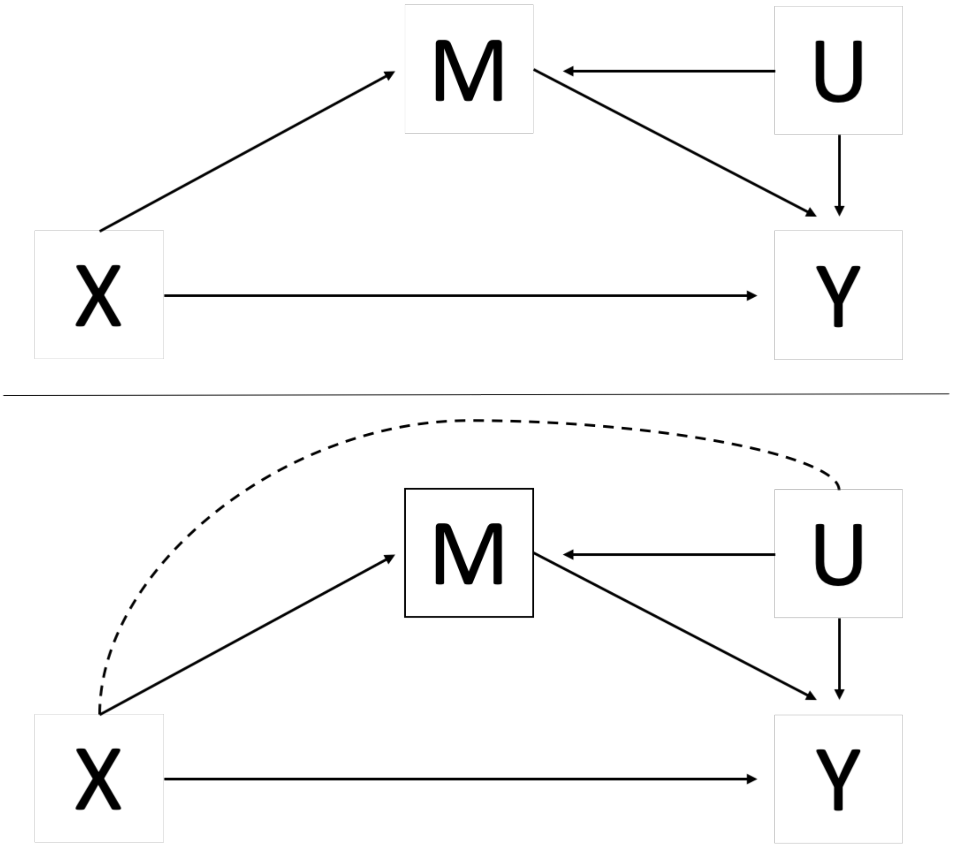

sFigure 4: Estimates of the proportion mediated and size of absolute bias when weak instrument bias is simulated in A) the exposure and B) the mediator for a true proportion mediated of 0.25 (solid line) (simulated N = 5000)

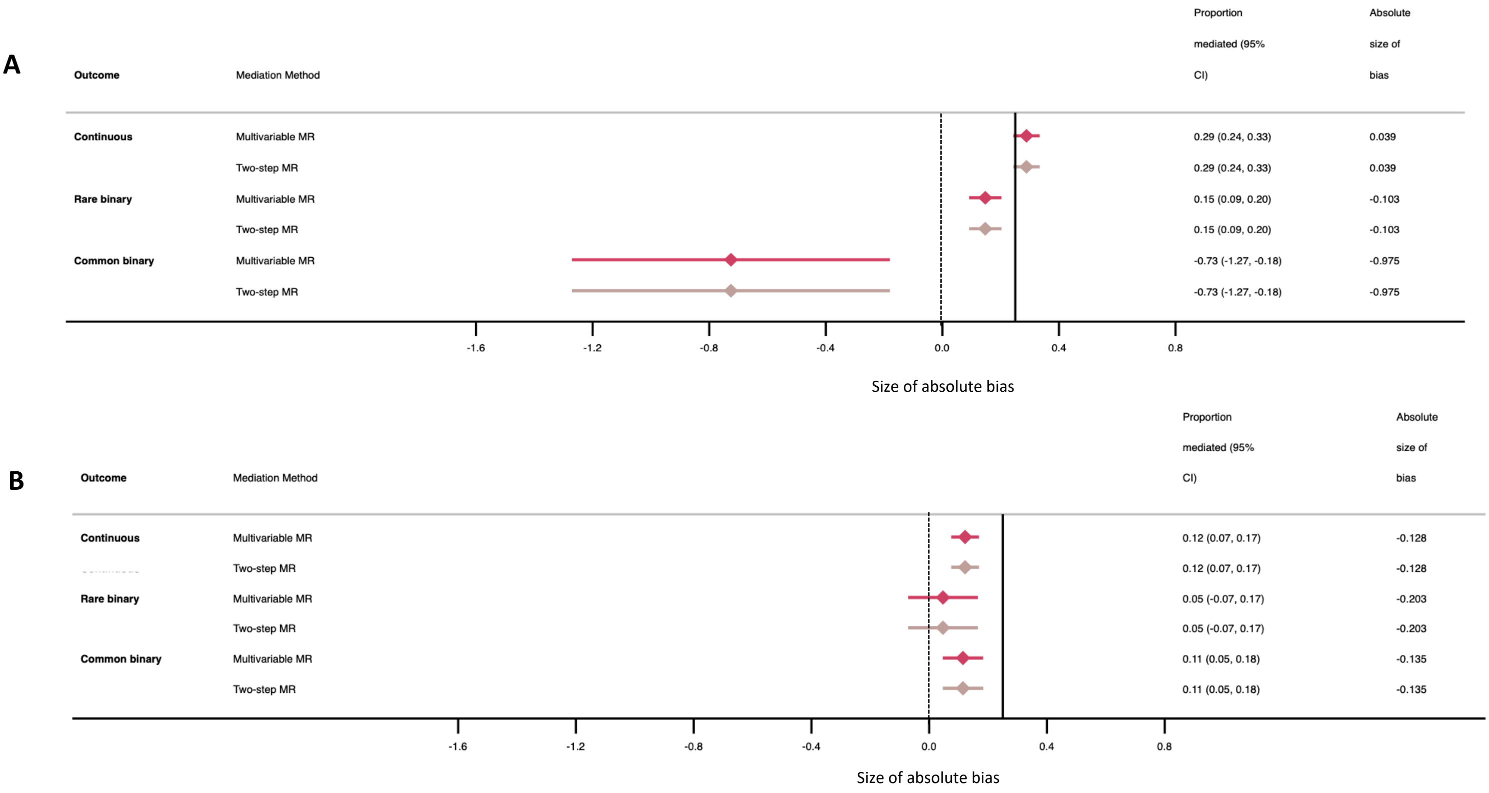
