## Supplementary material for "Mendelian randomisation for mediation analysis: current methods and challenges for implementation": Online Resource 2

### Electronic Supplementary Material 2: Applying Mendelian randomisation mediation methods to identify the role of body mass index and low-density lipoprotein cholesterol mediating the association between educational attainment and cardiovascular outcomes

Alice R Carter [1,2]*, Eleanor Sanderson [1,2], Gemma Hammerton [1,2,3], Rebecca C Richmond [1,2], George Davey Smith [1,2,4], Jon Heron [1,2,3], Amy E Taylor [1,2,4], Neil M Davies [1,2,5], Laura D Howe [1,2]

1. MRC Integrative Epidemiology Unit, University of Bristol, Bristol, UK
2. Population Health Sciences, Bristol Medical School, University of Bristol, Bristol, UK
3. Centre for Academic Mental Health, University of Bristol, Bristol, UK
4. National Institute for Health Research Biomedical Research Centre at the University Hospitals Bristol NHS Foundation Trust and the University of Bristol, Bristol, UK
5. K.G. Jebsen Center for Genetic Epidemiology, Department of Public Health and Nursing, NTNU, Norwegian University of Science and Technology, Norway.

Corresponding Author:
Alice Carter

Oakfield House,
Oakfield Grove,
Bristol,
BS8 2BN
0117 3310098
ORCiD ID: 0000-0003-2817-4195

### Methods

#### UK Biobank

At baseline, UK Biobank participants (N = 503 317) took part in questionnaires, interviews, anthropometric, physical and genetic measurements [1, 2]. We included 195 218 White British individuals, with complete data on genotypes, age, sex, educational attainment, cardiovascular outcomes, body mass index (BMI), low-density lipoprotein cholesterol (LDL-C), blood pressure (including hypertension), socioeconomic status (as measured by Townsend Deprivation Index at birth [TDI]) and place of birth. Individuals of White British descent were defined using both self-reported questionnaire data with similar genetic ancestry (principal components, PCs) to the European reference panel (from 10,000 genomes panel derived by UK Biobank) [3].

Data from the baseline assessment centre on highest qualifications completed, BMI, LDL-C, systolic blood pressure, hypertension, and all covariate measures (age, sex, place of birth and Townsend deprivation index at birth) were used for the analyses.

#### Genetic exclusion criteria

Individuals were excluded if their genetic sex differed to their gender reported at the assessment centre or for having aneuploidy of their sex chromosomes. Further individuals were excluded for being outliers for their heterozygosity and any missing genetic data. Related individuals were also excluded from analyses, and the remaining subset was a maximal set of unrelated individuals. This exclusion list was derived in-house using an algorithm applied to the list of all the related pairs provided by UK Biobank (3rd degree or closer) (sFigure 1). It preferentially removes the individuals related to the greatest number of other individuals until no related pairs remain [3].

#### Education

Participants reported their highest at the baseline assessment centre ranging from no qualifications (equivalent to leaving school after 7 years) up to degree level (equivalent to 20 years of schooling). These were converted to the International Standard Classification for Education (ISCED) coding of educational attainment (sTable 1) [4].

Mendelian randomisation studies require independent samples for the gene-exposure discovery genome-wide association study (GWAS) and analysis sample. If samples overlap MR estimates can be overestimated. Therefore, in this analysis, genetic variants were selected from GWAS that did not include UK Biobank, and as such, are not always the most recent GWAS [5, 6].

We used 72 independent single-nucleotide polymorphisms (SNPs) that attained genome-wide significance ($P<5x10^{-8}$) for education reported in main results from an earlier 2016 SSGAC GWAS meta-analysis of 293,723 individuals and were available in the UK Biobank genotyping platform, to create a weighted allele score [7]. Alleles were harmonised to all reflect education increasing SNPs and individual variants were recoded as 0, 1 or 2 according to the number of education increasing alleles. A genetic score for education was created by weighting each SNP by its relative effect size in the GWAS and summing all variants together in an additive model. Five instruments for education were not available in UK Biobank and proxy SNPs in perfect LD ($r^2$=1) were used (sTable 2).

For phenotypic sensitivity analyses to further test the non-collapsibility of odds ratios, a binary measure of low and high education was created. Individuals who left school with 10 years or less of education (equivalent to a highest qualification of GCSE, or equivalent) were classed as low education. Individuals who went into further education after high school were classed as having high education.

### Body mass index

Clinic nurses at baseline assessment centres measured participants' height (m) and weight (kg), which was used to calculate BMI ($kg/m^2$).

A polygenic score was created for BMI to use in MR analysis. We used 69 independent SNPs, available on the UK Biobank genotyping platform, which had attained genome-wide significance ($P<5x10^{-8}$) for BMI in both males and females of European ancestry in the Genetic Investigation of ANthropometric Traits (GIANT) Consortium genome-wide association study (GWAS), which did not include UK Biobank participants [8]. Alleles were harmonised to all reflect BMI increasing SNPs and individual variants were recoded as 0, 1 or 2 according to the number of BMI increasing alleles. A genetic score for BMI was created by weighting each SNP by its relative effect size in the GWAS and summing all variants together in an additive model.

In phenotypic sensitivity analyses a binary measure of BMI was created. Individuals with a BMI of less than $25kg/m^2$ were grouped together as normal or underweight individuals. Those with a BMI of $25kg/m^2$ or higher were grouped together as overweight or obese individuals.

### Low density lipoprotein cholesterol

Direct low-density lipoprotein cholesterol (LDL-C) was measured from serum samples collected at baseline, using the Enzymatic Selective Protection Method.

A polygenic score was created for use in MR analysis. We used 9 independent SNPs (sTable 3) which had attained genome-wide significance ($P<5x10^{-8}$) for LDL-C in both males and females of predominantly European ancestry from the Global Lipids Genetics consortium [9]. Genetic variants for LDL-C are often also association with high density lipoprotein cholesterol and triglycerides. To avoid bias from pleiotropy, any SNP that was association with LDL-C and at least one other lipid trait (as reported by Willer *et al*) was excluded from MR analysis ($N_{SNP}$=51). Alleles were harmonised to all reflect LDL-C increasing SNPs and individual variants were recoded as 0, 1 or 2 according to the number of LDL-C increasing alleles. The polygenic score was weighted by each SNP by its relative effect size in the GWAS and summing all variants together in an additive model.

### Blood pressure

Systolic and diastolic blood pressure were both recorded automatically and manually at the baseline assessment centre. All participants had an automatic reading, but manual readings were only taken for a subset. Each reading was taken twice, two minutes apart. This analysis uses the second reading of the automated blood pressure, where missing data were replaced with the first measure.

A binary measure of hypertension was created according to the World Health Organization's standard classification for hypertension (SBP ≥ 140 mm Hg and DBP ≥ 90 mm Hg) or if an individual was taking antihypertensive medication as recorded at the nurse's interviews.

### Cardiovascular disease

Cardiovascular disease diagnoses (including diagnoses of stroke, MI and CHD) and events were ascertained through linkage mortality data and hospital episode statistics (HES), with cases (all subtypes) defined according to ICD-9 (390-459) and ICD-10 codes (all I codes and G45) [10]. Individuals who had experienced a CVD event prior to the baseline assessment (prevalent cases) were excluded and only first event, incident cases following the assessment centre were considered. Date of diagnoses are provided by HES data, which was linked with the date of assessment centre provided by UK Biobank to identify incident and prevalent cases.

### Covariates

Variables considered as confounders were measured at the baseline assessment centres through interviews. Covariates considered were age, sex, place of birth (northing and easting co-ordinates), birth distance from London, and Townsend deprivation index at birth. Sex and ethnicity were confirmed according to genetic data. Place of Birth was adjusted for by the northing and easting birth location coordinates. Although the Townsend Deprivation Index (TDI) of historic birth locations are not recorded in UK Biobank, this has been estimated from the index of multiple deprivation indices using the current TDI of birth location as a proxy for historic birthplace TDI. Mendelian Randomisation models were also adjusted for the same confounders. Although a core assumption of MR is that the genetic variants are unrelated to confounders, there is some evidence of small associations with place of birth for the educational attainment variants in UK Biobank [11].

### Statistical Analysis

All applied analyses were carried out using Stata version 15 (StataCorp LP, Texas). The Stata package *ivreg2* was used to carry out analyses for continuous outcomes, whilst this was carried out in two separate stages for binary outcomes on the log odds or odds ratio scale.

Estimates of the total effect, direct effect, indirect effect and proportion mediated were carried out as described in the main methods. Confidence intervals for the indirect effect and proportion mediated were derived from bootstrapping with 100 replications.

Both BMI and LDL-C were considered as mediators individually and in multiple mediator models. These multiple mediator models were carried out as described in the main methods. Bidirectional univariable MR analysis was carried out to test whether mediator-mediator associations exist between BMI and LDL-C.

All phenotypic analyses were adjusted for potential confounders; age, sex, place of birth, birth distance from London, and Townsend deprivation index at birth. Mendelian Randomization analyses were adjusted for the same confounders, in addition to the 40 genetic principal components (derived by UK Biobank) to account for population structure.

All analysis code can be found at https://github.com/alicerosecarter/MediationMR

### Sensitivity analyses

In the applied phenotypic analysis, sensitivity analyses were carried out dichotomising education and/or BMI to a binary variable to further test non-collapsibility, where analyses were carried out on the log odds ratio scale. See supplementary methods for details.

Instrument strength was assessed by calculating F-statistics for univariable MR and conditional F-statistics for MVMR [12]. Sensitivity analyses for MR methods included using MR-Egger and MVMR-Egger to test for pleiotropy in the applied example [13, 14].

### Results

Descriptive characteristics of UK Biobank participants included in the real data example are shown in sTable 4.

### Effect of education on systolic blood pressure, CVD and hypertension

Both multivariable regression and univariable MR provided evidence to support a causal effect of education on systolic blood pressure, as well as for a role of BMI mediating this effect on the risk difference scale. Phenotypically, the difference method estimated the indirect effect for a one standard deviation increase in education on systolic blood pressure mediated via a one standard deviation increase in BMI to be -0.33 (-0.35 to -0.32) and the proportion mediated to be 28.4% (95% CI: 26.3% to 30.5%) (sTable 5). Using MVMR the indirect effect estimated was -0.55 (95% CI: -0.76 to -0.33). Despite the MVMR indirect effect and total effect being larger than the phenotypic difference estimate, this corresponded to a smaller proportion mediated of 16.8% (95% CI: 6.6% to 27.1%).

Using the phenotypic product of coefficients method, the indirect effect of a one standard deviation increase in education via a one standard deviation increase in BMI on systolic blood pressure was -0.33 (-0.35, -0.32) with a proportion mediated of 28.4% (95% CI: 26.5% to 30.3%) (sTable 5). Comparatively, using two-step MR, the indirect effect was estimated to be -0.55 (95% CI: -0.82 to -0.27) corresponding to a proportion mediated of 16.8% (95% CI: 6.5% to 27.2%).

Both multivariable regression and univariable MR provided evidence to support a causal effect of education on CVD, including for a mediating role of BMI. For example, the indirect effect via a one standard deviation increase in BMI on the effect of a standard deviation increase in education on incident CVD, was estimated to be $-2.45\times10^{-03}$ (95% CI for MVMR: -0.01 to $1.01\times10^{-03}$; 95% CI for two-step MR: -0.01 to $1.25\times10^{-03}$) using both MVMR and two-step MR. The estimate of the proportion mediated via both MVMR and two-step MR was 8.1% (95% CI for MVMR: -5.0% to 21.1%; 95% CI for two-step MR: -9.9% to 26.0%) .The estimates of the decomposed mediated effects were similar when analysed using the log odds ratio scale, however estimates had wider confidence intervals (sTable 5).

Mendelian randomisation suggested more education reduced risk of hypertension; however, estimates were imprecise and confidence intervals were consistent with an increased risk. This led to large confidence intervals around the estimate of the proportion mediated by BMI. On the risk difference scale, the proportion mediated that was estimated by both MVMR and two-step MR was 29.4% (95% CI for MVMR: 9.9% to 48.9%; 95% CI for two-step MR: 13.0% to 45.8%). Similar values were obtained using the log odds ratio scales (sTable 5).

For both CVD and hypertension, the decomposed mediated effects estimated on the odds ratio scale were discordant compared with those on either the risk difference or log odds ratio scale.

There was little evidence that LDL-C mediates the effect of education on systolic blood pressure, hypertension and CVD (sTable 6). Phenotypically, both the difference in coefficients method and product of coefficients method estimated 0.9% (95% CI (product method): 0.3% to 1.6%) of the effect of education on systolic blood pressure was mediated by LDL-C. In MR, both MVMR and two-step MR estimated the proportion mediated to be -1.8% (95% CI (two-step MR): -6.6 to 2.4). In both phenotypic and MR analyses, there was limited evidence that LDL-C mediated the effect of education on CVD or hypertension.

### Joint mediation by BMI and LDL-C

A one standard deviation increase in genetically instrumented BMI reduced mean LDL-C levels by -0.07 (95%CI: -0.10 to -0.04). However, a one standard deviation increase in LDL-C was not associated with BMI where the mean difference in BMI was 0.26 (94% CI: -0.38 to 0.90) (sTable 7).

Considering BMI and LDL-C jointly, in phenotypic mediation using the difference method 28.6% (95% CI: 26.4% to 30.8%) of the association between education and SBP was explained, 17.3% (95% CI: -13.6% to 21.1%) was explained when considering CVD as the outcome and 25.3% (95% CI: 23.3% to 27.3%) explained when hypertension was the outcome (both on the risk difference scale). In MR analyses, using MVMR to estimate the combined proportion mediated, 12.6% (95% CI: -2.7% to 27.9%) was explained by BMI and LDL-C on the association between education and SBP, 17.8% (95%

CI: 14.9% to 21.4%) was explained when hypertension was the outcome and 24.5% (95% CI: 22.0% to 27.1%) was explained when CVD was the outcome (both on the risk difference scale) (sTable 8).

#### Sensitivity analyses

Applied examples using phenotypic mediation methods were extended to examine the role of binary exposures or mediators on non-collapsibility. In both rare and common binary outcomes, where the education exposure was dichotomized to low (10 years of education or less) compared with high education (greater than 10 years of education) the difference in coefficients method and product of coefficients method estimated similar mediating roles by a continuous standard deviation increase in BMI. For example, the proportion mediated by BMI on the association between education (high vs low) and CVD was 15.8% (13.5% to 18.2%) for the difference method and 15.6% (15.1% to 16.2%) for the product of coefficients method. Where the mediator was binary (normal and underweight vs overweight and obese) the two methods diverged. For example, the proportion mediated by high versus low BMI on the association between a standard deviation increase in education and incident CVD was 8.6% (7.3% to 10.2%) for the difference in coefficients method and 45.3% (38.8% to 51.9%) for the product of coefficients method. This was similar when both the exposure and outcome were considered as binary. Similar results were also seen when considering common hypertension as the outcome (sTable 9). Where both the mediator and outcome are binary, counterfactual methods for mediation analysis should be considered.

All instruments had strong F statistics and conditional F statistics (sTable 10).

Both MR-Egger and MVMR-Egger provide little evidence to support pleiotropic effects of the instruments biasing results (sTable 11).

### Results in context

Our applied example demonstrates a causal total effect of education on systolic blood pressure, supporting results in the wider literature [15-17] and shows that BMI is a mediator of the association between education and systolic blood pressure. Given our previous analyses showing that systolic blood pressure is itself a mediator of the associations between education and CVD, this work suggests systolic blood pressure is downstream of BMI on the causal pathway, although we have not explored bi-directional association in this analysis.

Despite LDL-C being a major, modifiable risk factor for cardiovascular outcomes [18, 19], and there being some evidence from non-genetic instrumental variable analyses that levels of LDL-C decrease with increased education [20], it does not appear to explain any of the of educational inequalities on these outcomes. Although the instrument for LDL-C only comprised of 9 SNPs which did not have pleiotropic effects on either high-density lipoprotein cholesterol or triglycerides, the F-statistics and conditional F-statistics for these analyses remained high, and results are unlikely to be biased due to weak instrument bias.

Considering BMI and LDL-C jointly, increased the proportion mediated between education and CVD increased by 9% compared with the BMI individually. However, for each of the individual estimates of the proportion mediated and the joint estimate the confidence intervals were wide. Considering systolic blood pressure and hypertension as the outcomes, the proportion mediated decreased.

sTable 1: International Standard for Classification of Education codes mapped to UK Biobank self-report highest qualification to estimate years of education

| Qualification (As reported in UK Biobank) | ISCED | Years of education |
|---|---|---|
| **College or University degree** | 5 | 20 |
| **NVQ or HND or HNC or equivalent** | 5 | 19 |
| **Other prof. qual. eg: nursing, teaching** | 4 | 15 |
| **A levels/AS levels or equivalent** | 3 | 13 |
| **O levels/GCSEs or equivalent** | 2 | 10 |
| **CSEs or equivalent** | 2 | 10 |
| **None of the above** | 1 | 7 |
| **Prefer not to answer** | Excluded | |

sTable 2: Proxy SNPs for education instrument used in one-sample MR analysis

| GWAS SNP (Okbay) | SNP in LD used (UKBB) |
|---|---|
| rs114598875 | rs17538393 |
| rs148734725 | rs9878943 |
| rs9320913 | rs1487445 |
| rs8005528 | rs8008779 |
| rs192818565 | rs55943044 |

sTable 3: Independent SNPs used as instruments for LDL-C

| SNP | Effect allele | Other allele | Beta | P value |
|---|---|---|---|---|
| rs1801689 | C | A | 0.1028 | 9.81E-12 |
| rs2328223 | C | A | 0.0299 | 5.63E-09 |
| rs364585 | G | A | 0.0249 | 4.28E-10 |
| rs1250229 | C | T | 0.0243 | 3.13E-08 |
| rs8017377 | A | G | 0.0303 | 2.52E-15 |
| rs267733 | G | A | -0.0331 | 5.29E-09 |
| rs4942486 | C | T | -0.0243 | 2.26E-11 |
| rs5763662 | T | C | 0.0767 | 1.19E-08 |
| rs2710642 | A | G | 0.0239 | 6.09E-09 |

sTable 4: UK Biobank cohort descriptive statistics

| Variable | | All | Female | Male |
|---|---|---|---|---|
| | | Mean (SD) or N (%) | Mean (SD) or N (%) | Mean (SD) or N (%) |
| Sex | | | 118, 822 (55%) | 97, 537 (45%) |
| Age | | 56.53 (8.03) | 56.47 (7.91) | 56.59 (8.17) |
| Educational attainment (years) | 7 | 34 432 (16%) | 19 451 (16%) | 14 982 (15%) |
| | 10 | 38 245 (18%) | 24 633 (21%) | 13 612 (14%) |
| | 13 | 11 832 (5%) | 6 929 (6%) | 4 903 (5%) |
| | 15 | 26 748 (12%) | 16 196 (14%) | 10 552 (11%) |
| | 19 | 34 860 (16%) | 15 016 (13%) | 19 844 (20%) |
| | 20 | 70 242 (32%) | 36 597 (31%) | 33 645 (35%) |
| Body mass index | | 27.26 (4.69) | 26.91 (5.06) | 27.68 (4.15) |
| Low-density lipoprotein cholesterol | | 3.61 (0.85) | 3.66 (0.86) | 3.55 (0.84) |
| Systolic blood pressure | | 136.50 (18.68) | 133.76 (19.25) | 139.83 (17.38) |
| Incident cardiovascular disease | Control | 200 284 (93%) | 111 962 (94%) | 88 322 (91%) |
| | Case | 16 074 (7%) | 6 859 (6%) | 9 215 (9%) |
| Hypertension | Control | 159 719 (74%) | 90 219 (76%) | 69 500 (71%) |
| | Case | 56 640 (25%) | 28 603 (24%) | 28 037 (29%) |

sTable 5: Real-data example estimating the mediating role of BMI independently between education and systolic blood pressure, cardiovascular disease and hypertension, using multivariable observational methods and mendelian randomisation methods

| | | **Multivariable regression** | | | | | | **Mendelian Randomisation** | | | | | |
|---|---|---|---|---|---|---|---|---|---|---|---|---|---|
| **Outcome** | **Scale** | **Total effect (95% CI)** | **Direct effect (95% CI)** | **Indirect effect - difference (95% CI)** | **% mediated - difference (95% CI)** | **Indirect effect - product (95% CI)** | **% mediated - product (95% CI)** | **Total effect 95% CI)** | **Direct effect (95% CI)** | **Indirect effect - MVMR (95% CI)** | **% mediated - MVMR (95% CI)** | **Indirect effect - two-step MR (95% CI)** | **% mediated - two-step MR 95% CI)** |
| **Systolic blood pressure** | Mean difference | -1.17 (-1.24, -1.09) | -0.84 (-0.91, -0.76) | -0.33 (-0.35, -0.32) | 28.40 (26.26, 30.54) | -0.33 (-0.35, -0.32) | 28.40 (26.54, 30.27) | -3.24 (-4.13, -2.35) | -2.69 (-3.63, -1.76) | -0.55 (-0.76, -0.33) | 16.84 (6.59, 27.09) | -0.55 (-0.82, -0.27) | 16.84 (6.50, 27.18) |
| **Cardiovascular disease** | Risk difference | -0.01 (-0.01, -0.01) | -0.01 (-0.01, -4.61x10$^{-03}$) | -1.21x10$^{-03}$ (-1.35x10$^{-03}$, -1.08x10$^{-03}$) | 17.37 (14.02, 20.73) | -1.21x10$^{-03}$ (-1.36x10$^{-03}$, -1.07x10$^{-03}$) | 17.37 (14.12, 20.62) | -0.03 (-0.04, -0.02) | -0.03 (-0.04, -0.01) | -2.45x10$^{-03}$ (-0.01, 1.01x10$^{-03}$) | 8.05 (-5.00, 21.10) | -2.45x10$^{-03}$ (-0.01, 1.25x10$^{-03}$) | 8.05 (-9.94, 26.04) |
| | Log odds ratio | -0.10 (-0.12, -0.08) | -0.08 (-0.10, -0.07) | -0.02 (-0.02, -0.02) | 17.72 (14.12, 21.31) | -0.02 (-0.02, -0.02) | 18.26 (14.34, 22.19) | -0.45 (-0.65, -0.25) | -0.41 (-0.62, -0.2) | -0.04 (-0.09, 0.02) | 8.27 (-4.24, 20.78) | -0.04 (-0.10, 0.02) | 8.29 (-3.68, 20.25) |
| | Odds ratio | 0.90 (0.89, 0.92) | -0.40 (-0.61, -0.18) | -0.02 (-0.02, -0.01) | -1.81 (-2.02, -1.61) | -0.13 (-0.13, -0.12) | -14.30 (-15.01, -13.59) | 0.64 (0.52, 0.78) | 0.66 (0.54, 0.82) | -0.02 (-0.06, 0.01) | -3.78 (-8.90, 1.33) | -0.43 (-0.51, -0.35) | -67.28 (-84.73, -49.83) |
| **Hypertension** | Risk difference | -0.03 (-0.03, -0.03) | -0.02 (-0.02, -0.02) | -0.01 (-0.01, -0.01) | 25.58 (23.31, 27.85) | -0.01 (-0.01, -0.01) | 25.58 (23.25, 27.91) | -0.06 (-0.08, -0.04) | -0.04 (-0.07, -0.02) | -0.02 (-0.02, -0.01) | 29.36 (9.87, 48.85) | -0.02 (-0.03, -0.01) | 29.36 (12.95, 45.77) |
| | Log odds ratio | -0.13 (-0.14, -0.12) | -0.1 (-0.11, -0.09) | -0.03 (-0.03, -0.03) | 24.94 (22.83, 27.06) | -0.04 (-0.04, -0.03) | 27.43 (25.02, 29.83) | -0.30 (-0.41, -0.18) | -0.21 (-0.33, -0.09) | -0.09 (-0.11, -0.06) | 29.44 (11.38, 47.5) | -0.09 (-0.11, -0.06) | 29.56 (9.97, 49.15) |
| | Odds ratio | 0.88 (0.87, 0.89) | -0.22 (-0.34, -0.10) | -0.03 (-0.03, -0.03) | -3.27 (-3.45, -3.09) | -0.15 (-0.16, -0.14) | -17.15 (-17.88, -16.41) | 0.74 (0.66, 0.83) | 0.81 (0.72, 0.92) | -0.07 (-0.09, -0.04) | -9.10 (-12.49, -5.71) | -0.49 (-0.57, -0.40) | -65.63 (-77.37, -53.89) |

sTable 6: Real-data example estimating the mediating role of low-density lipoprotein cholesterol independently between education and systolic blood pressure, cardiovascular disease and hypertension, using multivariable observational methods and mendelian randomisation methods

| Outcome | Scale | Multivariable regression | | | | | | Mendelian Randomisation | | | | | |
|---|---|---|---|---|---|---|---|---|---|---|---|---|---|
| | | **Total effect (95% CI)** | **Direct effect (95% CI)** | **Indirect effect - difference (95% CI)** | **% mediated - difference (95% CI)** | **Indirect effect - product (95% CI)** | **% mediated - product (95% CI)** | **Total effect 95% CI)** | **Direct effect (95% CI)** | **Indirect effect - MVMR (95% CI)** | **% mediated - MVMR (95% CI)** | **Indirect effect - two-step MR (95% CI)** | **% mediated - two-step MR 95% CI)** |
| **Systolic blood pressure** | Mean difference | -1.17 (-1.24, -1.09) | -1.16 (-1.23, -1.08) | -0.01 (-0.02, -4.11x10$^{-03}$) | 0.94 (0.26, 1.74) | -0.01 (-0.02, -3.56x10$^{-03}$) | 0.94 (0.30, 1.58) | -3.24 (-4.13, -2.35) | -3.30 (-4.21, -2.39) | 0.06 (-0.09, 0.21) | -1.83 (-6.30, 2.64) | 0.06 (-0.09, 0.21) | -1.83 (-6.56, 2.89) |
| **Cardiovascular disease** | Risk difference | -0.01 (-0.01, -0.01) | -0.01 (-0.01, -0.01) | 1.68x10$^{-05}$ (2.48x10$^{-06}$, 3.10x10$^{-05}$) | -0.24 (-0.45, -0.03) | 1.68x10$^{-05}$ (1.06x10$^{-06}$, 2.29x10$^{-05}$) | -0.24 (-0.45, -0.03) | -0.03 (-0.04, -0.02) | -0.03 (-0.04, -0.02) | -5.29x10$^{-04}$ (-2.00x10$^{-03}$, 9.43x10$^{-04}$) | 1.74 (-3.91, 7.38) | -5.29x10$^{-04}$ (-2.01x10$^{-03}$, 9.52x10$^{-04}$) | 1.74 (-4.72, 8.19) |
| | Log odds ratio | -0.10 (-0.12, -0.08) | -0.10 (-0.12, -0.08) | -8.63x10$^{-05}$ (-2.12x10$^{-04}$, -3.92x10$^{-05}$) | 0.09 (-0.07, 0.24) | 9.72x10$^{-05}$ (-4.58x10$^{-04}$, 2.40x10$^{-04}$) | -0.10 (-0.26, 0.07) | -0.45 (-0.65, -0.25) | -0.44 (-0.64, -0.24) | -0.01 (-0.02, 0.01) | 1.8 (-2.95, 6.55) | -0.01 (-0.03, 0.02) | 1.80 (-3.17, 6.77) |
| | Odds ratio | 0.90 (0.89, 0.92) | -0.10 (0.89, 0.92) | -7.80x10$^{-05}$ (-2.07x10$^{-04,}$ 5.12x10$^{-05}$) | -0.01 (-0.02, 4.26x10$^{-03}$) | -0.01 (-0.01, -1.55x10$^{-03}$) | -0.69 (-1.20, -0.20) | 0.64 (0.52, 0.78) | 0.64 (0.53, 0.78) | -0.04 (-0.11, 0.04) | -0.81 (-2.26, 0.64) | -0.04 (-0.11, 0.04) | -5.73 (-19.30, 7.88) |
| **Hypertension** | Risk difference | -0.03 (-0.03, -0.03) | -0.03 (-0.03, -0.03) | 3.07x10-04 (1.02x10$^{-04}$, 5.13x10$^{-04}$) | -1.12 (-1.86, -0.38) | 3.07x10$^{-04}$ (1.08x10$^{-04}$, 5.06x10$^{-04}$) | -1.12 (-1.99, -0.25) | -0.06 (-0.08, -0.04) | -0.06 (-0.08, -0.04) | 8.27x10$^{-04}$ (-2.11x10$^{-03}$ 3.76x10$^{-03}$) | -1.37 (-6.43, 3.68) | 8.27x10-04 (-1.68x10$^{-03}$, 3.33x10$^{-03}$) | -1.37 (-5.64, 2.90) |
| | Log odds ratio | -0.13 (-0.14, -0.12) | -0.13 (-0.14, -0.12) | 1.53x10$^{-03}$ (5.98x10$^{-03}$, 2.46x10$^{-03}$) | -1.18 (-1.98, -0.39) | 1.43x10$^{-03}$ (4.03x10$^{-04}$, 2.46x10$^{-03}$) | -1.11 (-1.88, -0.35) | -0.30 (-0.41, -0.18) | -0.30 (-0.41, -0.19) | 4.13x10-03 (-4.23x10$^{-03}$, 0.01) | -1.40 (-4.28, 1.49) | 4.12x10$^{-03}$ (-6.86x10$^{-03}$, 0.02) | -1.39 (-6.70, 3.91) |
| | Odds ratio | 0.88 (0.87, 0.89) | -0.13 (0.87, 0.89) | 1.34x10$^{-03}$ (4.74x10$^{-04}$, 2.21x10$^{-03}$) | 0.15 (0.04, 0.26) | -0.01 (-0.01, -1.53x10$^{-03}$) | -0.57 (-0.96, -0.19) | 0.74 (0.66, 0.83) | 0.74 (0.66, 0.83) | -0.02 (-0.06, 0.01) | 0.41 (-0.46, 1.28) | -0.02 (-0.06, 0.01) | -3.14 (-9.16, 2.89) |

sTable 7: Effect of a one standard deviation increase of body mass index (BMI) on low-density lipoprotein cholesterol (LDL-C) and a one standard deviation increase in LDL-C on BMI in a Mendelian randomisation analysis

| Exposure | Outcome | Beta (95% CI) |
|---|---|---|
| BMI | LDL-C | -0.07 (-0.10, -0.04) |
| LDL-C | BMI | 0.26 (-0.38, 0.90) |

sTable 8: Real-data example estimating the joint mediating role of BMI and LDL-C between education and systolic blood pressure (SBP), hypertension and cardiovascular disease (CVD) using multivariable observational methods and mendelian randomisation methods, where the joint direct effect was estimated using the difference in coefficients method, or multivariable mendelian randomisation method

| Outcome | Scale | Multivariable regression | | | Mendelian Randomisation | | |
|---|---|---|---|---|---|---|---|
| | | Total effect (95% CI) | Direct effect (95% CI) | % mediated - difference (95% CI) | Total effect 95% CI) | Direct effect (95% CI) | % mediated - MVMR (95% CI) |
| Systolic blood pressure | Mean difference | -1.17(-1.25, -1.09) | -0.83 (-0.91, -0.76) | 28.59 (26.36, 30.83) | -3.241 (-4.14, -2.35) | -2.83 (-3.79, -1.87) | 12.58 (-2.72, 27.89) |
| Cardiovascular disease | Risk difference | -0.01 (-0.01, -0.01) | -0.01 (-0.01, -0.01) | 17.31 (13.55, 21.07) | -0.10 (-0.12, -0.08) | -0.08 (-0.10, -0.07) | 17.80 (14.19, 21.42) |
| | Log odds ratio | -0.03 (-0.04, -0.02) | -0.03 (-0.04, -0.01) | 11.68 (-3.12, 26.49) | -0.45 (-0.65, -0.25) | -0.40 (-0.61, -0.18) | 12.03 (-4.29, 28.36) |
| Hypertension | Risk difference | -0.03 (-0.03, -0.03) | -0.02 (-0.02, -0.02) | 25.32 (23.32, 27.32) | -0.13 (-0.14, -0.12) | -0.10 (-0.11, -0.09) | 24.52 (21.99, 27.05) |
| | Log odds ratio | -0.06 (-0.08, -0.04) | -0.05 (-0.07, -0.02) | 11.68 (-10.87, 34.24) | -0.30 (-0.41, -0.18) | -0.22 (-0.34, -0.10) | -33.60 (-148.64, 81.45) |

sTable 9: Evaluating non-collapsibility in real-data example with binary exposures and/or binary mediators with a rare binary and common binary outcome on the log odds ratio scale

| Outcome | Exposure (education) | Mediator (BMI) | Total effect | Indirect effect (difference method) | Proportion mediation (difference method) | Indirect effect (product method) | Proportion mediation (product method) |
|---|---|---|---|---|---|---|---|
| **CVD (rare)** | Education low vs high | Continuous BMI | -0.18 (-0.21, -0.14) | -0.03 (-0.03, -0.03) | 17.70 (13.53, 21.88) | -0.03 (-0.04, -0.03) | 18.29 (14.23, 22.34) |
| | Continuous education | Normal / underweight vs obese/overweight | -0.10 (-0.12, -0.08) | -0.01 (-0.01, -0.01) | 9.14 (6.93, 11.36) | -0.05(-0.06, -0.04) | 47.75 (36.05, 59.45) |
| | Education low vs high | Normal / underweight vs obese/overweight | -0.18 (-0.21, -0.14) | -0.02 (-0.02, -0.01) | 9.38 (6.83, 11.93) | -0.09 (-0.10, -0.07) | 48.46 (34.74, 62.17) |
| **Hypertension (common)** | Education low vs high | Continuous BMI | -0.23 (-0.25, -0.21) | -0.06 (-0.06, -0.05) | 24.62 (22.14, 27.10) | -0.06 (-0.07, -0.06) | 27.14 (24.63, 29.66) |
| | Continuous education | Normal / underweight vs obese/overweight | -0.13 (-0.14, -0.12) | -0.02 (-0.02, -0.02) | 14.52 (12.95, 16.11) | -0.10 (-0.05, -0.04) | 75.72 (67.95, 83.48) |
| | Education low vs high | Normal / underweight vs obese/overweight | -0.23 (-0.25, -0.21) | -0.03 (-0.04, -0.03) | 14.58 (12.88, 16.30) | -0.17 (-0.18, -0.16) | 75.74 (68.22, 83.25) |

Low education defined as 10 years or less years of education, equivalent to a highest qualification of GCSE/CSE or equivalent. High education defined as more than 10 years of education equivalent to post-secondary qualifications. Low education had a prevalence of 34%

BMI = body mass index

Normal or underweight defined as a BMI below 25 Kg/m$^2$. Overweight or obese defined as a BMI of 25Kg/m$^2$ of above. Normal weight or underweight had a prevalence of 34%.

sTable 10: F statistics to test instrument strength in real-data Mendelian randomization

| | **Education** | **BMI** | **LDL-C** |
|---|---|---|---|
| **Conditional variable** | | | |
| **Education** | **1571.22** | 2707.52 | 213.97 |
| **BMI** | 1374.21 | **3416.36** | 210.80 |
| **LDL-C** | 1555.39 | 3454.31 | **214.19** |
| **Education, BMI and LDL-C** | 1366.27 | 1326.44 | 216.34 |

F statistics for univariable MR analyses are in bold, all other estimates are conditional F statistics

BMI = Body mass index  LDL-C = low-density lipoprotein cholesterol sTable 11: MR-Egger and MVMR-Egger results for the applied example examining the mediating role of body mass index (BMI) and low-density lipoprotein cholesterol (LDL-C) on the association between education and systolic blood pressure, cardiovascular disease and hypertension, estimated on the mean or risk difference scale

| | **Systolic blood pressure** | | **Cardiovascular disease** | | **Hypertension** | |
|---|---|---|---|---|---|---|
| | Univariable MR-Egger | Multivariable MR-Egger* | Univariable MR-Egger | Multivariable MR-Egger* | Univariable MR-Egger | Multivariable MR-Egger* |
| **Education** | | | | | | |
| Constant | $-2.15x10^{-6}$ | | $-5.07x10^{-10}$ | | $4.52x10^{-9}$ | |
| 95% confidence interval | $-2.64x10^{-5}$ to $2.21x10^{-5}$ | | $-1.02x10^{-7}$to $1.01x10^{-7}$ | | $-4.27x10^{-7}$ to $4.36x10^{-7}$ | |
| P Value | 0.861 | | 0.922 | | 0.983 | |
| **BMI** | | | | | | |
| Constant | $-1.71x10^{-6}$ | $-1.57x10^{-6}$ | $-4.02x10^{-9}$ | $-3.98x10^{-9}$ | $-1.60x10^{-8}$ | $-1.72x10^{-8}$ |
| 95% confidence interval | $-2.72x10^{-5}$ to $2.38x10^{-5}$ | $-1.94x10^{-5}$ to $1.63x10^{-5}$ | $-9.15x10^{-8}$ to $9.95x10^{-8}$ | $-6.91x10^{-8}$ to $-7.71x10^{-8}$ | $-4.90x10^{-7}$ to $5.22x10^{-7}$ | $-3.1x10^{-7}$ to $3.44x10^{-7}$ |
| P Value | 0.894 | 0.863 | 0.934 | 0.915 | 0.996 | 0.918 |
| **LDL-C** | | | | | | |
| Constant | $-2.45x10^{-6}$ | $-2.14x10^{-6}$ | $-1.44x10^{-9}$ | $-5.13x10^{-10}$ | $-1.70x10^{-9}$ | $-4.04x10^{-9}$ |
| 95% confidence interval | $-4.35x10^{-5}$ to $3.86x10^{-5}$ | $-2.53x10^{-5}$ to $2.11x10^{-5}$ | $-1.54x10^{-7}$ to $1.51x10^{-7}$ | $-9.77x10^{-8}$ to $9.67x10^{-8}$ | $-6.38x10^{-7}$ to $6.35x10^{-7}$ | $-3.79x10^{-7}$ to $3.87x10^{-7}$ |
| P Value | 0.905 | 0.855 | 0.985 | 0.922 | 0.996 | 0.983 |

*Education adjusted for either BMI or LDL-C

sFigure 1: Flow chart for exclusions made in UK Biobank for resultant sample for mediation analysis

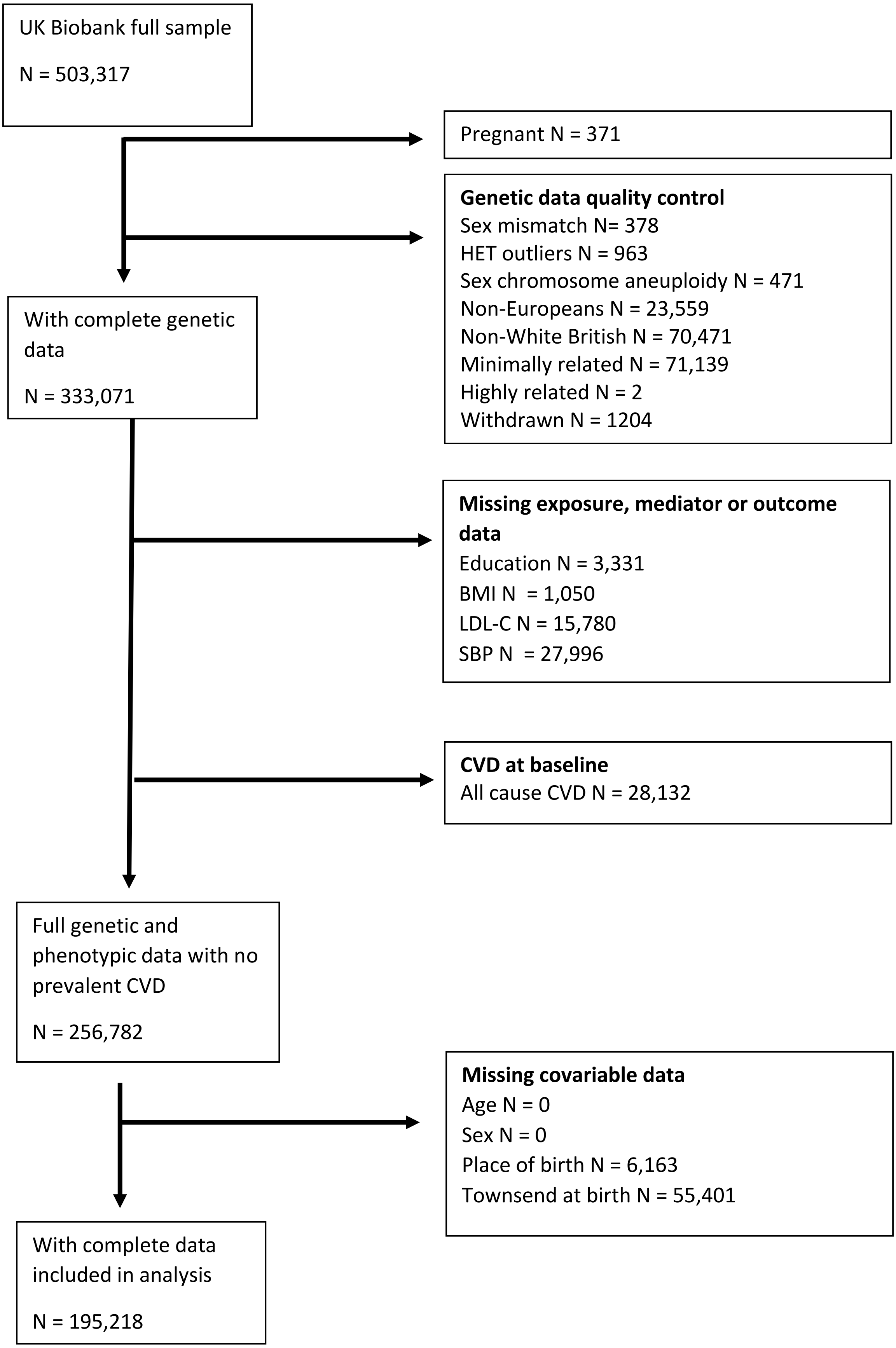
